# Fatty acid elongases 1-3 have distinct roles in mitochondrial function, growth and lipid homeostasis in *Trypanosoma cruzi*

**DOI:** 10.1101/2022.11.22.517509

**Authors:** Lucas Pagura, Peter C. Dumoulin, Cameron C. Ellis, Igor L. Estevao, Maria T. Mendes, Igor C. Almeida, Barbara A. Burleigh

## Abstract

Trypanosomatids are a diverse group of uniflagellate protozoa that include globally important pathogens such as *Trypanosoma cruzi*, the causative agent of Chagas disease. Trypanosomes lack the fatty acid synthase (FAS)-I system typically used for *de novo* synthesis of long chain fatty acids (LCFA) in other eukaryotes. Instead, these microbes have evolved a modular fatty acid elongase (ELO) system comprised of individual ELO enzymes that operate processively. The role of the ELO system in maintaining lipid homeostasis in trypanosomes has not been determined. Here we demonstrate that ELO2 and ELO3 are required for global lipidome maintenance in the insect stage of *T. cruzi* whereas ELO1 is dispensable for this function. Instead, ELO1 activity is needed to sustain mitochondrial activity and normal growth. The cross-talk between microsomal ELO1 and the mitochondrion is a novel finding that merits examination of the trypanosomatid ELO pathway as critical for central metabolism.

## Introduction

The fatty acid (FA) composition of lipid bilayers can profoundly influence the biophysical properties (fluidity, curvature, microdomain organization) and biological functions (secretion, signal transduction, adaptation to stress) of cellular membranes ^1–3^. As such, regulation of the fatty acid pools used for the synthesis and maintenance of biological membranes is a critical homeostatic function of all cells ^4^. While these regulatory processes have been well-studied in mammals and yeast ^5–7^, the mechanisms governing fatty acid and lipid homeostasis in pathogenic protozoa are poorly understood.

*Trypanosoma cruzi* is a member of a diverse group of flagellated protozoa that include parasitic organisms responsible for globally important human diseases such as leishmaniasis (*Leishmania spp*.), sleeping sickness (*Trypanosoma brucei*) and Chagas disease (*Trypanosoma cruzi*). Throughout its complex life cycle, *T. cruzi* colonizes hematophagous triatomine insects and mammalian hosts and switches between actively dividing and non-dividing forms in each setting. Survival in these disparate environments requires both morphological and metabolic adaptation ^8^. A critical feature of adaptation in *T. cruzi* is the ability to alter membrane lipid composition and to alter the fatty acyl moieties of the abundant glycosylphosphatidylinositols (GPIs) that anchor arrays of glycoproteins and glycolipids to the parasite surface ^9,10^. The biological consequences of fatty acid remodeling during development is best documented for the class of surface GPI-anchored mucins, where changes in FA chain length and/or degree of saturation can activate or dampen the pro-inflammatory activity of these molecules ^11^. It is currently unknown how *T. cruzi* regulates the composition or abundance of fatty acids needed for lipid synthesis ^12^, but similar to other cell types, this protozoan has the capacity to synthesize LCFA endogenously ^10,13^ and to take up fatty acids from the environment ^14^.

Bulk synthesis of LCFA in most eukaryotes, is achieved using a cytosolic type-I fatty acid synthase (FAS-I) ^15^ that is absent in trypanosomatids. A separate mitochondrial FAS-II system also exists, which has a more restricted role in cells, as a nutrient-sensitive coordinator of mitochondrial oxidative phosphorylation ^16^. Short chain fatty acid (SCFA) intermediates generated by FAS-II can also be diverted to synthesize other molecules such as lipoic acid, a critical cofactor needed for the stabilization and activity of a subset of mitochondrial enzymes ^17,18^. Similar to other eukaryotes, trypanosomes also have a mitochondrial FAS-II ^12,13^. In addition, trypanosomatids have develop a specialized microsomal fatty acid elongase (ELO) system for the synthesis of bulk LCFA ^10,13^.

First described in *T. brucei* ^13^, the trypanosomatid ELO system consists of individual fatty acid elongase enzymes (ELOs 1-5 in *T. cruzi* ^19^) that exhibit distinct specificities for fatty acyl-CoA substrates based on carbon chain length and/or degree of unsaturation ^13^. ELOs 1-4 are numbered according to their relative position in a sequential pathway (ELO1, ELO2, etc.), where the product of one ELO becomes the substrate for the next ^13^. Studies using isolated membranes indicate that the ELO enzymes can also function independently to elongate exogenous fatty acyl substrates of the appropriate chain length ^13^. ELOs utilize malonyl-CoA as a two-carbon donor to extend fatty acyl-CoA substrates in a four-step cycle involving: (1) a β-ketoacyl-CoA synthase (the ‘ELO’) responsible for condensing a pre-existing fatty acyl-CoA with malonyl-CoA; this is the first and rate-limiting step in each elongation cycle and guides substrate specificity; (2) a β-ketoacyl-CoA reductase (KCR); (3) β-hydroxyacyl-CoA dehydrase (DEH); and (4) a trans-2-enoyl-CoA reductase (EnCR) ^10,13^. The ELO1 cycle elongates short-chain fatty acyl-CoA substrates (C4, C6, and C8) to generate C10, but unlike FAS-I, ELO1 cannot initiate fatty acid synthesis using acetyl-CoA, instead it use butyryl-CoA as the main substrate ^13^. ELO2 extends C10 and C12 substrates to generate C14, and ELO3 elongates C14 to generate C16 and C18 fatty acids. ELO4 produces very long-chain fatty acids (up to 26 carbons in length) ^20^ and ELO5 acts as a polyunsaturated fatty acid elongase ^19^. Apart from the initial characterization of the microsomal ELO pathway in *T. brucei* ^13^ and the demonstration of functional ELOs in *T. cruzi* ^13,19,21^ and *Leishmania* ^10,13,19^, the relative contribution of this pathway toward lipidome maintenance in trypanosomatids has not been determined.

In this study, we examined the role of the ELO system in supporting lipid homeostasis in axenic *T. cruzi* epimastigotes (EPI). As 16- and 18-carbon FA species represent ∼60% of total FA species in this organism ^20,22,23^, we focused our analysis on the ELO pathway enzymes, ELOs 1-3, which are expected to produce C16 and C18 from SCFA substrates in a processive pathway ^13^. Unbiased lipidomic analyses of loss-of-function mutants, Δ*elo1*, Δ*elo2* and Δ*elo3*, revealed that ELO2 and ELO3 both contribute significantly to the maintenance of global lipidomic profiles in *T. cruzi*. Despite marked lipidome remodeling in Δ*elo2* and Δ*elo3* EPI, lipidomic changes were well-tolerated with little impact on the growth of these mutants in culture. In contrast, ELO1 was found to be dispensable for LCFA synthesis and global lipidome maintenance. Instead, ELO1 expression was required to support mitochondrial metabolism and proliferation. Further analysis revealed that loss of ELO1 function was associated with decreased protein lipoylation which was rescued by octanoic acid (C8:0) supplementation or genetic complementation with functional *elo1*. Together, these findings highlight the modular nature of the trypanosomatid ELO pathway and reveal an unexpected role for ELO1 in supporting aspects of mitochondrial metabolism in *T. cruzi* EPI, including impact on the generation of SCFA precursors for lipoic acid synthesis, a function normally attributed to mitochondrial FAS-II.

## Results

### Generation of ELO-deficient T. cruzi mutants

The *T. cruzi* fatty acid elongase genes, *elo1* (TcCLB.506661.30/TcCLB.511245.130) *elo*2 (TcCLB.506661.20/TcCLB.511245.140) and *elo*3 (TcCLB.506661.10/TcCLB.511245.150) (Fig. 1A), were individually targeted for disruption in axenic *T. cruzi* epimastigotes (EPI) using CRISPR/Cas9 and integration of homology-directed repair cassettes (Fig. 1B). Cloning of transfected EPI during initial drug selection post-transfection facilitated recovery of parasites with insertions in both alleles with the successful generation of three independent ELO knockouts, Δ*elo*1, Δ*elo*2, and Δ*elo*3, as confirmed by Southern blot (Fig. 1C) and PCR (Fig. 1D). Genetic complementation of each Δ*elo* mutant was achieved with a modified pTREX expression vector encoding a C-terminal GFP-tagged copy of the relevant full-length *elo* gene on the mutant background (Fig. 1E). Consistent with the reported microsomal localization of the ELO pathway enzymes in *T. brucei* ^13^, all ELO-GFP proteins localized to the endoplasmic reticulum in *T. cruzi* EPI as confirmed by colocalization with the ER chaperone BiP ^24^ (Fig. 1E).

**Figure 1.**
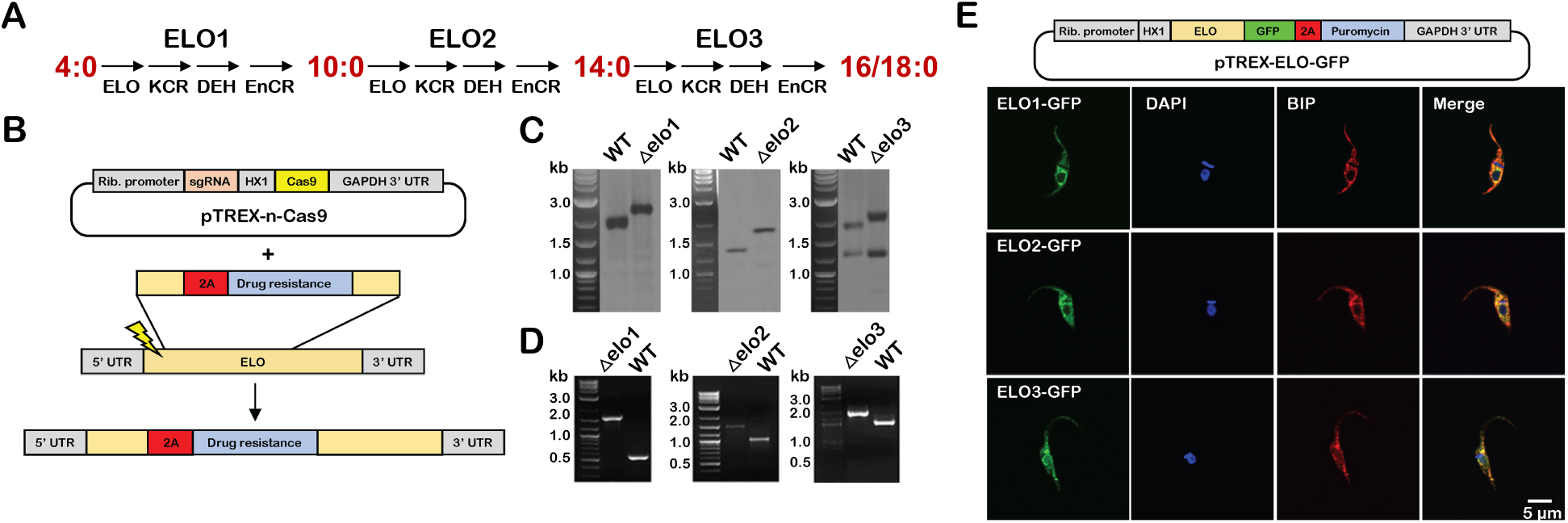
Generation of ELO-deficient and genetically-complemented *T. cruzi* lines. **A**. Schematic of ELO pathway with main FA substrates and products shown (red). ELO: elongase, KCR: ketoacyl-CoA reductase, DEH: hydroxyacyl-CoA dehydrase, EnCR: trans-2-enoyl-CoA reductase. **B**. General strategy for the targeted disruption of genomic loci in *T. cruzi* epimastigotes using a CRISPR/Cas9-mediated homology-directed repair approach. **C**. Southern blots of digested genomic DNA isolated from WT or different *T. cruzi elo* mutants and **D**. gDNA PCRs confirm integration of KO constructs at the correct locus. **E**. Schematic of pTREX expression vector used for genetic complementation of Δ*elo* mutants. Expression of GFP-tagged ELOs in genetically complemented Δ*elo* mutants epimastigotes reveals overlap of the ELO-GFP (green) signal with the endoplasmic reticulum chaperone BiP (red). Parasite nuclear and kinetoplast DNA were visualized using DAPI (blue).

### Lipidomic characterization of T. cruzi ELO mutants

As the main system used for bulk long-chain fatty acids (LCFA) synthesis in trypanosomatids, the ELO pathway is assumed to play a critical role in membrane lipid synthesis and remodeling in this group of organisms. Still, the contribution of this pathway in shaping the lipidome in these organisms has yet to be determined experimentally. To evaluate the impact of disruption of ELO pathway enzymes on the lipid composition of *T. cruzi* parasites, we performed label-free, quantitative ultra-high performance liquid chromatography high-resolution tandem mass spectrometry (UHPLC-HR-MS/MS) of lipids extracted from WT, Δ*elo*1, Δ*elo*2, Δ*elo*3 and genetically-complemented EPI lines. A total of 1,133 lipid species were identified after manual curation of LipidSearch-assigned IDs (Table S1), the majority of which were found to be significantly altered in abundance (p<0.05; one-way ANOVA) across all mutants as compared to WT EPI (Fig. 2A). Two-dimensional principal component analysis (PCA) identified overall trends in the lipidome data (Fig. 2B), where the Δ*elo*3 mutant emerged as the distinct outlier, well separated from the other parasite lines along both PC1 and PC2 (Fig. 2B). The Δ*elo*1 EPI lipidome was the least divergent from WT and Δ*elo*2 separated from both of these lines along PC2 (Fig. 2B). When broken down according to major lipid subclass (Fig. S1) PCA clustering patterns were found to be similar to that observed for the total lipidome (Fig. 2B), indicating that global lipidomic differences between of WT and Δ*elo* mutant *T. cruzi* EPI are likely not driven by specific lipid subclasses. Volcano plots displaying pairwise comparisons of WT and individual Δ*elo* mutants (Fig. 2C) and heatmap data highlighting the 50 lipids that changed most dramatically in the mutants as compared to WT EPI (Fig. 2D) demonstrate that lipidomic changes are most prominent in Δ*elo2* and Δ*elo3* mutants. Consistent with this conclusion, 232 and 149 lipids were found to be changed ≥2 or ≤0.5 -fold change (p-value <0.05) in Δ*elo2* and Δ*elo3* respectively (Table S2). In contrast, comparatively few lipids (48) exhibited ≥2 or ≤0.5 -fold change in abundance in the Δ*elo1* mutant when compared to WT EPI (Fig. 2C,D) (Table S2).

**Figure 2.**
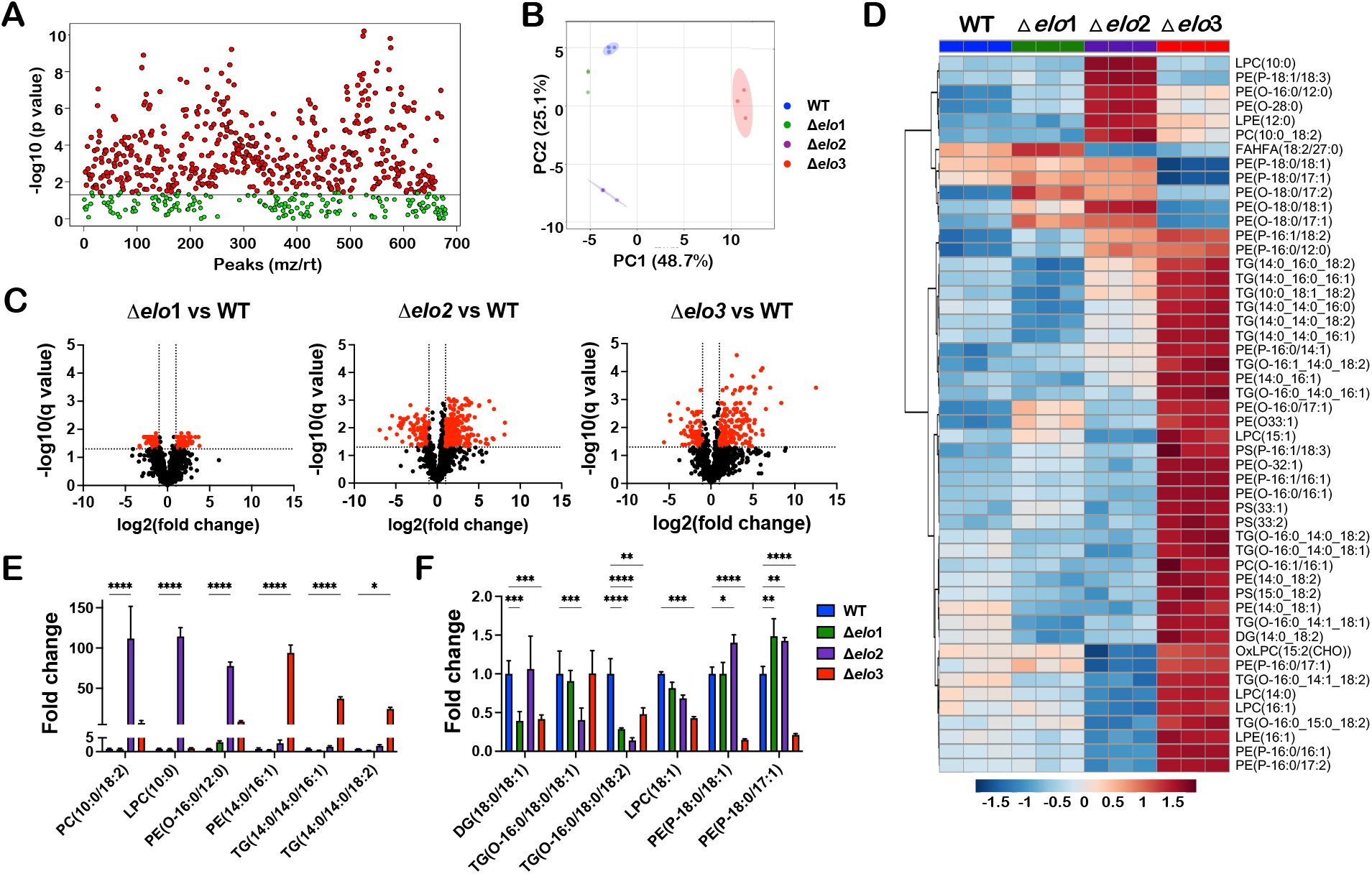
Lipidomic characterization of *T. cruzi* Δ*elo* mutants. Lipids identified in WT and Δ*elo* mutant *T. cruzi* epimastigotes (Folch’s lower phase) using UHPLC-HR-LC-MS/MS **A**. *One-way ANOVA* with Tukey’s post-hoc analysis showing significant differences in abundance (p<0.05). **B**. Principal component analysis (PCA) of total lipidomes plotted for WT, Δelo1, Δelo2 and Δelo3 *T. cruzi* EPI. The first two principal components are shown (PC1 and PC2) with the proportion of variance for each component in parenthesis. Biological replicates for each experimental group are represented and the 95% confidence interval indicated in shaded circle. **C**. Volcano plots based on fold-change (x-axis) and adjusted p-value (q-value; y-axis) reveal differences in lipid profiles. Horizontal line indicates significant q value (q≤0.05), vertical lines indicate ≥2-fold change-≤0.5 cut-off. Lipids that are significantly altered in abundance (≥2-fold change or fold change-≤0.5; adjusted p-value ≤0.05) are represented as red dots in each pair-wise comparison. **D**. Heat map displaying the 50 most significantly up or down regulated lipids when comparing Δ*elo* mutants with WT EPI. Highlighted examples of lipids in the dataset that are significantly increased (**E**) or decreased (**F**) in one or more Δ*elo* mutant as compared to WT *T. cruzi* EPI. Two-way ANOVA with Tukey’s multiple comparisons test was applied (*p<0.05, **p<0.01, ***p<0.001, ****p<0.0001). PE, phosphatidylethanolamine; TG, triacylglycerol; PS, phosphatidylserine; Cer, ceramides; DG, diacylglycerol; PC, phosphatidylcholine; LPC, *lyso*-phosphatidylcholine; LPE, *lyso*-phosphatidylethanolamine; PI, phosphatidylinositol; FAHFA, fatty acyl esters of hydroxy fatty acids.

Consistent with the anticipated function of ELO3 in extending 14-carbon fatty acids to C16 and C18 species, we noted a marked reduction in C18:0 and C18:1-containing lipids in Δ*elo*3 EPI and a concomitant increase in lipids containing C14:0, C14:1, C16:0, and C16:1 in this mutant (Fig. 2D,E,F). A similar trend was observed for Δ*elo*2 EPI, where lipids containing C10:0 and C12:0 were elevated and lipids containing C14:0 and C16:0 FA were present at decreased levels (Fig. 2D,E,F). Given that the majority of lipid alterations observed in the Δ*elo* mutants were restored in the genetically complemented lines (Fig. S2), we conclude that the observed lipidomic changes are due to the loss of specific ELO enzyme function. Combined, these data provide the first demonstration that ELO2 and ELO3 play important homeostatic roles in lipidome maintenance in *T. cruzi*, but that ELO1 contributes little to LCFA synthesis and global lipidomic profiles in this organism.

### Metabolic labeling and FA substrate elongation in WT and Δelo mutants

In addition to *de novo* synthesis of fatty acids, *T. cruzi* take up lipids from the environment and can incorporate exogenous fatty acids into their own lipids ^14,25–28^. The observation that ELO2- and ELO3-deficient EPI experience global lipidomic changes, characterized by a decrease in lipids containing 18-carbon FA side chains, despite continuous culture in medium containing 10% serum, suggests that serum lipids/exogenous fatty acids are insufficient to overcome loss of ELO activity. To examine the possibility that *T. cruzi* Δ*elo* mutants may experience defects in their ability to take up or utilize exogenous FA for lipid synthesis, we performed metabolic tracing studies using [1-14C]-labeled fatty acids of varying chain lengths, C4:0, C14:0, C16:0, C18:0. Lipids extracted from ^14^C-FA labeled parasites were resolved using reverse-phase thin layer chromatography (RP-TLC) to identify major lipid classes into which label was incorporated (Fig. 3A,B). Although differences in label intensity were observed when different ^14^C-FA were used for labeling, ^14^C-label was incorporated into neutral (Fig. 3A) and polar (Fig. 3B) lipids across all parasite lines (Fig. 3A,B). A notable exception was the Δ*elo1* mutant which failed to incorporate carbons from ^14^C-butyric acid (C4:0) into parasite lipids (Fig. 3A,B).

**Figure 3.**
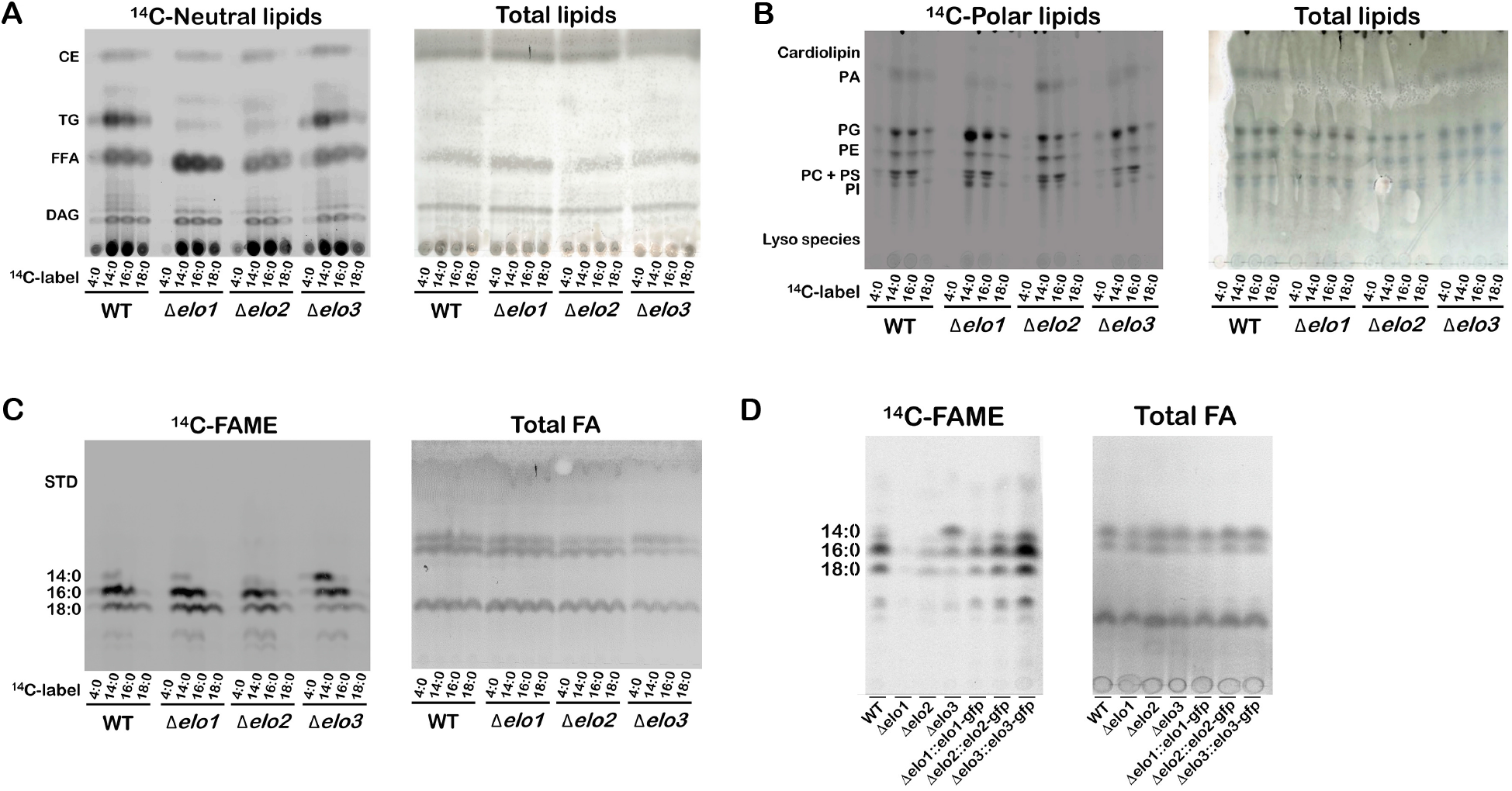
Elongation and utilization of exogenous fatty acid substrates by *T. cruzi* WT and Δ*elo* mutants. Metabolic tracing showing exogenous FA incorporation and elongation in WT and Δ*elo* mutants by thin layer chromatography (TLC). Exponentially growing epimastigotes were incubated with 0.3 μCi/ml of different 1-^14^C-FA, as indicated (^14^C-label) for 18 h prior to lipid extraction. Left and right panels show ^14^C-labeled lipids and total lipids (sulfuric acid-copper staining), respectively. Neutral and polar lipids (**A**,**B**, respectively) isolated from 1-^14^C-FA labeled *T. cruzi* EPI and resolved by RP-TLC and exposed to film for 3 days. **C**. Fatty acid methyl esters (FAME) derived from 1-^14^C-FA-labeled lipids resolved by RP-TLC; film exposed for 3-day. **D**. ^14^C-FAME resulting from labeling of *T. cruzi* WT, Δ*elo* mutants and genetically-complemented lines with ^14^C-butyric acid, resolved by RP-TLC and exposed to film for 10 days.

To more directly assess the impact of specific ELO-deficiencies on FA elongation in intact parasites, fatty acid methyl esters (FAMEs), generated from total lipids extracted from each parasite line following ^14^C-FA labeling were resolved by RP-TLC (Fig. 3C). As expected, carbons from exogenous [1-14C]-FA were detected in LCFA (C14:0, C16:0, and C18:0) in all parasite lines, except for Δ*elo1*, which failed to incorporate ^14^C from labeled butyric acid into LCFA species (Fig. 3C,D). Given the relatively faint signal in WT parasites obtained when ^14^C-butyric acid was used as a metabolic tracer (Fig. 3A-C), additional labeling experiments were performed to optimize detection of labeled lipids following incubation with ^14^C-butyric acid (Fig. 3D). Results confirm that failure to incorporate ^14^C from butyric acid into LCFAs is unique to the Δ*elo1* mutant but was fully restored in genetically-complemented Δ*elo1::elo1*-*gfp* parasites (Fig. 3D). Given that Δ*elo1* EPIs readily elongate ^14^C-myristic acid (C14:0) and ^14^C-palmitic acid (C16:0) substrates (Fig. 3C), it is unlikely that impaired import of fatty acids explains the inability of this mutant to elongate ^14^C-butyric acid. Together, these results confirm the function of *T. cruzi* ELO1 as a short-chain fatty acid (SCFA) elongase that utilizes butyric acid (butyryl-CoA) as a substrate for elongation ^29^. Moreover, the unimpeded ability of Δ*elo1* EPI to elongate C14:0 and C16:0 substrates, presumably through the actions of ELO2 and ELO3, is consistent with a modular ELO system where individual ELOs can function independently of the other pathway enzymes ^29^.

The results of metabolic labeling and ^14^C-FAME analysis also point to overlapping functions and partial redundancy among the ELOs examined here. For example, in the absence of ELO2 expression, *T. cruzi* EPI can still generate ^14^C-labeled LCFA species from ^14^C-butyric acid, suggesting that some extended capabilities of ELO1 and/or ELO3 can compensate to some degree for the loss of ELO2 (Fig. 3C). Similarly, despite the notable accumulation of ^14^C-labeled 14:0 fatty acid species in Δ*elo*3 EPI following incubation with ^14^C-myristic acid (Fig. 3C) or ^14^C-butyric acid (Fig. 3D), consistent with an impaired capacity of the Δ*elo*3 mutant to utilize C14:0 as a substrate for elongation, conversion of ^14^C-myristic acid to ^14^C-labeled 16:0 and 18:0 still occurs in this mutant (Fig. 3A). Functional overlap and redundancy within the ELO pathway may explain why disruption of ELOs individually fails to promote more profound lipidome perturbation in the knockout parasite lines.

### ELO1 deficiency is associated with impaired growth of T. cruzi epimastigotes

To evaluate the biological impact of ELO gene disruption in *T. cruzi*, we first examined the growth kinetics of Δ*elo1*, Δ*elo2*, and Δ*elo3* EPI in LIT medium containing 10% serum (Fig. 4). Strikingly, ELO1-deficient *T. cruzi* EPI exhibit marked growth impairment in the log-phase (doubling time of 60 ± 12 h) compared to WT parasites (31 ± 7 h). Δ*elo2* EPIs exhibit a mild growth defect (41 ± 9 h), whereas the Δ*elo3* mutant replicates at rates similar to WT (Fig. 4). The reduced rates of growth observed for Δ*elo1* and Δ*elo2* EPIs do not involve changes in the parasite cell cycle, as the proportion of parasites associated with different phases of the cell cycle (G1-S-G2/M) remained unaltered in each mutant as compared to WT (Fig. S3). The growth deficits observed for Δ*elo1* and Δ*elo2* mutants were rescued following stable expression of ELO1-GFP or ELO2-GFP in the respective mutant lines (Fig. 4). As expected, growth of genetically complemented Δ*elo*3 parasites exhibited similar kinetics as WT and Δ*elo3* EPI (Fig. 4). In contrast to genetic complementation, attempts to rescue growth of Δ*elo1* or Δ*elo2* EPI by supplementation of the medium with exogenous fatty acids of varying chain lengths (14:0, 16:0, 18:0, or 18:1) failed to fully restore growth of these mutants, despite mild beneficial effects of supplemental C18 FAs on parasite growth (Fig. S4).

**Figure 4.**
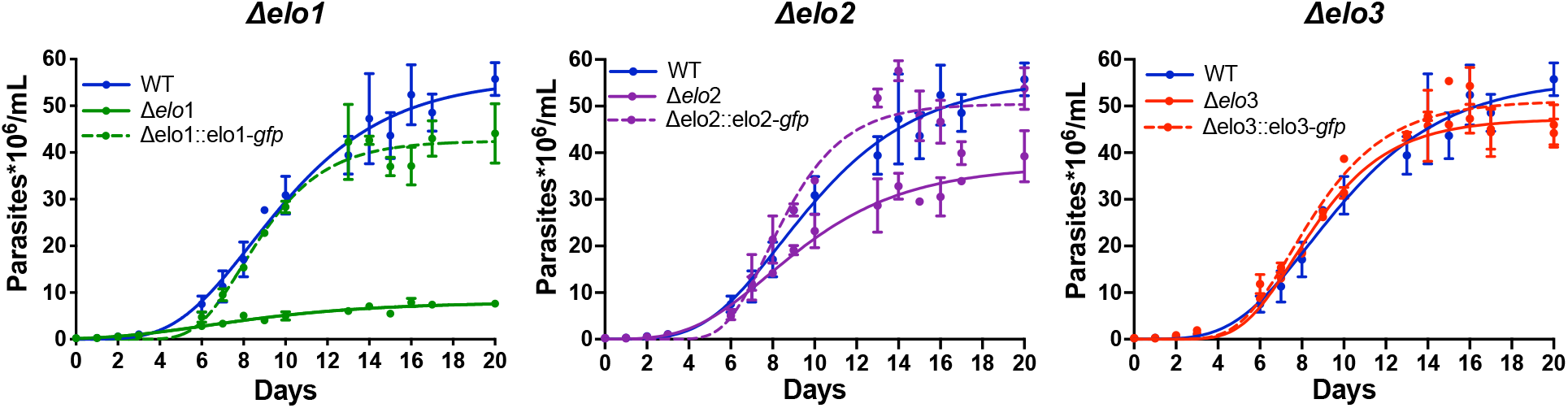
Growth characteristics of *T. cruzi* Δ*elo* mutants and complemented lines. The growth of WT, Δ*elo* mutants and genetically-complemented *T. cruzi* EPI lines in LIT medium containing 10% FBS was followed for 20 days. The growth of each Δ*elo* mutant and cognate genetically complemented line as compared to WT EPI is shown on separate graphs. The mean and standard deviation calculated for parasite density (parasites × 10^6^/mL medium; y-axis) for 3 biological replicates against time (days; x-axis) is plotted.

Overall, our data demonstrate that when targeted individually, ELO1, ELO2, and ELO3 are dispensable for *T. cruzi* EPI growth and that the extent of lipidome perturbation resulting from loss of ELO activity correlates poorly with growth phenotypes in the Δ*elo* mutants. Thus, loss of ELO1 function may promote specific changes in the parasite lipidome not captured in our analysis and/or other metabolic changes that lead to growth impairment in Δ*elo1* and to a lesser extent Δ*elo2* EPI.

### ELO1-deficiency leads to metabolic changes and mitochondrial dysfunction

To examine whether disruption of specific fatty acid elongases in *T. cruzi* triggered other metabolic changes in the Δ*elo* mutants that might explain observed growth phenotypes, we performed an unbiased metabolomic analysis of log-phase WT EPI, each Δ*elo* mutant and the respective genetically-complemented lines. LC-MS/MS analysis identified ∼300 metabolites with high confidence in *T. cruzi* EPI extracts (Table S3). PCA allowed visualization of the overall trend in the metabolite data, where WT and Δ*elo*3 EPI clustered together and the Δ*elo*2 and Δ*elo*1 mutants formed distinct clusters that were well separated from WT (Fig. 5A). These trends were reflected in the heatmap generated with the top 50 metabolites found to be altered in abundance when Δ*elo* mutants were compared to WT *T. cruzi* EPI (Fig. 5B) and visualized as pair-wise comparisons in volcano plots (Fig. 5C). Among the mutants, Δ*elo1* exhibited the greatest degree of metabolic perturbation with significant changes occurring for 74 metabolites (<u>≥</u>2 or ≤0.5 fold-change; q-value <0.05), 48 of which returned to WT levels in Δ*elo1*:*:elo1-gfp* EPI (Fig. S5). These changes included metabolites associated with glucose metabolism, the pentose phosphate pathway, nicotinamide synthesis, and the citric acid cycle (Fig. 5D, Table S4). By comparison, 37 metabolites were significantly altered in abundance in Δ*elo2* EPI, with the majority restored to WT levels in Δ*elo*2:*:elo*2*-gfp* parasites. In contrast, the Δ*elo*3 mutant showed little evidence of metabolic perturbation with only 6 metabolites exhibiting significant changes in abundance as compared to WT. Collectively, these data suggest that metabolite perturbation, and not global lipidomic alterations, correlate with reduced growth phenotypes in Δ*elo1* EPI. Elevation of specific metabolites such as 2-deoxyglucose-6-phosphate (6-fold) and malonyl-CoA (50-fold) in Δ*elo1* EPI (Fig. 5D, Table S4), which have the potential to inhibit glycolysis and fatty acid oxidation, respectively, may be indicative of dysregulated energy metabolism in this mutant.

**Figure 5.**
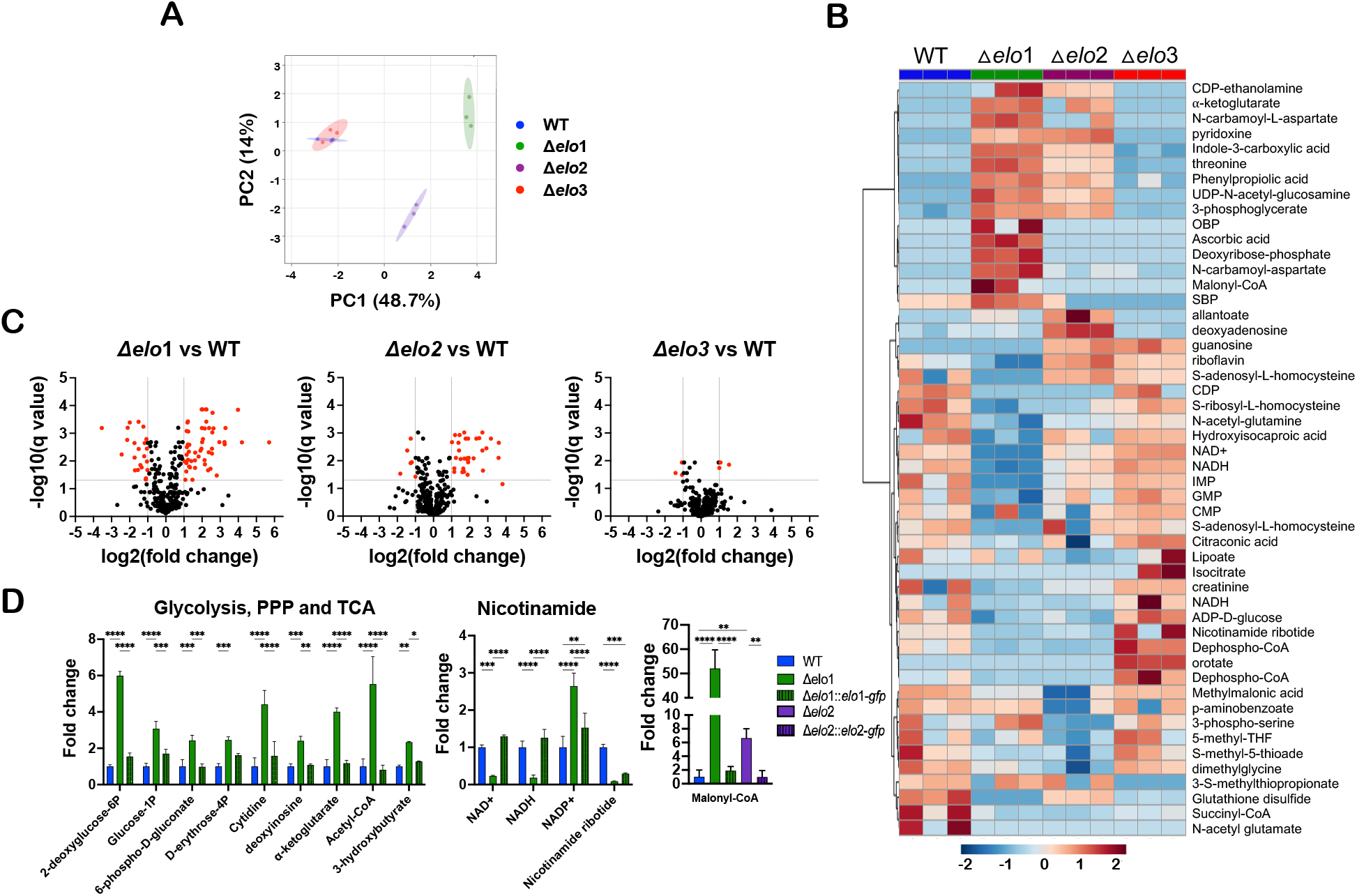
Metabolite perturbation in Δ*elo* mutant *T. cruzi* epimastigotes. Principal component analysis (PCA) scores plot **(A)** and heat map **(B)** comparing steady state metabolite profiles of *T. cruzi* WT and Δelo mutant EPI. **C**. Volcano plots based on fold-change (x-axis) and adjusted p-value (q-value; y-axis). Horizontal line represents significant q value (q≤0.05), vertical lines indicate ≥2-fold change-≤0.5 cut-off. Highlighted metabolites (red circles) are significantly altered in abundance (≥2-fold change or fold change-≤0.5; adjusted p-value ≤0.05) in pair-wise comparisons of WT EPI and individual Δ*elo* mutant strains. **D**. Relative abundance of metabolites shown to be significantly altered in Δ*elo1* mutant EPI and restored to WT levels in Δ*elo1::elo1-gfp*. Two-way ANOVA with Tukey’s multiple comparisons test was applied (*p<0.05, **p<0.01, ***p<0.001, ****p<0.0001). Results show metabolites from different metabolic pathways significantly changing in Δ*elo*1 and less extent in Δ*elo*2 EPI.

Malonyl-CoA is a pivotal molecule in the regulation of fatty acid synthesis and oxidation ^30^. It is generated from acetyl-CoA and is the 2-carbon donor used in the synthesis/elongation of fatty acids. Malonyl-CoA also inhibits fatty acid oxidation (FAO) in the mitochondria by blocking the CPT1-dependent import of LCFA into the mitochondria ^31,32^. To determine if FAO is impaired in *T. cruzi* Δ*elo* mutants, log-phase EPI were incubated for 4 h with ^3^H-palmitate (C16:0) in the absence of glucose, followed by the measurement of ^3^H_2_O released into the supernatant. We observed a significant reduction of FAO in Δ*elo1* EPI that was restored in the genetically complemented line, whereas no significant differences were observed in FAO in Δ*elo*2 and Δ*elo*3 mutants as compared to WT EPI (Fig. 6A).

**Figure 6.**
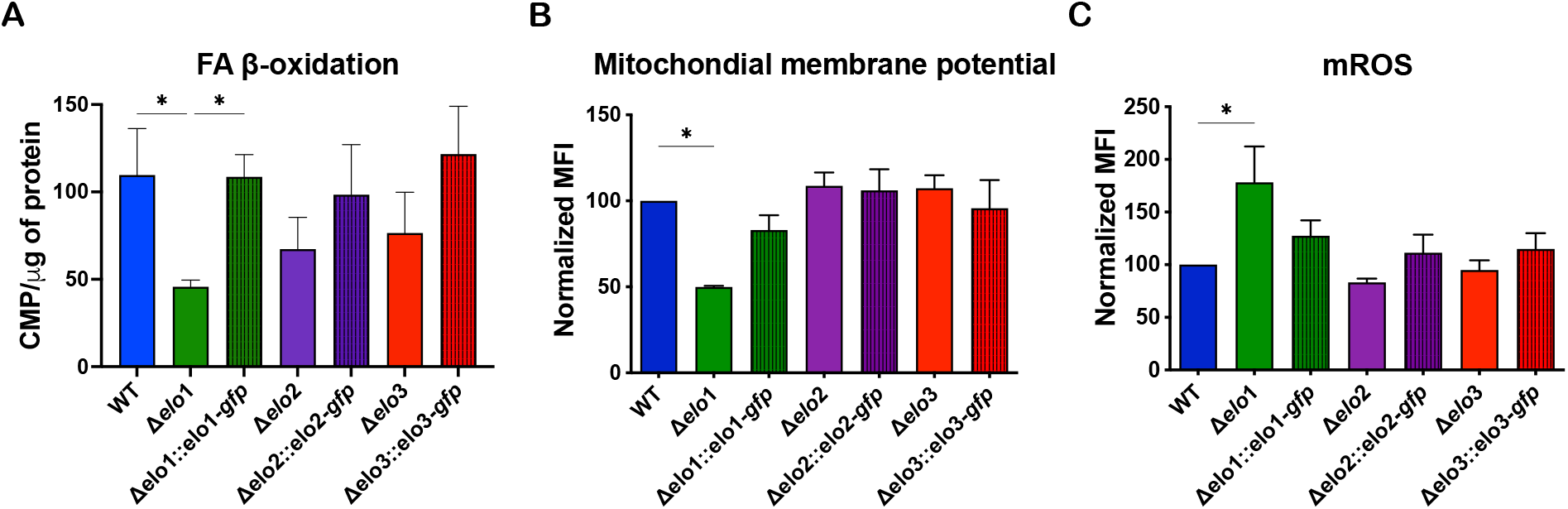
Fatty acid β-oxidation and mitochondrial function. **A**. Fatty acid oxidation was measured in all *T. cruzi* EPI lines following the detection of ^3^H_2_0 released from parasites into the culture supernatant after a 4 hours incubation of log-phase EPI with ^3^H-palmitate. Counts were acquired for 5 minutes in a scintillation counter and normalized to μg of protein in the pelleted parasites. **B**. Mitochondrial membrane potential measured in *T. cruzi* EPI lines by TMRE staining and flow cytometric analysis. **C**. Mitochondrial reactive oxygen species measured by MitoSOX™ staining and analyzed by flow cytometry. One-way ANOVA with Dunnett’s multiple comparisons test was applied (*p<0.05).

Reduced FAO and changes in abundance of metabolites involved in nicotinamide and NAD synthesis (Fig. 5D) led us to ask if mitochondrial function might be affected in the Δ*elo1* mutant. As general indicators of mitochondrial fitness, we measured mitochondrial membrane potential and mitochondrial reactive oxygen species (mROS) and found that both of these measures are altered in the Δ*elo1* mutant (Fig. 6B,C). Mitochondrial membrane potential was reduced in Δ*elo1* mutant by ∼50% as compared to WT parasites, and no significant changes in membrane potential were observed in Δ*elo*2 or Δ*elo*3 EPI (Fig. 6B). Additionally, mROS levels were differentially elevated in Δ*elo1* EPI (Fig. 6C) and both membrane potential and mROS levels restored to WT levels in genetically-complemented Δ*elo1::elo1-gfp* parasites (Fig. 6B,C). Thus, loss of ELO1 function leads to mitochondrial dysfunction in *T. cruzi* EPI, whereas the metabolic and/or lipidomic changes accompanying the loss of ELO2 or ELO3 do not impinge on mitochondrial function. Reduced mitochondrial membrane potential in Δ*elo1* EPI could indicate reduced electron transport chain (ETC) activity, possibly through the depletion of NADH and FADH2 produced in the TCA cycle and during FAO.

### ELO1-deficiency affects protein lipoylation

The metabolic tracing experiments above confirmed that ELO1-deficient *T. cruzi* EPI are defective in their ability to elongate butyric acid (C4:0). To explore the possibility that the loss of FA products or intermediates generated by ELO1 (e.g., C8:0 and C10:0) might underlie growth and mitochondrial phenotypes observed in Δ*elo*1 mutant EPI, growth curve experiments were conducted in complete LIT medium supplemented, or not, with 35 μM octanoic acid (C8:0) or decanoic acid (C10:0). C10:0 supplementation failed to alter growth of WT or Δ*elo*1 EPI, whereas C8:0 exerted a small but significant growth-enhancing effect on this mutant (Fig. 7A). As octanoic acid (C8:0) is used to synthesize lipoic acid, a co-factor needed for the stabilization and activity of mitochondrial enzymes such as pyruvate dehydrogenase, a key regulator of oxidative metabolism in mitochondria ^33^, we sought to determine whether protein lipoylation levels were altered in Δ*elo*1 mutant EPI. As a covalent post-translational modification, lipoylation is readily detected by western blot using an antibody that recognizes lipoic acid ^18^. Using this approach, several bands corresponding to lipoylated proteins were detected in protein lysates prepared from WT *T. cruzi* EPI, as previously reported ^18^. The protein lipoylation signal, normalized to tubulin, was significantly reduced in Δ*elo*1 mutant EPI (49%) but restored to WT levels in genetically-complemented Δ*elo*1::*elo*1-*gfp* EPI (Fig. 7B). Supplementation of the medium with C8:0 partially restored protein lipoylation in the Δ*elo*1 mutant whereas supplemental C10:0 had no detectable effect on protein lipoylation in WT EPI or the Δ*elo*1 mutant (Fig. 7B). These results provide the first evidence of a functional link between ELO1 activity and protein lipoylation in trypanosomatids.

**Figure 7.**
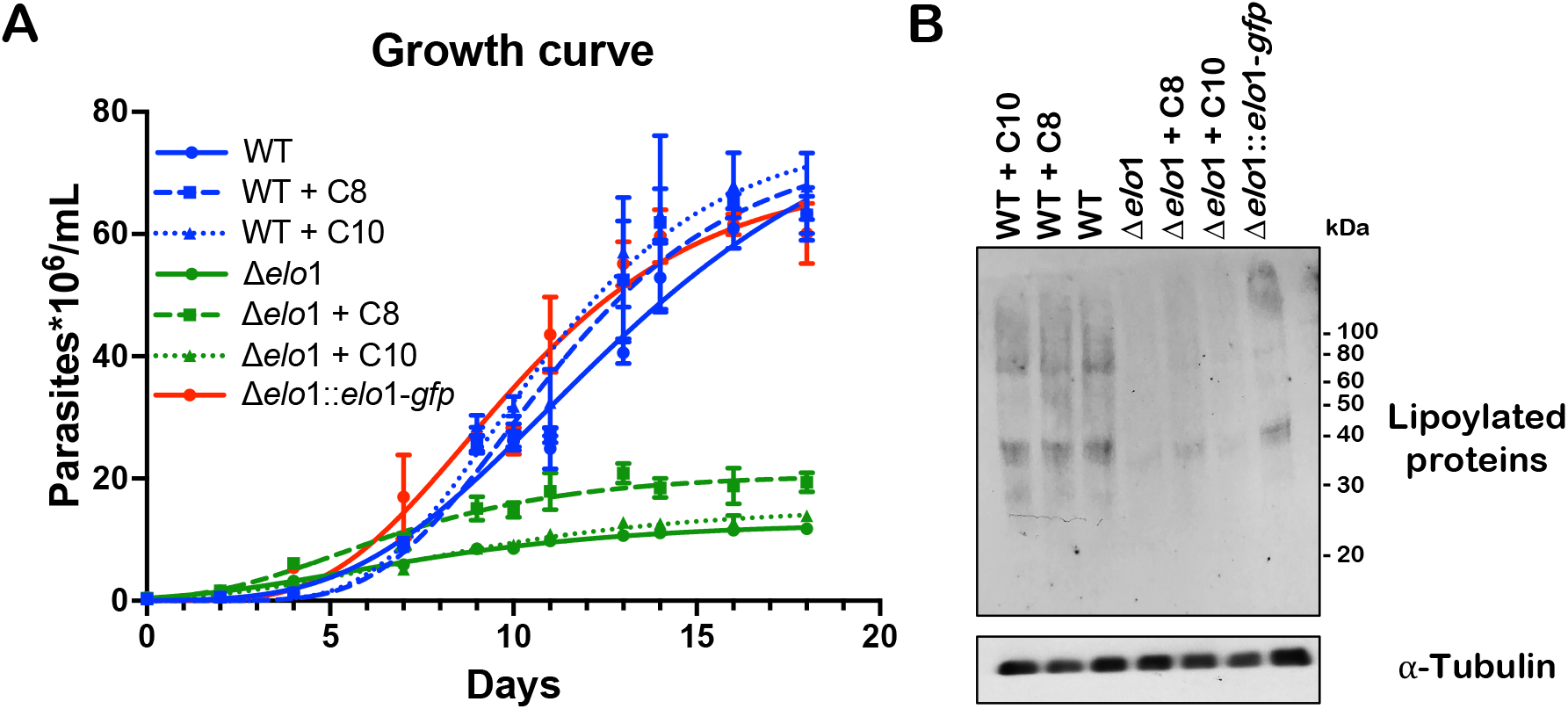
Protein lipoylation is impaired in Δ*elo*1 mutants. **A**. Growth curves of WT, Δ*elo*1 and Δ*elo*1::*elo*1 epimastigotes grown in LIT media with and without supplementation with 35 μM C8:0 or C10:0. **B**. Western blot of lysates of *T. cruzi* EPI harvested on the final day of the growth curve (A) in the 35 μM of C8 or C10 FA and probed with antibodies to lipoic acid (top). Blots were stripped and re-probed with anti-Tubulin antibody (bottom).

## Discussion

Long-chain fatty acids (LCFA) play a critical role in the growth and developmental cycle of *Trypanosoma cruzi*, where 16- and 18-carbon species are the most abundant FA in this organism ^12,20,22,23^. Like many cells, *T. cruzi* has the capacity to synthesize LCFA *de novo* and to scavenge FA from the environment ^12,28^. Bulk fatty acid synthesis, via the ELO system, takes place at the cytoplasmic face of the endoplasmic reticulum and is physically and biochemically distinct from the mitochondrial FAS-II system ^12,13^. In this study, we generated genetic mutants to examine the role(s) of the first three fatty acid elongases in the *T. cruzi* ELO system, ELO1, ELO2, and ELO3, predicted to be responsible for the processive generation of C16:0 and C18:0 from C4:0 (butyryl-CoA) ^13^. Phenotypic analysis of individual Δ*elo* mutants and their genetically-complemented lines has provided novel perspectives on the roles of these ELOs in lipidome maintenance and parasite growth.

Our metabolic tracing studies confirmed that WT *T. cruzi* EPI possess a functional ELO system that can generate LCFA from C4:0 in a processive manner. The short chain fatty acid elongase, ELO1, is required to elongate C4:0 precursors, as this capacity was abolished in Δ*elo*1 mutant *T. cruzi* EPI and restored with genetic complementation. Despite its role in the generation of LCFA from C4:0, ELO1 is not required for global lipidome maintenance in *T. cruzi* EPI, as ELO1-deficient parasites can still produce C18:0 from C14:0 and C16:0 precursors, presumably via the actions of ELO2 and/or ELO3. These findings argue that operation of a processive ELO system (e.g., from C4:0 to C18:0) is not essential for lipidome maintenance in *T. cruzi* EPI and predict similar outcomes for related trypanosomatids, for which the global impact of ELO knockout has not been determined ^13^.

In contrast to ELO1, both ELO2 and ELO3 exerted a clear influence over the lipidome in *T. cruzi* EPI. Marked changes in the lipidome profiles of both Δ*elo*3 and Δ*elo*2 mutant EPI were consistent with the loss of specific ELO activities. For instance, complex lipids (e.g., phospholipids, sphingolipids, neutral lipids) containing C16 and C18 FA moieties were found to be significantly diminished in Δ*elo*3 and Δ*elo*2 EPI, whereas lipids containing the uncommonly used C10, C12, and C14 fatty acyl species were elevated in these mutants. These global lipidomic changes were attributable to the loss of specific ELO function as reversion to WT profiles was evident in the respective genetically-complemented lines. Notably, global lipidomic changes were observed in Δ*elo*3 and Δ*elo*2 mutant EPI despite continuous passage of parasites in medium containing serum. Thus, serum lipids and exogenous fatty acids appear to be insufficient to compensate for the loss of ELO2 or ELO3 function and the resultant lipidomic changes occurring in these mutants. Metabolic tracing studies demonstrate the ability of Δ*elo* mutants to incorporate ^14^C-labeled LCFA precursors into phospholipid and neutral lipid pools at levels similar to WT EPI, although we cannot exclude the possibility that the Δ*elo*3 and Δ*elo*2 mutants exhibit quantitative differences in the import or utilization of exogenous FA.

One of the unexpected findings of this study was the inverse relationship between global lipidome perturbation and growth of *T. cruzi* Δ*elo* mutants. ELO1-deficient parasites exhibited severe growth impairment with relatively few lipidomic changes, whereas Δ*elo*2 and Δ*elo*3 mutants showed little or no impairment of growth, respectively, despite clear lipidome perturbation. Notably, media supplementation with exogenous LCFA failed to rescue growth inhibition observed in Δ*elo*1 or Δ*elo*2 mutant EPI, despite evidence of uptake and incorporation of exogenous FA into parasite neutral and polar lipids. Combined, these observations indicate that the severe growth defect associated with the loss of ELO1 expression is unlikely to be related to the failure of this mutant to generate, or to acquire, LCFA.

Accordingly, changes in polar metabolite profiles in log-phase Δ*elo* mutant EPI were found to be better aligned with growth phenotypes of these mutants. Specifically, the greatest metabolic perturbation occurred in the Δ*elo1* mutant (impaired growth) while minimal changes occurred in the Δ*elo3* mutant (no growth impairment). Of the 74 metabolites that exhibited significant changes in abundance in Δ*elo*1 EPI, the most striking was malonyl-CoA, which increased 50-fold over WT in the Δ*elo*1 mutant and was restored to near WT levels in genetically-complemented parasites. Given the pivotal role of malonyl-CoA in fatty acid and energy metabolism, the accumulation of this metabolite is expected to trigger dramatic metabolic changes in the parasite. Malonyl-CoA is consumed in the synthesis of FA (in both the ELO and mitochondrial FAS-II systems) and conditions that lead to high levels of malonyl-CoA are known to suppress fatty acid entry into mitochondria through the direct inhibition of carnitine palmitoyltransferase (CPT-1) ^34–36^. Consistent with the anticipated metabolic consequences of malonyl-CoA accumulation, fatty acid oxidation (FAO) was significantly decreased in Δ*elo*1 mutant EPI and restored to WT levels in genetically-complemented Δ*elo*1::*elo*1-*gfp* parasites. Why malonyl-CoA accumulates to such a great degree in ELO1-deficient parasites, but not in the other ELO knockouts, is perplexing, given that this two-carbon donor is utilized in the elongation of fatty acyl-CoA substrates by all ELO enzymes. One possibility is that the ELO1-dependent fatty acid elongation cycle functions at a higher rate than the combined rates of all other ELOs, leading to excess malonyl-CoA in the absence of ELO1. Alternatively, excess malonyl-CoA may be symptomatic of reduced mitochondrial FAS-II activity in the Δ*elo*1 mutant.

Certainly, there are indicators of mitochondrial dysfunction in ELO1-deficient *T. cruzi* EPI, including reduced mitochondrial membrane potential, increased ROS production and significant alteration in levels of metabolites related to energy and redox metabolism. In addition, the finding that ELO1 expression impacts protein lipoylation further connects this short-chain fatty acid elongase to mitochondrial metabolism. The activity of several mitochondrial enzymes, including pyruvate dehydrogenase and α-ketoglutarate dehydrogenase, requires the post-translational attachment of lipoic acid (6,8-dithiooctanoic acid) which is derived from octanoic acid (C8:0) ^18,33^. Protein lipoylation is significantly diminished in Δ*elo*1 EPI and partially restored by the addition of C8:0, an intermediate product of both ELO1-dependent FA elongation and of mitochondrial FAS-II activity. While octanoic acid used for the synthesis of lipoic acid is assumed to be produced by FAS-II ^18,33,37^, our results reveal a previously unrecognized link between microsomal ELO1 activity and protein lipoylation in *T. cruzi* (Fig. 8). At this point, it is unknown whether octanoic acid generated by ELO1 directly contributes to lipoic acid pools in the parasite, or whether the global metabolic perturbation and mitochondrial dysfunction, demonstrated in ELO1-deficient parasites, impacts FAS-II activity indirectly. As supplemental C8:0 provides only a minor boost in growth to Δ*elo*1 EPI our data suggest that growth impairment accompanying loss of ELO1 function is likely the culmination of multiple metabolic and cellular changes.

**Figure 8.**
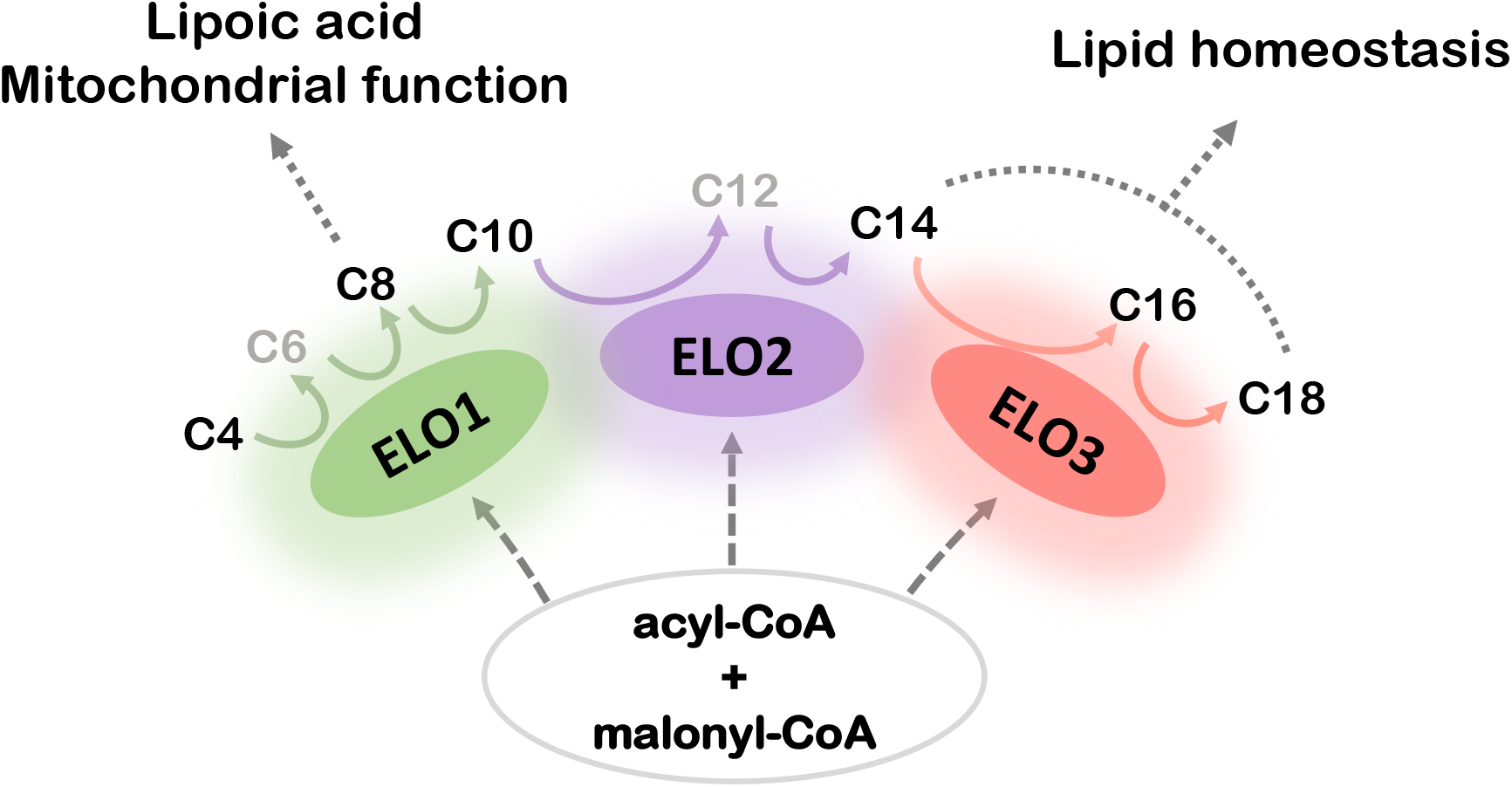
ELO pathway model. Scheme of the ELO pathway showing contribution of ELO enzyme to lipid homeostasis. Acyl-CoA and malonyl-CoA are the substrates for the ELO enzymes. Main substrates and products indicated in black, intermediate products indicated in gray. Glow indicates function overlapping between the ELO enzymes. Solid colored arrows match the main ELO enzyme activities. ELO1 elongates C4 up to C8, that can be presumably used for lipoic acid synthesis, or elongated to C10. ELO 2 elongates C10 up to C14, that will be taken up by ELO3 and elongated up to C16 or C18 and used for complex lipid synthesis. Exogenous FA can be incorporated at any level of the ELO pathway for further elongation.

Overall, this study reveals previously unrecognized complexity in the trypanosomatid ELO pathway. Although ELO1 can function in a processive system that produces LCFA via the serial elongation of SCFA and involving ELO2 and ELO3, our data indicate that LCFA synthesis and lipidome maintenance are not the primary functions of this enzyme. Instead, ELO1 activity is required to promote mitochondrial function and normal growth of *T. cruzi* EPI. Whether causal or symptomatic of other unidentified metabolic/physiologic changes in the parasite that occur with loss of ELO1, the link between ELO1 activity and mitochondrion is a novel finding that highlight the trypanosomatid ELO pathway as a critical metabolic regulator.

## Experimental procedures

### Reagents

Fatty acids were purchased from Sigma-Aldrich, St. Louis, MO, and 100x (3.5 mM) stocks of butyric acid, decanoic acid, myristic acid, palmitic acid, and stearic acid were prepared by dissolving each fatty acid in 2 mL ethanol at 60°C, followed by dilution in pre-warmed (37°C) PBS containing 100 mg/mL BSA. Oleic Acid solution was purchased suitable for cell culture. All radiolabeled compounds were purchased from American Radiolabeled Chemicals Inc., St. Louis, MO.

### Trypanosoma cruzi maintenance and growth curves

Tulahuén strain *Trypanosoma cruzi* (ATCC PRA-33) EPIs were grown axenically at 27°C in liver infusion tryptose (LIT), supplemented with 10% FBS, 100 U/mL penicillin, 10 μg/mL streptomycin, and maintained in log-phase by weekly passage. For comparative growth analysis, log-phase *T. cruzi* EPIs were diluted to a density of 1×10^6^ parasites/mL in LIT and incubated at 27°C for 15-20 days. 25 μL samples were collected every 24 h, fixed in 75 μL of 2% paraformaldehyde/PBS, and stored at 4°C. Once all samples were collected, the number of parasites present in 100 μL was counted using a MACSQuant VYB Flow Cytometer. For the biochemical supplementation experiments, LIT was supplemented with fatty acids at a final concentration of 35 μM, and parasite growth curves were derived.

### CRISPR/Cas9-mediated disruption of ELO genes in T. cruzi

Gene disruption was accomplished using a homology-directed repair, Cas9-mediated system modified from Lander et al ^38,39^. Two guide RNA targeting sequences were designed for each target gene using the EuPaGDT system ^40^. Mutations to the specificity of guides in pTREX-n-Cas9 were performed using a Q5 mutagenesis kit (NEB, Ipswich, Massachusetts). Guides targeting the ELO genes were inserted using the primers described in Table 1: Elongase 1 (*elo1*, TcCLB.506661.30, g-*elo*1-1 and g-*elo*1-2 for g163 and g-*elo*1-3 and g-e*lo*1-4 for g151. Elongase 2 (*elo*2, TcCLB.506661.20), g-*elo*2-1 and g-*elo*2-2 for g100 and g-*elo*2-3 and g-*elo*2-3 for g92. Elongase 3 (*elo3*, TcCLB.506661.10), g-*elo*3-1 and g-*elo*3-for g116 and g-*elo*3-3 and g-*elo*3-4 for g132. Donor DNA containing the drug resistance cassette was generated by PCR using ultramers with 100-bp homology to regions flanking the predicted Cas9 cut site and an in-frame P2A ribosomal skip peptide. Ultramer primers (Table 1): u-*elo*1-1 and u-*elo*1-2 were used for *elo*1, u-*elo*2-1, and u-*elo*2-2 for *elo*2 and u-*elo*3-1 and u-*elo*3-2 for *elo*3. Parasites were transfected simultaneously with pTREX-Cas9-gRNA plasmids and donor DNA.

**Table 1:**
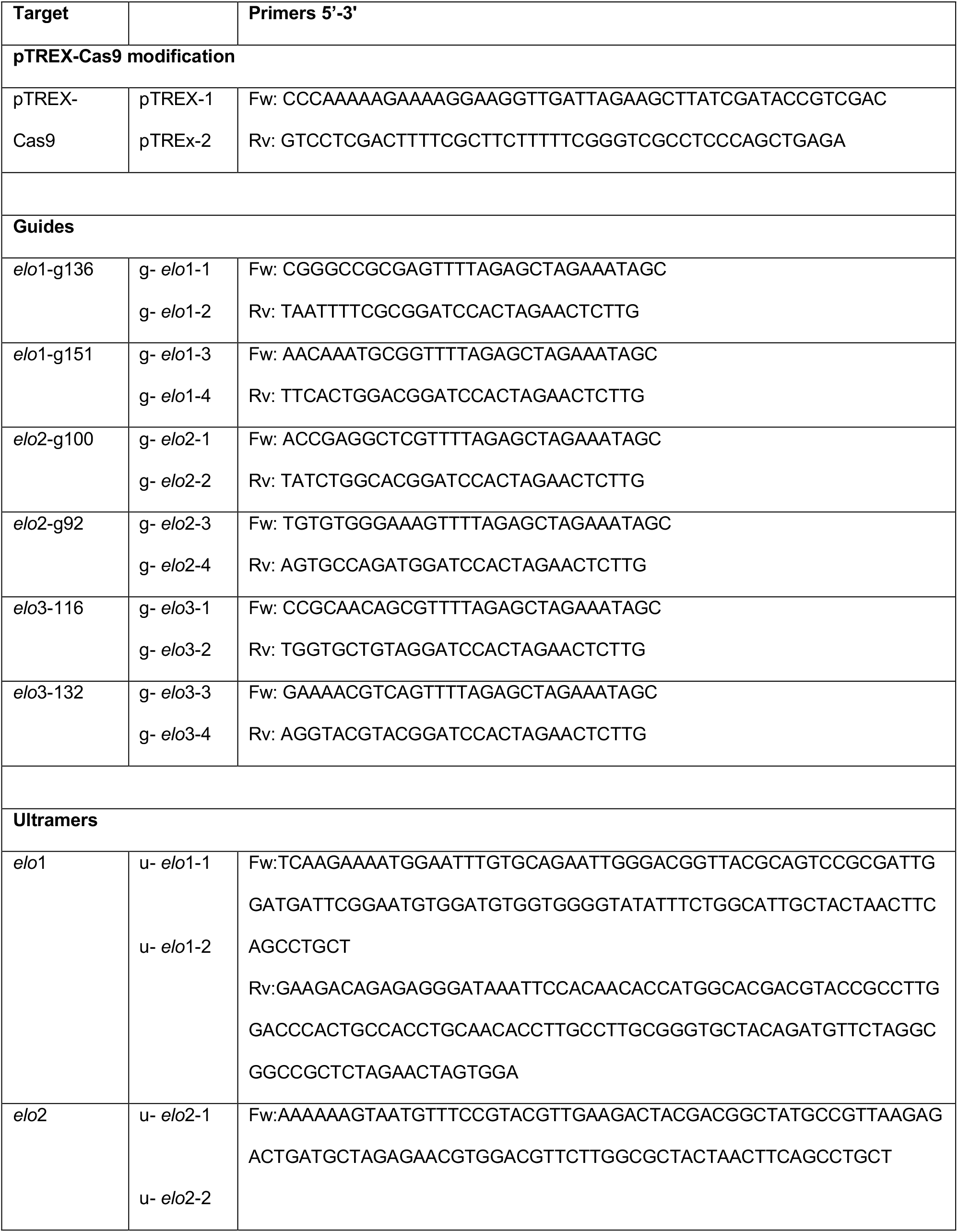
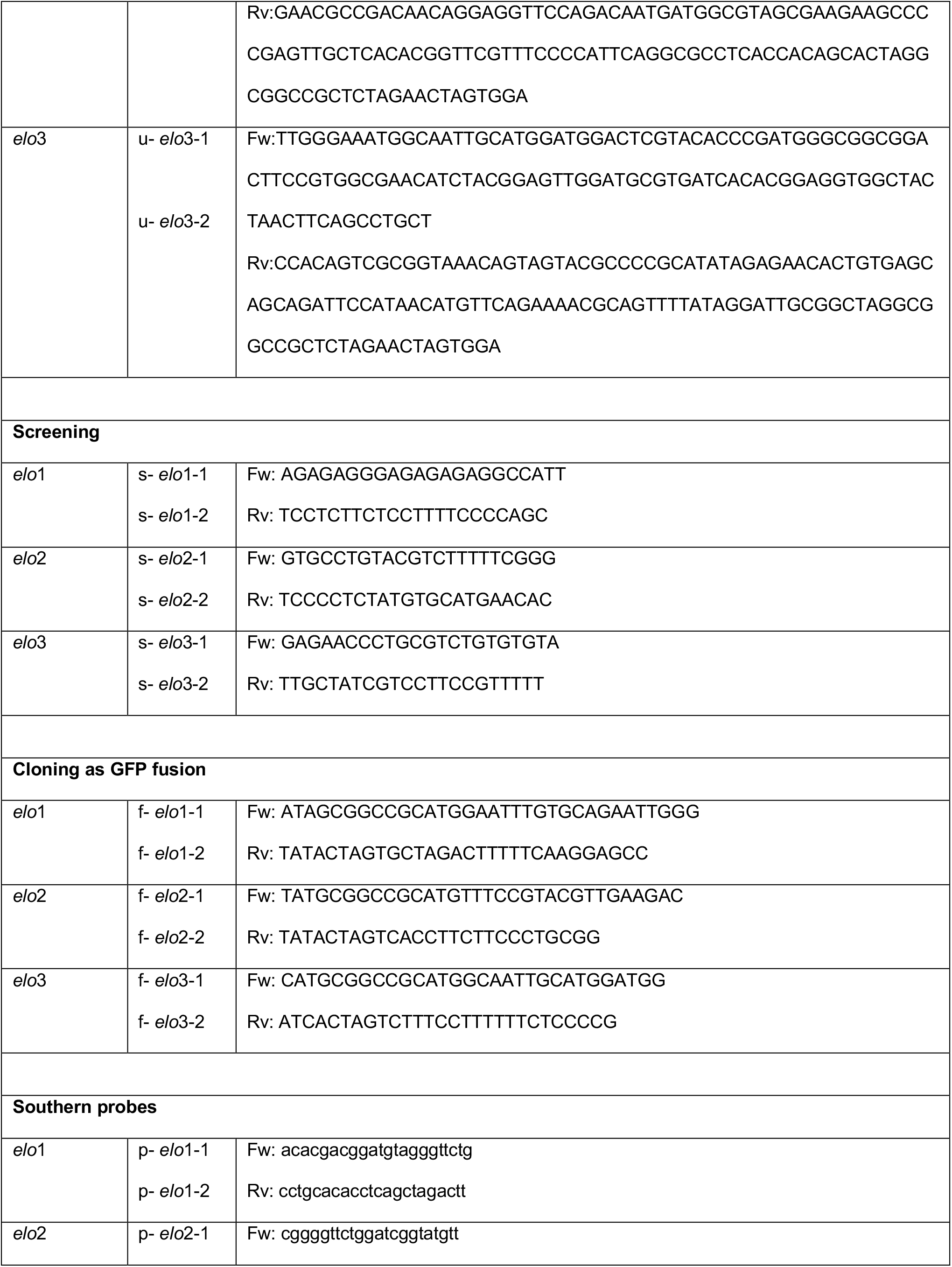
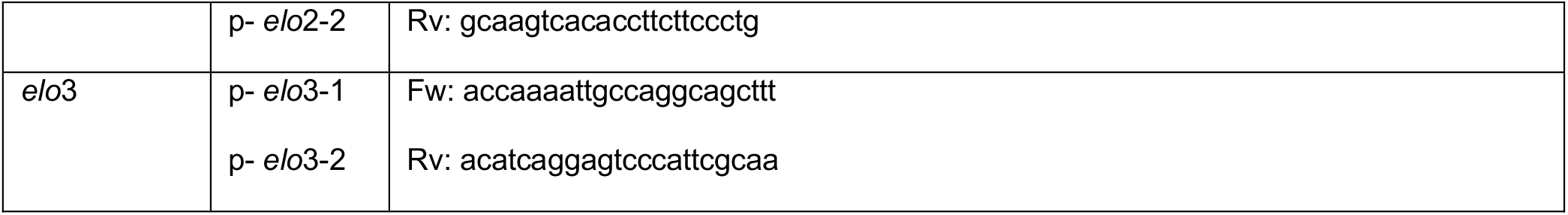
Primers.

### Parasite transfection

Log-phase *T. cruzi* epimastigotes were transfected with plasmid DNA using an Amaxa nucleofector, program U-33 in Tb-BSF ^41^. Transfected parasites were allowed to recover for 24 hours and subsequently cloned in the presence of a drug corresponding to the resistance marker present in the donor DNA. Clones were screened by PCR using the following primers: s-Elo1-1 and s-Elo1-2 for *Elo*1, s-Elo2-1 and s-Elo2-2 for *Elo*2, and s-Elo3-1 and s-Elo3-2 for *Elo*3 (Table 1).

### Southern Blot

Southern blots were accomplished following the specifications of ECL Direct Nucleic Acid Labeling and Detection System (GE Healthcare, UK). Briefly, genomic DNA was digested with PstI and EcoRI (*Elo*1) or DraI and EcoRI (*Elo*2 and *Elo*3), separated on an agarose gel, and blotted into a nitrocellulose membrane. Membranes were incubated with specific probes and developed using ECL reagent. Probes were generated by PCR using the following primers listed in Table 1: p-Elo1-1 and p-Elo1-2 for *Elo*1, p-Elo2-1 and p-Elo2-2 for *Elo*2 and p-Elo3-1 and p-Elo3-2 for *Elo*3.

### Genetic complementation

*Elo* genes were amplified by PCR using the following primers (Table 1): f-Elo1-1 and f-Elo1-2 for *Elo*1, f-Elo2-1, and f-Elo2-2 for *Elo*2, and f-Elo3-1 and f-Elo3-2 *Elo*3. PCR products were cloned into the pTREX-GFP plasmids between the Not1 and Spe1 restriction sites. Log-phase EPIs were transfected with pTREX-GFP-Elo plasmids. Transfected EPIs were allowed to recover for 24 h and, subsequently, cloned in the presence of drug corresponding to the resistance marker present in the plasmid.

### Metabolic tracing and thin-layer chromatography

*T. cruzi* EPIs (1×10^8^) were incubated with 0.3 μCi/mL ^[1-14]^C-labeled fatty acids for 18 h, and total lipids were extracted with chloroform:methanol (2:1, v/v) and dried under N_2_ gas. Modified Folch’s partition, using chloroform:methanol:water (4:2:1.5 v/v/v/) ^42^, was carried out on two-thirds of total lipids, and neutral and polar lipids were resolved by TLC using silica-gel matrix. Fatty acid methyl esters (FAME) were generated from one-third of the remaining total lipids, by incubation with 1 M KOH in methanol for 2 h at 68°C. Folch’s partition was carried out on the generated FAMEs, and fatty acids were resolved by reverse phase (RP)-TLC, using silica-gel 60 RP-18 matrix (Sigma-Aldrich). Mobile phases used were: acetone:methanol:acetic acid:chloroform:water (15:13:12:40:8, v/v/v/v/v) for polar lipids, hexane: diethyl-ether: acetic acid (80:20:3, v/v/v) for neutral lipids, and chloroform:methanol:water (5:15:1, v/v/v) for FA length chain analysis on RP-TLC. The equivalent of 2×10^7^ parasites was spotted per lane. Labeled lipids were detected by phosphor-imaging (Typhoon FLA 7000, GE).

### Protein quantification and lipid extraction

Epimastigotes (2×10^8^) were resuspended in 500 μL ice-cold PBS, and a small aliquot (30 μL) was removed for protein quantification (Pierce BCA Protein Assay Kit; Thermo Fisher, Waltham, Massachusetts). For lipid quantification, EquiSPLASH mix (Avanti Polar Lipids, Alabaster, AL) deuterated standards were spiked into the first extraction solvent mixture (chloroform:methanol:PBS, 1:2:0.8, v/v/v). For every 50 μg of protein, 1 μL EquiSPLASH mix (Avanti Polar Lipids) was added just prior to cell extraction. Parasite pellets were further subjected to three sequential extractions with chloroform:methanol (2:1, v/v), pooled, and dried under N_2_ gas, then subjected to modified Folch’s partition ^42^. Folch’s lower (LP) were separated, pooled, dried under N_2_ gas, and stored at -20°C until analysis. Folch’s LP were redissolved in 100 μL chloroform:methanol (1:2, v/v) for UHPLC-LC-MS/MS analysis.

### Liquid Chromatography-High Resolution Tandem Mass Spectrometry

Ten microliters of each biological sample were run in two technical replicates, separated by reverse phase Kinetex C18 EVO 2.6 μm, 100Å, 150 × 2.1 mm (Phenomenex) column attached to a Dionex Ultimate 3000 UHPLC (Thermo Fisher Scientific). The column was heated to 45°C with a flow rate of 0.5 mL/min throughout the entire run. Equilibrated with 30% solvent A (50% acetonitrile, 10 mM ammonium formate, 0.1% formic acid), 70% solvent B (88% isopropanol, 10% acetonitrile, 2 mM ammonium formate, 0.02% formic acid) and was maintained for 3 min, raised to 43% solvent B over 3 min, 45% over 0.2 min, 65% over 9.8 min, 85% over 6 min, 100% over 2 min where the plateau was maintained for 5 min dropped to 30% solvent B over 0.1 and re-equilibrated for 2.9 min before next sample injection. Data was acquired from eluting metabolites on a Q-Exactive Plus Hybrid Quadrupole Orbitrap Mass Spectrometer equipped with a HESI-II source. Acquisition was performed using Top10 ions in data-dependent MS^2^ Full MS parameters: scan range 250-1800 *m/z*, resolution 70,000, AGC target 1e^6^, max IT 75ms; dd-MS^2^ parameters: resolution 17,500, AGC target 1e^5^, isolation window 1.2 *m/z*, normalized collision energy N(CE) stepped 20, 30, 40, modified ^43^. Samples were run in both positive and negative ion modes before processing.

### Data analysis, statistics, and graphics generation

Positive and negative ion mode raw data files were processed on MS DIAL version 4.16 (RIKEN) ^44^. Processed samples were normalized EquiSPLASH concentration values from the abs. amount in extraction solvent volume then set to pmol/10^6^ cells by manually inputting deuterated standard and alignment ID values. Normalized data were filtered by reference match ^44^. Lipids with multiple ID’s were identified by retention time values and combined. Filtered lipids were utilized to generate heatmaps and principal component analysis (PCA) by uploading to Metaboanalyst 4.0, submitted as concentrations, filtered by the relative standard deviation (RSD), normalized to a wild-type as a reference sample, log-transformed, and auto-scaled as per the workflow of Metaboanalyst ^44^. Principal component analysis (PCA) was generated in addition to generated heatmap specifics including a distance measurement set to Euclidean, clustering average, top 50 PLS-DA VIP. Bar graphs were generated by averaging the raw data values in Prism GraphPad v8.3.1.

### Metabolomics

Epimastigotes (2×10^8^) in exponential growth phase were pelleted by centrifugation at 1,200 x *g* for 5 min and resuspended in 4 mL of methanol at -80°C and incubated for 30 min at - 80°C. Samples were centrifuged at full speed for 5 min, and the supernatant was transferred to a new tube. The extraction procedure was repeated twice and resulting supernatants were combined, and dried under N_2_ gas. Dried samples were resuspended in 20 μL of HPLC water before running on a 5500 QTRAP hybrid triple quadrupole mass spectrometer as described ^45^.

### Cell cycle analysis by flow cytometry

Cell cycle was analyzed as previously reported ^46^. Briefly, exponentially growing *T. cruzi* epimastigotes were fixed on ice in 4% paraformaldehyde/PBS for 20 min. After fixation parasites were centrifuged at 4,000 x *g* for 10 min and the resulting pellet was resuspended in PBS. Immediately prior to acquisition parasites were pelleted at 1,500 x *g* for 10 min and resuspended in a 0.1% Triton X-100/PBS permeabilization solution containing 10 ng/mL DAPI (Sigma-Aldrich) for a minimum of 30 min on ice and analyzed using a MACSQuant VYB flow cytometer. Parasites were identified based on size and DAPI staining. Proliferation modeling based on signal intensity was generated using FlowJo (Tree Star) proliferation software. Greater than 10,000 events in the final EPI gate were acquired for each sample.

### Immunofluorescence microscopy

Parasites were fixed in 1% PFA-PBS for 10 min and spotted onto poly-L-lysine coated slides and allowed to adhere for 30 min. Adhered parasites were washed with PBS and permeabilized with 0.1% Triton X-100-PBS for 10 min, washed again with PBS, and blocked with 3% BSA (Sigma-Aldrich) in PBS for 1 h. Then, parasites were incubated with anti-BIP antibodies in 1% BSA in PBS for 1 h (A generous gift from Dr. J. D. Bangs, U. Buffalo), and washed 3 times with PBS before incubation for 1 h with goat anti-rabbit IgG secondary antibody Alexa fluor 594 (Thermo Fisher Scientific, Waltham, MA), and 100 ng/mL DAPI (Sigma-Aldrich). Finally, parasites were washed with PBS and mounted with ProLong Antifade (Thermo Fisher Scientific). EPIs were analyzed using a Yokogawa CSU-X1 spinning disk confocal system paired with a Nikon Ti-E inverted microscope equipped with an iXon Ultra 888 EMCCD camera. All images were acquired with the 100x objective and image processing was completed using FIJI ^47^.

### Mitochondrial membrane potential

Mitochondrial membrane potential was measured using TMRE Mitochondrial membrane Potential Assay Kit (Abcam, USA), according to the manufacturer’s instructions. Fluorescence intensity was measured by flow cytometry (MACSQuant VYB) for >10,000 events in the final EPI gate were acquired for each sample. Each experiment was carried out in triplicate in at least two independent experiments. Fluorescence intensity was analyzed using FlowJo software (version 10.7.2).

### Superoxide determination

Mitochondrial superoxide determination was accomplished following the specifications of MitoSox Red mitochondrial superoxide indicator (Invitrogen, Waltham, Massachusetts). Briefly, EPI (5×10^6^ cells) in exponential growth were collected and resuspended in 1 mL of LIT media, and 100 μL were settled in 96-well plate. 100 μL of MitoSox staining solution or LIT was added to the samples and incubated 60 min at 27°C. Fluorescence intensity was acquired by flow cytometry. Greater than 10,000 events in the final EPI gate were acquired for each sample. Each experiment was carried out as biological triplicates and at least two experimental rounds. Fluorescence intensity was analyzed with FlowJo 10.7.2.

### Fatty acid oxidation assay

Fatty acid β-oxidation assay was performed, as described ^48^. Briefly, EPI (1×10^8^ cells) in exponential growth were pelleted by centrifugation at 1,200 x *g* for 5 min and resuspended in 1 mL of PBS with 10 μCi of ^3^H-palmitate and 0.25 mM carnitine, or without supplementation. 100 μL samples were collected after 4 h of incubation and centrifuged at 3,000 x *g* for 5 min. The supernatant was transferred to a new tube and 100 μL of 10% TCA was added and incubated for 15 min. The supernatant was mixed with 5% TCA and 10% BSA, incubated at RT for 15 min, and extracted with chloroform:methanol (2:1, v/v) and 300 μL of a 2 M potassium chloride : 2 M hydrochloride mix was added. The supernatant was mixed with 5 mL of Ultima Gold XR (Perkin Elmer, Waltham, Massachusetts). The count-per-minute was determined for 5 min for each sample. The count-per-minute (cpm) with a ^3^H-labeled medium was used as background to correct all samples. Final results were obtained after normalization using the protein concentration.

### Western Blot

EPI pellets were washed twice with PBS and resuspended in lysis buffer (0.5% Nonidet P-40, 500 mM NaCl, 5 mM EDTA, 1 mM DTT, 50 mM Tris-Base, 0.4% SDS, pH 7.4) at 1×10^6^ parasites/μL. Samples were sonicated using 3 pulses of 30 seconds at 100% amplitude (Q700 sonicator, QSONICA, Newton, Connecticut), with 15 sec breaks between to rest the tubes on the ice. Samples were centrifuged at 16,000 x *g* for 20 minutes and supernatant were collected. Equivalent to 1×10^6^ parasites were resolved by electrophoresis on a Mini-Protean TGX™ precast gel (Bio-rad, California). Lipoylated proteins were detected with rabbit anti-lipoic acid antibody (ab 58724, abcam, Waltham, Massachusetts). Bound antibodies were detected with horseradish peroxidase-linked anti-rabbit IgG (GE Healthcare, UK), and ECL Prime Western Blotting Detection Kit (GE Healthcare, UK). Loading control was performed with mouse anti α-Tubulin (Sigma-Aldrich), and bounded antibodies were detected with horseradish peroxidase-linked anti-mouse IgG (GE Healthcare, UK). Quantification was performed after normalization of lipoic acid signal to tubulin, using FIJI ^47^.

## Acknowledgements

We wish to thank Jay Bangs (U. Buffalo) for the anti-BiP antibody and Madalyn Won for constructive comments. This work was funded by the National Institutes of Health grant R01 AI114622 awarded to B.A.B. and American Heart Association Founders Affiliate Postdoctoral fellowship ID:825688 awarded to L.P.

