## Supplemental Figures for "Fatty acid elongases 1-3 have distinct roles in mitochondrial function, growth and lipid homeostasis in *Trypanosoma cruzi*"

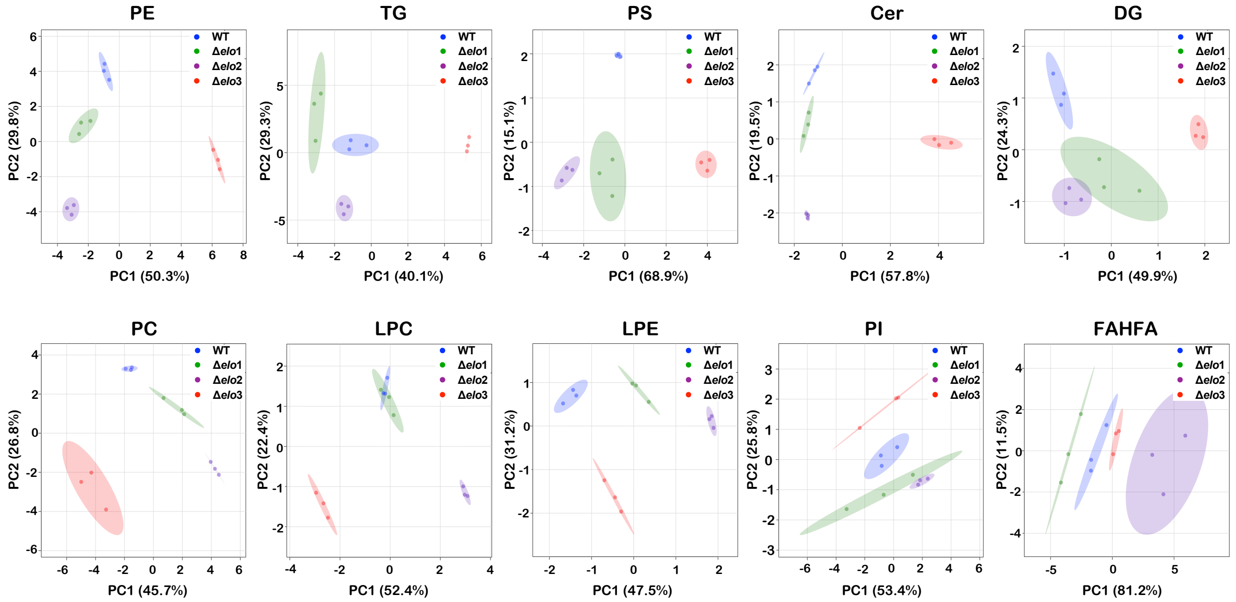


**Figure S1. PCA plots of WT and Δ*elo* mutants by lipid class.** Principal component analysis of lipid species plotted for the indicated lipid subclasses: PE, phosphatidylethanolamine; TG, triacylglycerol; PS, phosphatidylserine; Cer, ceramides; DG, diacylglycerol; PC, phosphatidylcholine; LPC, *lyso*-phosphatidylcholine; LPE, *lyso*-phosphatidylethanolamine; PI, phosphatidylinositol; FAHFA, fatty acyl esters of hydroxy fatty acids. The first two principal components are plotted (PC1 and PC2) with the proportion of variance for each component in parenthesis and the 95% confidence interval indicated in shaded areas.

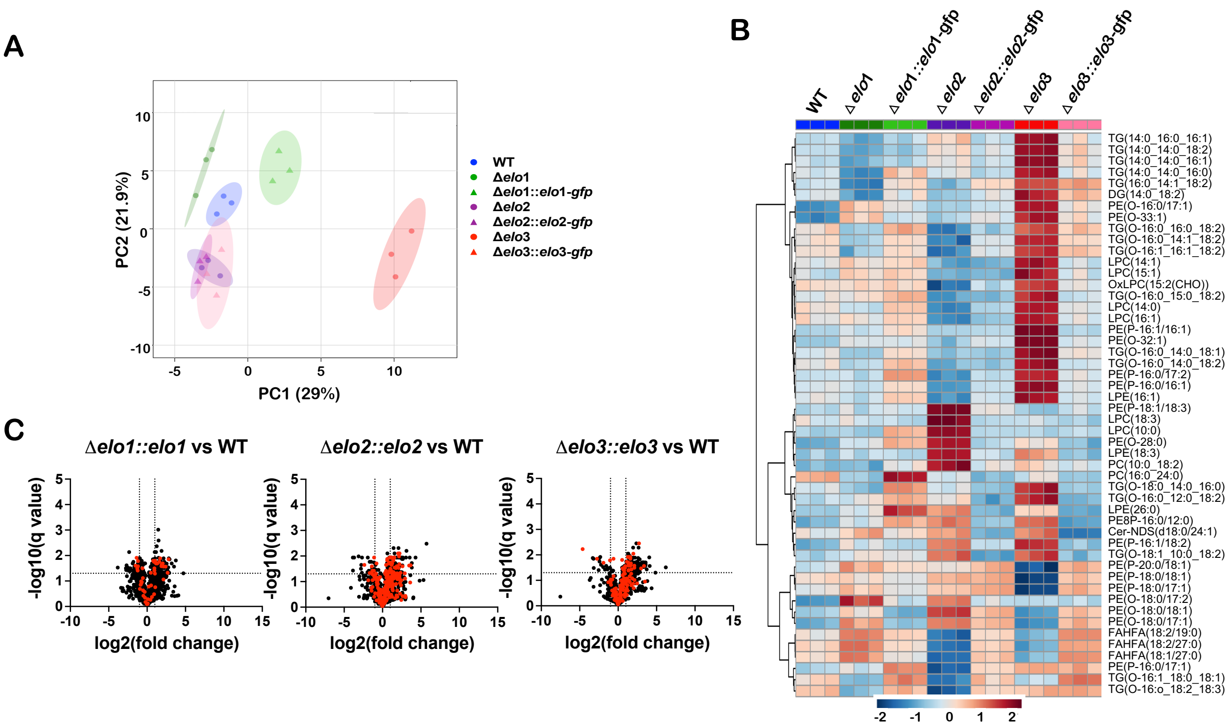


**Figure S2. Lipidomic characterization of *T. cruzi* Δ*elo* mutants and genetically complemented lines. A.** Principal component analysis (PCA) of total lipidome data plotted for WT, each Δ*elo* mutant and corresponding genetically-complemented mutant *T. cruzi* EPI. The first two principal components are shown (PC1 and PC2) with the proportion of variance for each component in parenthesis. Biological replicates for each experimental group are represented and the 95% confidence interval indicated in shaded circle. **B.** Heat map displaying the 50 most significantly up or down regulated lipids when comparing Δ*elo* mutants and genetically-complemented *T. cruzi* EPI with WT parasites. **C.** Volcano plots generated based on fold-change (x-axis) and adjusted p-value (q-value; y-axis) in pairwise comparisons between WT and genetically-complemented lines: Δ*elo1::elo1-gfp,* Δ*elo2::elo2-gfp,* and Δ*elo3::elo3-gfp T. cruzi* EPI. Horizontal line represent significant q value (q≤0.05), vertical lines indicate ≥2-fold change-≤0.5 cut-off. Lipids found to be significantly altered in abundance (≥2-fold change or fold change-≤0.5; adjusted p-value ≤0.05) in the corresponding Δ*elo* mutant (Fig. 2C) are highlighted (red circles) for each comparison. PE, phosphatidylethanolamine; TG, triacylglycerol; PS, phosphatidylserine; Cer, ceramides; DG, diacylglycerol; PC, phosphatidylcholine; LPC, *lyso*-phosphatidylcholine; LPE, *lyso*-phosphatidylethanolamine; PI, phosphatidylinositol; FAHFA, fatty acyl esters of hydroxy fatty acids.

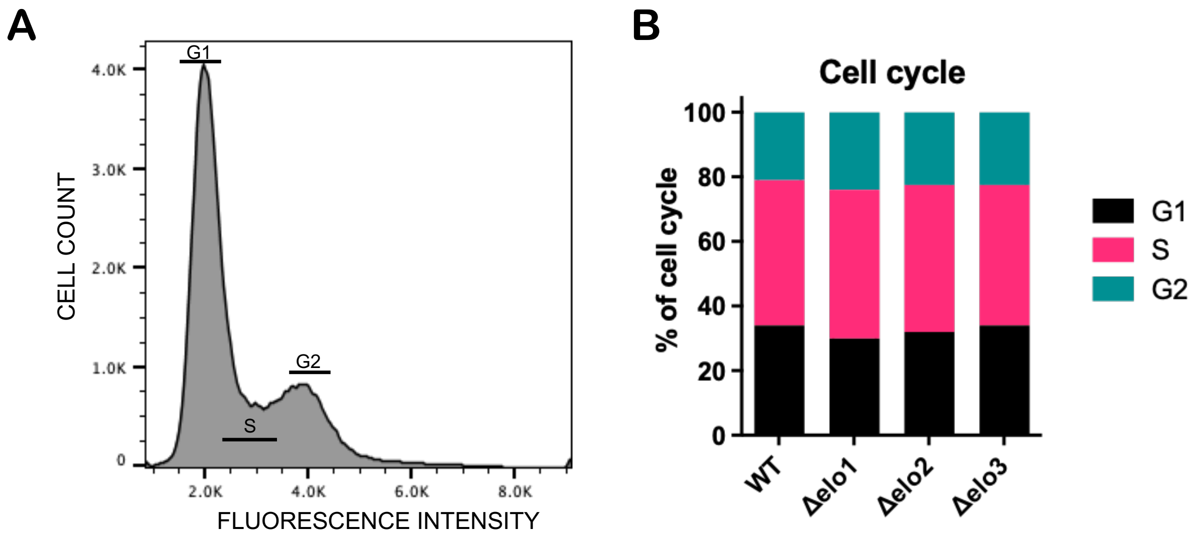


**Figure S3. Cell cycle analysis of *T. cruzi* EPI.** Flow cytometry was used to assess the cycle-cycle status of log-phase *T. cruzi* EPI populations. **A.** Representative histogram obtained following flow cytometry of DAPI-stained *T. cruzi* EPI showing the number of cells in each cell-cycle stage, inferred by measuring the DNA content (DAPI staining). **B.** Percentage of parasites in the G1, S and G2 stages of the cell- cycle.

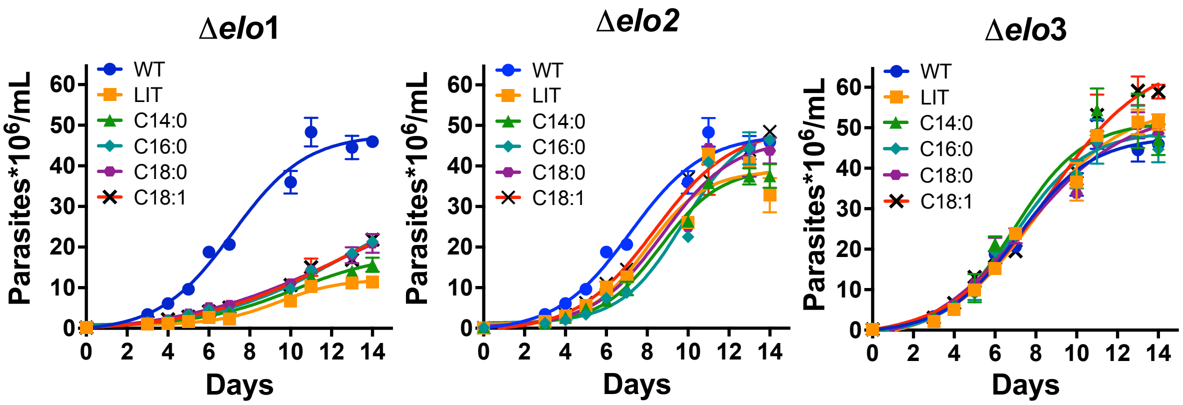


**Figure S4. LCFA supplementation has minimal impact on *T. cruzi* epimastigote growth.** The growth of Δ*elo* mutant *T. cruzi* EPI lines in LIT medium containing 10% FBS, with or without supplementation with 35 μM of indicated fatty acid were compared to WT EPI in LIT medium + 10% FBS and shown on separate graphs. The mean and s.d. calculated for parasite density (parasites x $10^6$/ml medium; y-axis) for 3 biological replicates against time (days; x-axis) is plotted. LIT: LIT + 10% FBS; no FA added. C14:0; myristic acid, C16:0; palmitic acid, C18:0; stearic acid, C18:1, oleic acid.

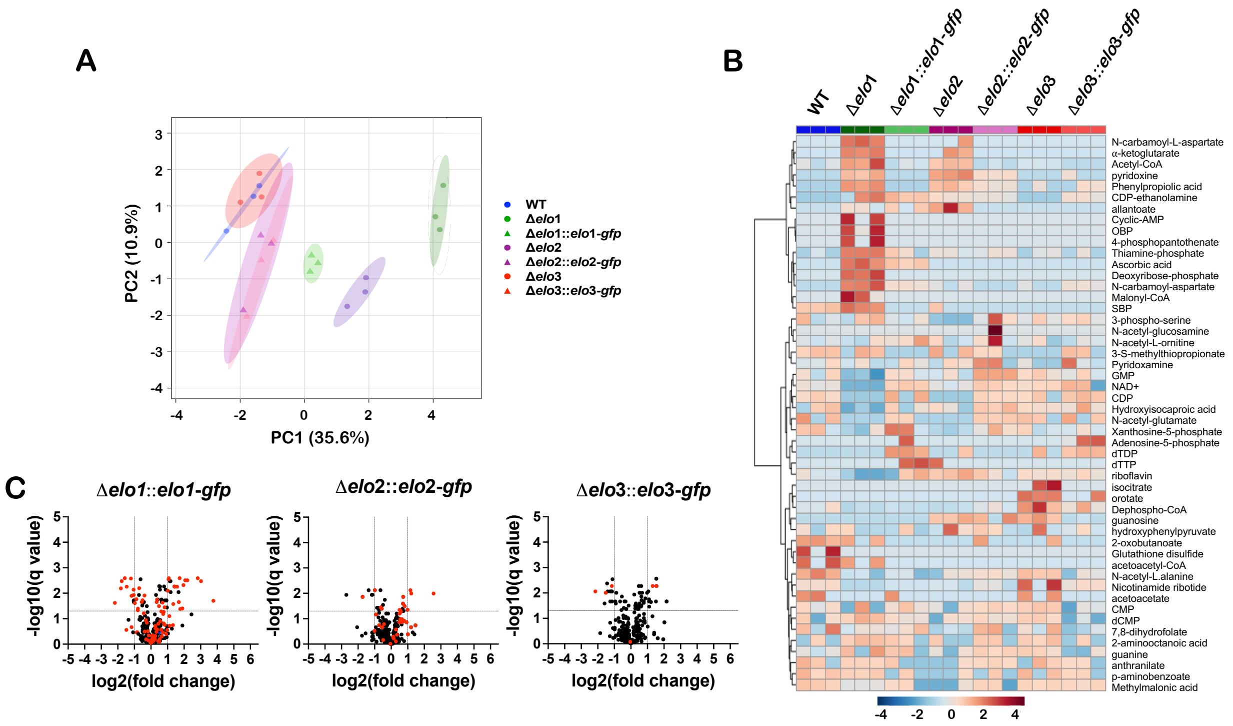


**Figure S5. Metabolomic analysis of WT, Δ*elo* mutants and genetically complemented strains**. **A.** Principal component analysis (PCA) plot of metabolite data plotted for WT, each Δ*elo* mutant and corresponding genetically-complemented mutant *T. cruzi* lines. The first two principal components are shown (PC1 and PC2) with the proportion of variance for each component in parenthesis. Biological replicates for each experimental group are represented and the 95% confidence interval indicated in shaded circle. **B.** Heat map displaying the 50 metabolites displaying the greatest change in parasite groups as compared to WT *T. cruzi* EPI. **C.** Volcano plots generated based on fold-change (x-axis) and adjusted p-value (q-value; y-axis) in pairwise comparisons between WT and genetically-complemented lines: Δ*elo1::elo1-gfp,* Δ*elo2::elo2-gfp,* and Δ*elo3::elo3-gfp T. cruzi* EPI. Horizontal lines represent significant q value (q≤0.05), vertical lines indicate ≥2-fold change-≤0.5 cut-off. Metabolites found to significantly altered in abundance (≥2-fold change or fold change-≤0.5; adjusted p-value ≤0.05) in each corresponding Δ*elo* mutant (Fig. 5C) is highlighted (red circles) for each comparison.
