## Supplemental Table 3 for "Fatty acid elongases 1-3 have distinct roles in mitochondrial function, growth and lipid homeostasis in *Trypanosoma cruzi*"

| Metabolite name | formula | PubChem Identifier (KEGG/HMDB) | | |
|---|---|---|---|---|
| | | | elo1 | elo2 |
| 1-Methyl-Histidine | C7H11N3O2 | C01152 | 0.436562927 | 0.434275839 |
| 1-Methyladenosine | C11H15N5O4 | C02494 | 7.016904768 | 1.382670149 |
| 2-deoxyglucose-6-phosphate | C6H13O8P | C06369 | 5.991703036 | 3.846127086 |
| 3-hydroxybutyrate | C4H8O3 | HMDB0000011 | 2.332154448 | 1.266342125 |
| 3-phosphoglycerate | C3H7O7P | C00197 | 5.929541287 | 5.169202869 |
| 6-phospho-D-gluconate | C6H13O10P | C00345 | 2.433199676 | 1.424925455 |
| a-ketoglutarate | C5H6O5 | C00026 | 4.007989909 | 3.127072107 |
| acetyl-CoA | C23H38N7O17P3S | C00024 | 5.529191393 | 3.654549016 |
| allantoate | C4H8N4O4 | C00499 | 4.706743383 | 14.08243856 |
| Aminoadipic acid | C6H11NO4 | C00956 | 3.984671475 | 2.602584415 |
| Ascorbic acid | C6H8O6 | C00072 | 3.902036439 | 0.931248538 |
| asparagine | C4H8N2O3 | C00152 | 2.2841834 | 1.269013896 |
| betaine | C5H11NO2 | C00719 | 0.444115774 | 0.563002906 |
| Carbamoyl phosphate | CH4NO5P | C00169 | 2.355173647 | 1.275208684 |
| CDP-choline | C14H27N4O11P2 | C00307 | 0.423730328 | 0.956351427 |
| CDP-ethanolamine | C11H20N4O11P2 | C00570 | 4.132408372 | 1.855261235 |
| cholesterol | C27H46O | C00187 | 3.502673033 | 2.151611341 |
| creatine | C4H9N3O2 | C00300 | 0.352254896 | 0.554838013 |
| cyclic-AMP | C10H12N5O6P | C00575 | 2.189817449 | 1.313220331 |
| cytidine | C9H13N3O5 | C00475 | 4.411401453 | 1.207817774 |
| D-erythrose-4-phosphate | C4H9O7P | C00279 | 2.455960718 | 2.216060496 |
| D-glucosamine-6-phosphate | C6H14NO8P | C00352 | 5.163976003 | 3.024136323 |
| D-glyceraldehdye-3-phosphate | C3H7O6P | C00118 | 1.075882946 | 0.670270228 |
| D-sedoheptulose-1-7-phosphate | C7H15O10P | C05382 | 3.751331682 | 2.217793863 |
| dATP | C10H16N5O12P3 | C00131 | 0.652110137 | 2.828431856 |
| deoxyadenosine | C10H13N5O3 | C00559 | 1.225780156 | 4.647497959 |
| deoxyinosine | C10H12N4O4 | C05512 | 2.410276967 | 1.526476256 |
| dephospho-CoA | C21H35N7O13P2S | C00882 | 0.306394833 | 0.280161513 |
| dihydroorotate | C5H6N2O4 | C00337 | 1.70406926 | 7.588999596 |
| dimethylglycine | C4H9NO2 | C01026 | 0.660212594 | 0.415476323 |
| dTTP | C10H17N2O14P3 | C00459 | 2.197709806 | 3.697939877 |
| fructose-6-phosphate | C6H13O9P | C05345 | 9.824848198 | 6.504781376 |
| GDP | C10H15N5O11P2 | C00035 | 0.484255601 | 0.535192496 |
| glucono-δ-lactone | C6H10O6 | C00198 | 3.937837051 | 0.798162627 |
| glucose-1-phosphate | C6H13O9P | C00103 | 3.072317491 | 1.441043864 |
| glucose-6-phosphate | C6H13O9P | C00668 | 7.775386731 | 5.782910908 |
| glutamine | C5H10N2O3 | C00064 | 5.841101613 | 3.756005961 |
| glutamine | C5H10N2O3 | C00064 | 5.841101613 | 3.756005961 |
| glutathione disulfide | C20H32N6O12S2 | C00127 | 0.300791309 | 0.763792635 |
| glycerate | C3H6O4 | C00258 | 2.221762353 | 1.292969084 |
| GTP | C10H16N5O13P3 | C00044 | 0.46731283 | 1.369084461 |
| guanine | C5H5N5O | C00242 | 2.054468719 | 1.180073786 |
| hexose-phosphate | C6H13O9P | C05345 | 2.215780096 | 2.033398686 |
| HMG-CoA | C27H44N7O20P3S | C00356 | 2.770438741 | 1.673599172 |
| homocysteic acid | C4H9NO5S | C16511 | 0.854602794 | 0.499836996 |
| hydroxyproline | C5H9NO3 | C01157 | 0.473954886 | 0.730875065 |
| IMP | C10H13N4O8P | C00130 | 0.342213773 | 0.808868655 |
| Indole-3-carboxylic acid | C9H7NO2 | HMDB03320 | 4.082883399 | 2.716330184 |
| malonyl-CoA | C24H38N7O19P3S | C00083 | 52.08464324 | 6.649047273 |
| N-acetyl-glucosamine | C8H15NO6 | C00140 | 4.463446599 | 3.777500205 |
| N-acetyl-glutamine | C7H12N2O4 | HMDB06029 | 0.395508498 | 0.618518967 |
| N-Acetyl-L-alanine | C5H9NO3 | C01073 | 0.251326601 | 0.417937949 |
| N-acetyl-L-aspartic acid | C6H9NO5 | C01042 | 3.055731595 | 1.938111656 |
| N-carbamoyl-L-aspartate | C5H8N2O5 | C00438 | 9.438928761 | 2.364805867 |
| NAD+ | C21H27N7O14P2 | C00003 | 0.23132809 | 0.905236195 |
| NADH | C21H29N7O14P2 | C00004 | 0.183730878 | 0.86575556 |
| NADP+ | C21H28N7O17P3 | C00006 | 2.646459478 | 1.338112356 |
| nicotinamide | C6H6N2O | C00153 | 0.387715779 | 3.699675724 |
| Nicotinamide ribotide | C11H15N2O8P | C00455 | 0.086143603 | 0.365579462 |

| | | | | |
|---|---|---|---|---|
| O-acetyl-L-serine | C5H9NO4 | C00979 | 2.116297425 | 1.761831576 |
| octulose-1,8-bisphosphate (OBP) | C5H11O8P | | 5.231862869 | 1.206705128 |
| Phenylpropiolic acid | C9H6O2 | HMDB00563 | 4.431922276 | 3.268968689 |
| phenylpyruvate | C9H8O3 | C00166 | 5.025054885 | 1.28624446 |
| phosphoenolpyruvate | C3H5O6P | C00074 | 5.846157981 | 6.498159385 |
| Phosphorylcholine | C5H15NO4P | C00588 | 2.235509023 | 3.252041202 |
| propionyl-CoA | C24H40N7O17P3S | C00100 | 2.342520314 | 1.104362795 |
| pyridoxine | C8H11NO3 | C00314 | 9.517525151 | 12.2052316 |
| riboflavin | C17H20N4O6 | C00255 | 0.494346872 | 1.66876672 |
| S-adenosyl-L-homocysteine | C14H20N6O5S | C00021 | 0.227610272 | 0.989862634 |
| S-ribosyl-L-homocysteine | C9H17NO6S | C03539 | 0.376460469 | 0.59338057 |
| shikimate-3-phosphate | C7H11O8P | C03175 | 0.463026795 | 0.706548981 |
| spermine | C10H26N4 | C00750 | 3.068554535 | 1.443112279 |
| Thiamine pyrophosphate | C12H19N4O7P2S | C00068 | 4.668230861 | 2.660794606 |
| thiamine-phosphate | C12H17N4O4PS | C01081 | 4.736799364 | 1.559227404 |
| threonine | C4H9NO3 | C00188 | 6.206739958 | 3.430588393 |
| UDP-D-glucose | C15H24N2O17P2 | C00029 | 15.85665196 | 9.037686349 |
| UDP-N-acetyl-glucosamine | C17H27N3O17P2 | C00043 | 18.45434481 | 12.07427948 |
| UMP | C9H13N2O9P | C00105 | 0.485079527 | 0.617343589 |
| uracil | C4H4N2O2 | C00106 | 1.639038873 | 2.287654875 |
| Uric acid | C5H4N4O3 | C00366 | 2.320830103 | 1.411233932 |

| Mean Fold change to WT | | | | q values | | | | |
|---|---|---|---|---|---|---|---|---|
| elo3 | elo1::elo1-gfp | elo2::elo2-gfp | elo3:elo3-gfp | elo1 | elo2 | elo3 | elo1::elo1-gfp | elo2::elo2-gpf |
| 0.375170713 | 0.284016097 | 0.301829675 | 0.225566529 | 0.00674 | 0.011414 | 0.027816 | 0.006492 | 0.013729 |
| 0.94152799 | 1.426061528 | 0.52612471 | 0.809362781 | 0.033257 | 0.650051 | 0.977846 | 0.806768 | 0.216794 |
| 2.102757408 | 1.538514067 | 2.239658064 | 2.919879327 | 0.000182 | 0.007922 | 0.011606 | 0.051898 | 0.007573 |
| 1.012335929 | 1.271515412 | 1.153118902 | 0.880152529 | 0.000376 | 0.026752 | 0.934728 | 0.022348 | 0.120786 |
| 1.07620828 | 2.979156065 | 1.220767856 | 1.290139649 | 0.000929 | 0.001593 | 0.665514 | 0.005562 | 0.266186 |
| 1.375727065 | 0.976340855 | 0.942872049 | 1.071575644 | 0.015875 | 0.325493 | 0.434748 | 0.865492 | 0.892761 |
| 1.769848463 | 1.174966484 | 1.896330797 | 2.153444823 | 0.002085 | 0.01925 | 0.288404 | 0.630489 | 0.138272 |
| 0.715030001 | 0.814803145 | 0.498111104 | 0.505317862 | 0.019163 | 0.008163 | 0.568565 | 0.648819 | 0.306424 |
| 1.641944661 | 7.086788687 | 1.326417079 | 1.055723513 | 0.00038 | 0.069715 | 0.184031 | 0.002565 | 0.196381 |
| 0.724808077 | 0.806425229 | 1.467711081 | 1.562346437 | 0.000644 | 0.002261 | 0.092145 | 0.113253 | 0.04707 |
| 1.162298333 | 2.017128763 | 1.136794651 | 1.392756499 | 0.003618 | 0.721332 | 0.476311 | 0.008046 | 0.286391 |
| 0.975704923 | 0.703532845 | 0.811586885 | 0.761957099 | 0.000409 | 0.099136 | 0.879227 | 0.019028 | 0.062609 |
| 0.868264012 | 0.680942591 | 0.895509526 | 0.808373327 | 0.002249 | 0.008163 | 0.247284 | 0.022348 | 0.367057 |
| 1.0275991 | 1.069020055 | 1.259704246 | 1.256915696 | 0.000862 | 0.030938 | 0.809589 | 0.447237 | 0.062609 |
| 1.077207167 | 1.069455317 | 1.229130012 | 0.911073119 | 0.000578 | 0.099136 | 0.456486 | 0.557323 | 0.264976 |
| 1.148068294 | 2.657181469 | 1.494761529 | 1.719143827 | 0.001617 | 0.007922 | 0.510126 | 0.005444 | 0.038622 |
| 0.396414253 | 2.50946752 | 0.777359467 | 0.559606195 | 0.006929 | 0.110776 | 0.394833 | 0.177903 | 0.751176 |
| 0.786337152 | 0.43334083 | 0.638527808 | 0.462882372 | 0.000383 | 0.000959 | 0.011606 | 0.00266 | 0.007573 |
| 1.264756423 | 1.11259653 | 0.959760925 | 1.334430711 | 0.023824 | 0.400015 | 0.661042 | 0.738609 | 0.926957 |
| 1.464168681 | 1.578220173 | 1.068767525 | 0.63736766 | 0.009276 | 0.61752 | 0.40155 | 0.490768 | 0.910197 |
| 1.107880831 | 1.617012493 | 1.108751916 | 1.21495458 | 0.002585 | 0.019026 | 0.634182 | 0.023389 | 0.569358 |
| 1.267147617 | 3.413576172 | 2.300774544 | 1.436083667 | 0.004861 | 0.02576 | 0.662875 | 0.041419 | 0.216168 |
| 0.478676976 | 0.648401253 | 0.388879334 | 0.450675715 | 0.164872 | 0.001593 | 0.011606 | 0.002831 | 0.007573 |
| 1.17599055 | 1.480345534 | 1.571649063 | 1.794083423 | 0.004238 | 0.025368 | 0.510126 | 0.125779 | 0.102947 |
| 1.243886195 | 2.701105458 | 1.653950935 | 1.715214469 | 0.019392 | 0.008662 | 0.228028 | 0.005562 | 0.013729 |
| 0.801153418 | 1.900481454 | 0.712484195 | 0.882024479 | 0.513664 | 0.00818 | 0.796126 | 0.265326 | 0.691945 |
| 1.484661633 | 1.078269377 | 1.106299214 | 1.527001638 | 0.004612 | 0.073199 | 0.162647 | 0.602416 | 0.507857 |
| 1.919743544 | 0.221196995 | 0.605546718 | 2.316826483 | 0.021678 | 0.029522 | 0.156487 | 0.024105 | 0.182012 |
| 0.940398548 | 1.656872288 | 2.011441447 | 1.795226089 | 0.35006 | 0.003583 | 0.82812 | 0.578712 | 0.411599 |
| 0.903186173 | 0.509949717 | 0.527797463 | 0.533181719 | 0.039659 | 0.012229 | 0.575729 | 0.024486 | 0.068221 |
| 1.713737156 | 4.590900957 | 1.75375384 | 2.461604594 | 0.011768 | 0.032088 | 0.075219 | 0.003111 | 0.032117 |
| 0.808196253 | 4.197694674 | 1.415651795 | 1.409677345 | 0.000644 | 0.000959 | 0.327394 | 0.003111 | 0.044975 |
| 0.659369858 | 0.91135697 | 0.845036723 | 1.112194215 | 0.003618 | 0.017588 | 0.092145 | 0.458319 | 0.138272 |
| 1.085442523 | 1.011614843 | 0.737043884 | 0.6375243 | 0.000138 | 0.064225 | 0.733378 | 0.847828 | 0.114281 |
| 1.264644593 | 1.707495009 | 1.088184759 | 1.310235685 | 0.005014 | 0.049305 | 0.327394 | 0.053852 | 0.677097 |
| 0.789953046 | 3.900010935 | 1.404245259 | 1.185431857 | 0.002085 | 0.001533 | 0.327097 | 0.004825 | 0.138272 |
| 2.00864194 | 1.401367382 | 2.314266276 | 2.490686092 | 0.003618 | 0.001593 | 0.018287 | 0.032226 | 0.010155 |
| 2.00864194 | 1.401367382 | 2.314266276 | 2.490686092 | 0.003618 | 0.001593 | 0.018287 | 0.032226 | 0.010155 |
| 0.483063602 | 0.485982045 | 0.498749683 | 0.348040852 | 0.002261 | 0.070394 | 0.032395 | 0.011689 | 0.027138 |
| 0.979249463 | 1.197907555 | 1.184750404 | 1.374907154 | 0.00593 | 0.379474 | 0.977846 | 0.474819 | 0.706441 |
| 0.811635835 | 1.29186526 | 0.983229795 | 0.979317597 | 0.04224 | 0.125767 | 0.463621 | 0.265326 | 0.944558 |
| 1.546307638 | 1.667423281 | 1.587444949 | 1.32954862 | 0.01045 | 0.43913 | 0.184031 | 0.065127 | 0.088592 |
| 1.205700869 | 2.084276731 | 1.087155119 | 1.205687143 | 0.000832 | 0.00214 | 0.526058 | 0.002831 | 0.509781 |
| 1.411068696 | 1.451569079 | 0.671075358 | 1.141503705 | 0.047704 | 0.411958 | 0.687084 | 0.630489 | 0.73052 |
| 1.041247874 | 0.57891222 | 0.684064441 | 0.595845412 | 0.243512 | 0.037686 | 0.89111 | 0.040335 | 0.216794 |
| 0.948795061 | 0.763548762 | 0.886815492 | 0.743922255 | 0.001617 | 0.043718 | 0.570229 | 0.012318 | 0.138272 |
| 0.965206569 | 1.019658109 | 1.192516511 | 0.861373456 | 0.020927 | 0.387687 | 0.924809 | 0.860561 | 0.468293 |
| 0.738977781 | 0.841156209 | 1.640557401 | 1.689959351 | 0.000138 | 0.00118 | 0.12803 | 0.144987 | 0.025568 |
| 0.96814465 | 1.893497742 | 0.970116397 | 1.173575592 | 0.002124 | 0.022802 | >0,999999 | 0.403589 | 0.977289 |
| 1.244342053 | 1.786331858 | 1.273466616 | 0.869332473 | 0.013351 | 0.011414 | 0.673461 | 0.176049 | 0.427639 |
| 0.726399078 | 0.769735244 | 0.606007466 | 0.608699291 | 0.017043 | 0.042794 | 0.217444 | 0.321746 | 0.34785 |
| 0.645752923 | 0.334483089 | 0.505393037 | 0.368540645 | 0.000409 | 0.001593 | 0.017904 | 0.002565 | 0.007573 |
| 2.488641551 | 1.647638018 | 1.097550719 | 0.987052298 | 0.009497 | 0.131742 | 0.296869 | 0.342687 | 0.892761 |
| 1.757197899 | 3.313954077 | 1.467272685 | 2.860125247 | 0.002085 | 0.007922 | 0.213732 | 0.002565 | 0.102947 |
| 1.194033335 | 1.292132072 | 1.260430809 | 1.437204928 | 0.000637 | 0.281686 | 0.147711 | 0.01779 | 0.083314 |
| 1.137242822 | 1.256033468 | 0.969249534 | 1.728882666 | 0.005814 | 0.450826 | 0.510126 | 0.329925 | 0.970363 |
| 1.393819192 | 1.527891887 | 1.17011663 | 1.091648382 | 0.010046 | 0.272252 | 0.327097 | 0.273106 | 0.614894 |
| 1.368825785 | 0.550779017 | 1.217490073 | 0.790233882 | 0.131339 | 0.009292 | 0.533832 | 0.342687 | 0.782401 |
| 2.914832182 | 0.295021416 | 0.672469559 | 0.973499813 | 0.000644 | 0.004277 | 0.013962 | 0.003365 | 0.040477 |

| | | | | | | | | |
|---|---|---|---|---|---|---|---|---|
| 1.085599591 | 0.996112829 | 0.9437893 | 0.854529043 | 0.047704 | 0.325991 | 0.924809 | 0.904101 | 0.848001 |
| 0.56400611 | 1.046871326 | 0.456151879 | 0.41838609 | 0.000645 | 0.151915 | 0.092145 | 0.64834 | 0.01144 |
| 1.572502817 | 1.362155802 | 1.981511684 | 2.140915271 | 0.000672 | 0.008362 | 0.247284 | 0.157975 | 0.044608 |
| 1.514308195 | 1.352322551 | 1.06829751 | 1.825703423 | 0.021724 | 0.736944 | 0.774711 | 0.750646 | 0.970363 |
| 0.993856635 | 3.200421243 | 1.110551943 | 1.273430173 | 0.00537 | 0.004277 | 0.980019 | 0.022348 | 0.601893 |
| 1.176401625 | 0.73982318 | 0.741182029 | 0.675033807 | 0.003131 | 0.000959 | 0.314638 | 0.061387 | 0.164126 |
| 0.480220578 | 2.164625445 | 0.65182271 | 1.254839425 | 0.009045 | 0.701802 | 0.395531 | 0.078854 | 0.375886 |
| 1.729344702 | 3.731831879 | 5.906415654 | 4.466173477 | 0.002511 | 0.002261 | 0.327394 | 0.039677 | 0.010155 |
| 1.287357082 | 1.769694668 | 1.186912816 | 1.257563841 | 0.021306 | 0.037999 | 0.247284 | 0.035561 | 0.690929 |
| 0.815692043 | 0.877852727 | 0.822475223 | 1.254343465 | 0.001739 | 0.842626 | 0.247284 | 0.311565 | 0.19256 |
| 0.892823569 | 0.577879008 | 0.992452741 | 0.916655594 | 0.014026 | 0.078333 | 0.620652 | 0.04658 | 0.970363 |
| 1.260826682 | 0.901695064 | 1.1737779 | 1.030595953 | 0.045988 | 0.298018 | 0.452139 | 0.709214 | 0.807701 |
| 0.53870745 | 2.182982335 | 1.395481504 | 0.7299498 | 0.017758 | 0.627775 | 0.582337 | 0.19008 | 0.782401 |
| 1.232115054 | 2.115298379 | 1.507141725 | 1.607198143 | 0.000138 | 0.004277 | 0.247823 | 0.002565 | 0.052614 |
| 1.258715025 | 1.982767316 | 1.674183135 | 1.775449969 | 0.000138 | 0.168949 | 0.510126 | 0.280513 | 0.138272 |
| 0.523361521 | 0.611649801 | 1.415174567 | 1.546434791 | 0.000665 | 0.001593 | 0.011606 | 0.006333 | 0.062609 |
| 1.151430711 | 8.12140716 | 1.718255034 | 2.077115254 | 0.000142 | 0.001593 | 0.23336 | 0.003365 | 0.120786 |
| 1.96247053 | 13.65804836 | 2.762128945 | 4.064952139 | 0.002124 | 0.007922 | 0.011606 | 0.019869 | 0.182012 |
| 1.026253178 | 1.710496018 | 1.565642907 | 1.329308619 | 0.01045 | 0.042794 | 0.900367 | 0.152616 | 0.050586 |
| 1.131985523 | 1.453568401 | 0.644266873 | 0.819791304 | 0.002124 | 0.00118 | 0.247284 | 0.007252 | 0.036141 |
| 1.690958738 | 1.19831329 | 0.993467733 | 0.884463206 | 0.002557 | 0.032897 | 0.164627 | 0.193267 | 0.97217 |

| |
|---|
| elo3::elo3-gfp |
| 0.008714 |
| 0.75957 |
| 0.005338 |
| 0.218241 |
| 0.097783 |
| 0.823822 |
| 0.03456 |
| 0.269703 |
| 0.75957 |
| 0.019231 |
| 0.005338 |
| 0.021697 |
| 0.085788 |
| 0.031568 |
| 0.031568 |
| 0.024446 |
| 0.474691 |
| 0.002895 |
| 0.45163 |
| 0.45163 |
| 0.305381 |
| 0.618034 |
| 0.005338 |
| 0.148778 |
| 0.021697 |
| 0.822942 |
| 0.034196 |
| 0.022659 |
| 0.605054 |
| 0.026059 |
| 0.082568 |
| 0.076712 |
| 0.374863 |
| 0.021697 |
| 0.172326 |
| 0.344673 |
| 0.005338 |
| 0.005338 |
| 0.009814 |
| 0.240983 |
| 0.906631 |
| 0.277656 |
| 0.305381 |
| 0.828062 |
| 0.097783 |
| 0.038855 |
| 0.601983 |
| 0.021697 |
| 0.828062 |
| 0.75957 |
| 0.136417 |
| 0.004234 |
| 0.944851 |
| 0.021697 |
| 0.009814 |
| 0.067496 |
| 0.828062 |
| 0.665643 |
| 0.834588 |

| |
|---|
| 0.388639 |
| 0.008714 |
| 0.01794 |
| 0.558347 |
| 0.148778 |
| 0.047058 |
| 0.484497 |
| 0.021697 |
| 0.269703 |
| 0.202856 |
| 0.721031 |
| 0.907314 |
| 0.680563 |
| 0.009814 |
| 0.125496 |
| 0.013768 |
| 0.008714 |
| 0.143588 |
| 0.148778 |
| 0.097783 |
| 0.721031 |
