## Supplemental Table 2 for "Fatty acid elongases 1-3 have distinct roles in mitochondrial function, growth and lipid homeostasis in *Trypanosoma cruzi*"

| Metabolite name | formula | PubChem Identifier (KEGG/HMDB) | |
|---|---|---|---|
| 1,3-diphopshateglycerate | C3H8O10P2 | C00236 | N/A |
| 1-Methyladenosine | C11H15N5O4 | C02494 | 7490.498233 |
| 1-Methyl-Histidine | C7H11N3O2 | C01152 | 22095536.35 |
| 2,3-dihydroxybenzoic acid | C7H6O4 | C00196 | 84450.63754 |
| 2,3-Diphosphoglyceric acid | C3H8O10P2 | C01159 | 42572.92276 |
| 2-Aminooctanoic acid | C8H17NO2 | HMDB00991 | 328764.9522 |
| 2-dehydro-D-gluconate | C6H10O7 | C00629 | 183974.8396 |
| 2-deoxyglucose-6-phosphate | C6H13O8P | C06369 | 417490.9324 |
| 2-Hydroxy-2-methylbutanedioic acid | C5H8O5 | C02612 | 1511697.919 |
| 2-hydroxygluterate | C5H8O5 | C02630 | 10028203.19 |
| 2-Isopropylmalic acid | C7H12O5 | C02504 | 323401.4067 |
| 2-ketohaxanoic acid | C6H10O3 | HMDB01864 | 185515.3907 |
| 2-keto-isovalerate | C5H8O3 | C00141 | 5430609.752 |
| 2-oxo-4-methylthiobutanoate | C5H8O3S | C01180 | 28722.28026 |
| 2-oxobutanoate | C4H6O3 | C00109 | 446375.6353 |
| 3-hydroxybutyrate | C4H8O3 | HMDB0000011 | 12055799.37 |
| 3-methylphenylacetic acid | C9H10O2 | HMDB02222 | 389583.2779 |
| 3-phosphoglycerate | C3H7O7P | C00197 | 5320857.73 |
| 3-phosphoserine | C3H8NO6P | C01005 | 11268.9737 |
| 3-S-methylthiopropionate | C4H8O2S | C08276 | 447690.5647 |
| 4-aminobutyrate | C4H9NO2 | C00334 | 372283.0182 |
| 4-phosphopantothenate | C9H18NO8P | C03492 | 293471.081 |
| 4-Pyridoxic acid | C8H9NO4 | C00847 | 150763.7234 |
| 5-methoxytryptophan | C12H14N2O3 | HMDB02339 | 18038.63615 |
| 5-methyl-THF | C20H25N7O6 | C00440 | 42183.01037 |
| 5-phosphoribosyl-1-pyrophosphate | C5H13O14P3 | C00119 | 82061.39382 |
| 6-phospho-D-gluconate | C6H13O10P | C00345 | 79248.3983 |
| 7,8-dihydrofolate | C19H21N7O6 | C00415 | 6116.000308 |
| 7-methylguanosine | C11H16N5O5 | HMDB01107 | 240884.3581 |
| acadesine | C9H14N4O5 | D02742 | 181112.6445 |
| acetoacetate | C4H6O3 | C00164 | 388535.36 |
| acetoacetyl-CoA-posi | C25H40N7O18P3S | C00332 | 521597.1068 |
| Acetylcarnitine DL | C9H18NO4 | C02571 | 69819736.21 |
| acetyl-CoA-posi | C23H38N7O17P3S | C00024 | 875456.4127 |
| Acetyllysine | C8H16N2O3 | C02727 | 97067246.81 |
| acetylphosphate | C2H5O5P | C00227 | 2087886.282 |
| aconitate | C6H6O6 | C00417 | 535829.2538 |
| adenine | C5H5N5 | C00147 | 1685090.631 |
| adenosine | C10H13N5O4 | C00212 | 2159832.667 |
| adenosine 5-phosphosulfate | C10H14N5O10PS | C00224 | 290660.4211 |
| Adenylosuccinate | C14H18N5O11P | HMDB0248010 | 2506.807197 |
| ADP-D-glucose | C16H25N5O15P2 | C00498 | 70289.81491 |
| ADP-nega | C10H15N5O10P2 | C00008 | 25429475.19 |
| a-ketoglutarate | C5H6O5 | C00026 | 183082.6194 |
| alanine | C3H7NO2 | C00041 | 114007408.4 |
| allantoate | C4H8N4O4 | C00499 | 133984.1394 |
| allantoin | C4H6N4O3 | C01551 | 2833710.682 |
| Aminoadipic acid | C6H11NO4 | C00956 | 4019088.585 |
| aminoimidazole carboxamide ribonucleotide | C9H15N4O8P | C04677 | 110758.2568 |
| AMP | C10H14N5O7P | C00020 | 80167449.06 |
| anthranilate | C7H7NO2 | C00108 | 547705.7089 |
| arginine | C6H14N4O2 | C00062 | 161732548.8 |
| L-Arginosuccinic acid | C10H18N4O6 | HMDB0000052 | 136649.689 |
| Ascorbic acid | C6H8O6 | C00072 | 271993.9669 |
| asparagine | C4H8N2O3 | C00152 | 3238248.916 |
| aspartate | C4H7NO4 | C00049 | 5694136.446 |
| ATP-nega | C10H16N5O13P3 | C00002 | 20544468.08 |
| Atrolactic acid | C9H10O3 | HMDB00475 | N/A |
| betaine | C5H11NO2 | C00719 | 138420750.8 |
| betaine aldehyde | C5H12NO | C00576 | 510504.6938 |
| biotin | C10H16N2O3S | C00120 | 679599.9606 |

| | | | |
|---|---|---|---|
| Carbamoyl phosphate | CH4NO5P | C00169 | 14964073.16 |
| carnitine | C7H15NO3 | C00318 | 119869666 |
| CDP-choline | C14H27N4O11P2 | C00307 | 4990380.548 |
| CDP-ethanolamine | C11H20N4O11P2 | C00570 | 274025.6074 |
| CDP-nega | C9H15N3O11P2 | C00112 | 421177.8104 |
| Cellobiose | C12H22O11 | C00185 | 5857.972136 |
| cholesterol | C27H46O | C00187 | 1121038.098 |
| cholesteryl sulfate | C27H46O4S | HMDB00653 | 17668530.36 |
| Cholic acid | C24H40O5 | C00695 | 20392.91098 |
| choline | C5H14NO | C00114 | 573500.5579 |
| Citraconic acid | C5H6O4 | C02226 | 1673644.64 |
| citrate | C6H8O7 | C00158 | 11230460.71 |
| citrate-isocitrate | C6H8O7 | C00158 | 26228398.13 |
| citrulline | C6H13N3O3 | C00327 | 4170203.878 |
| CMP | C9H14N3O8P | C00055 | 5437843.757 |
| coenzyme A-posi | C21H36N7O16P3S | C00010 | 56268.7499 |
| creatine | C4H9N3O2 | C00300 | 63466084.64 |
| Creatinine | C4H7N3O | C00791 | 35472681.74 |
| CTP-nega | C9H16N3O14P3 | C00063 | 150136.7103 |
| cyclic bis(3->5) dimeric GMP | C20H24N10O14P2 | C16463 | 1641.229816 |
| cyclic-AMP | C10H12N5O6P | C00575 | 248668.8457 |
| cystathionine | C7H14N2O4S | C02291 | 14938871.63 |
| cysteine | C6H12N2O4S2 | C00491 | 5745.510672 |
| cysteine sulfinate | C3H7NO4S | C00606 | 3362.714173 |
| Cystine | C6H12N2O4S2 | C00491 | 18395.06752 |
| cytidine | C9H13N3O5 | C00475 | 89818.87439 |
| cytosine | C4H5N3O | C00380 | 115014.4594 |
| dAMP | C10H14N5O6P | C00360 | 1711721.509 |
| dATP-nega | C10H16N5O12P3 | C00131 | 142218.9896 |
| dCDP-nega | C9H15N3O10P2 | C00705 | 191289.5928 |
| dCMP | C9H14N3O7P | C00239 | 600364.1573 |
| dCTP-nega | C9H16N3O13P3 | C00458 | 106222.8976 |
| Dehydroascorbic acid | C6H6O6 | HMDB0001264 | 98599.49534 |
| deoxyadenosine | C10H13N5O3 | C00559 | 38682.2135 |
| Deoxycholic acid | C26H43NO5 | C04483 | 2507.383404 |
| deoxyguanosine | C10H13N5O4 | C00330 | 142137.9542 |
| deoxyinosine | C10H12N4O4 | C05512 | 353520.4958 |
| deoxyribose-phosphate | C5H11O7P | C00673 | 276161.3207 |
| deoxyuridine | C9H12N2O5 | C00526 | 10375.56427 |
| dephospho-CoA-nega | C21H35N7O13P2S | C00882 | 159555.9002 |
| dephospho-CoA-posi | C21H35N7O13P2S | C00882 | 12722.9972 |
| D-erythrose-4-phosphate | C4H9O7P | C00279 | 897284.9984 |
| dGDP-nega | C10H15N5O10P2 | C00361 | 22490613.18 |
| D-glucarate | C6H10O8 | C00818 | 373009.4257 |
| D-gluconate | C6H12O7 | C00257 | 2434332.36 |
| D-glucono-?-lactone-6-phosphate | C6H11O9P | C01236 | 35269.4221 |
| D-glucosamine-1-phosphate | C6H14NO8P | C00352 | 5083.145878 |
| D-glucosamine-6-phosphate | C6H14NO8P | C00352 | 30379.86549 |
| D-glyceraldehdye-3-phosphate | C3H7O6P | C00118 | 3320971.138 |
| dGMP | C10H14N5O7P | C00362 | 36247.54039 |
| dGTP | C10H16N5O13P3 | C00286 | 19063199.76 |
| dihydroorotate | C5H6N2O4 | C00337 | 447885.062 |
| dihydroxy-acetone-phosphate | C3H7O6P | C00111 | 6327426.609 |
| Diiodothyronine | C15H13I2NO4 | HMDB00582 | 1689.604292 |
| dimethylglycine | C4H9NO2 | C01026 | 1582945.257 |
| DL-Pipecolic acid | C6H11NO2 | C00408 | 95683314.28 |
| D-sedoheptulose-1-7-phosphate | C7H15O10P | C05382 | 4047730.976 |
| dTDP-nega | C10H16N2O11P2 | C00363 | 235635.976 |
| dTMP | C10H15N2O8P | C00364 | 135240.4298 |
| dTMP-nega | C10H15N2O8P | C00364 | 1970282.74 |
| dTTP-nega | C10H17N2O14P3 | C00459 | 83556.65001 |
| dUMP-nega | C9H13N2O8P | C00365 | 79766.38464 |

| | | | |
|---|---|---|---|
| dUTP-nega | $C_9H_{15}N_2O_{14}P_3$ | C00460 | 19110.41365 |
| ethanolamine | $C_2H_7NO$ | C00189 | 42051.16202 |
| FAD | $C_{27}H_{33}N_9O_{15}P_2$ | C00016 | 485926.3981 |
| Flavone | $C_{15}H_{10}O_2$ | C15608 | 6927.795207 |
| FMN | $C_{17}H_{21}N_4O_9P$ | C00061 | 88273.55147 |
| folate | $C_{19}H_{19}N_7O_6$ | C00504 | 3212.120299 |
| fructose-1,6-bisphosphate | $C_6H_{14}O_{12}P_2$ | C05378 | 6881612.067 |
| fructose-6-phosphate | $C_6H_{13}O_9P$ | C05345 | 531280.3081 |
| fumarate | $C_4H_4O_4$ | C00122 | 5709756.796 |
| GDP | $C_{10}H_{15}N_5O_{11}P_2$ | C00035 | 1322901.876 |
| Geranyl-PP | $C_{10}H_{20}O_7P_2$ | C00341 | 50468.60851 |
| glucono-$\delta$-lactone | $C_6H_{10}O_6$ | C00198 | 46574.33789 |
| glucosamine | $C_6H_{13}NO_5$ | C00329 | 12983.23712 |
| glucose-1-phosphate | $C_6H_{13}O_9P$ | C00103 | 8316089.332 |
| glucose-6-phosphate | $C_6H_{13}O_9P$ | C00668 | 438252.1341 |
| glutamate | $C_5H_9NO_4$ | C00025 | 63882695.31 |
| glutamine | $C_5H_{10}N_2O_3$ | C00064 | 2462653.506 |
| glutathione | $C_{10}H_{17}N_3O_6S$ | C00051 | 282891.806 |
| glutathione disulfide | $C_{20}H_{32}N_6O_{12}S_2$ | C00127 | 4282955.939 |
| glutathione | $C_{10}H_{17}N_3O_6S$ | C00051 | 235634.6139 |
| glycerate | $C_3H_6O_4$ | C00258 | 80660.53434 |
| glycerol 3-phosphate | $C_3H_9O_6P$ | C00093 | 27946784.24 |
| Glycerophosphocholine | $C_8H_{21}NO_6P$ | C00670 | 109003689 |
| glycine | $C_2H_5NO_2$ | C00037 | 2093.545671 |
| glycolate | $C_2H_4O_3$ | C00160 | 51338.22606 |
| glyoxylate | $C_2H_2O_3$ | C00048 | 105659.8 |
| GMP | $C_{10}H_{14}N_5O_8P$ | C00144 | 13074585.04 |
| GTP | $C_{10}H_{16}N_5O_{13}P_3$ | C00044 | 794603.3295 |
| Guanidoacetic acid | $C_3H_7N_3O_2$ | C00581 | 29123588.57 |
| guanine | $C_5H_5N_5O$ | C00242 | 939512.642 |
| guanosine | $C_{10}H_{13}N_5O_5$ | C00387 | 328084.5399 |
| guanosine 5-diphosphate,3-diphosphate | $C_{10}H_{11}N_5O_{17}P_4$ | C01228 | 178070.1238 |
| hexose-phosphate | $C_6H_{13}O_9P$ | C05345 | 21281860.91 |
| histidine | $C_6H_9N_3O_2$ | C00135 | 5568474.769 |
| histidinol | $C_6H_{11}N_3O$ | C00860 | 5827.563993 |
| HMG-CoA | $C_{27}H_{44}N_7O_{20}P_3S$ | C00356 | 4282.976548 |
| homocysteic acid | $C_4H_9NO_5S$ | C16511 | 162658.1334 |
| homocysteine | $C_4H_9NO_2S$ | C00155 | 48913.29464 |
| homoserine | $C_4H_9NO_3$ | C00263 | 54802.78067 |
| Hydroxyisocaproic acid | $C_6H_{12}O_3$ | HMDB00746 | 80879746.11 |
| Hydroxyphenylacetic acid | $C_8H_8O_3$ | C05852 | 357424.605 |
| hydroxyphenylpyruvate | $C_9H_8O_4$ | C01179 | 10510.30656 |
| hydroxyproline | $C_5H_9NO_3$ | C01157 | 37455034.52 |
| hypoxanthine | $C_5H_4N_4O$ | C00262 | 52236990.7 |
| IDP | $C_{10}H_{14}N_4O_{11}P_2$ | C00104 | 3247854.412 |
| Imidazoleacetic acid | $C_6H_8N_2O_2$ | C02835 | 209711.8948 |
| IMP | $C_{10}H_{13}N_4O_8P$ | C00130 | 10333159.72 |
| indole | $C_8H_7N$ | C00463 | 937845.5738 |
| Indole-3-carboxylic acid | $C_9H_7NO_2$ | HMDB03320 | 2087939.389 |
| Indoleacrylic acid | $C_{11}H_9NO_2$ | HMDB00734 | 784911.4213 |
| inosine | $C_{10}H_{12}N_4O_5$ | C00294 | 7587643.377 |
| isocitrate | $C_6H_8O_7$ | C00311 | 228510.8815 |
| itaconic acid | $C_5H_6O_4$ | HMDB0002092 | 808939.3585 |
| Kynurenic acid | $C_{10}H_7NO_3$ | C01717 | 2755836.128 |
| Kynurenine | $C_{10}H_{12}N_2O_3$ | C00328 | 265687.3322 |
| lactate | $C_3H_6O_3$ | C00186 | 69302660.97 |
| L-arginino-succinate | $C_{10}H_{18}N_4O_6$ | C03406 | 170787.9818 |
| leucine-isoleucine | $C_6H_{13}NO_2$ | C00123 | 227432731.3 |
| lipoate | $C_8H_{14}O_2S_2$ | C00725 | 7546.6044 |
| lysine | $C_6H_{14}N_2O_2$ | C00047 | 41096759.06 |
| malate | $C_4H_6O_5$ | C00149 | 34076504.61 |
| Maleic acid | $C_4H_4O_4$ | C01384 | 5197549.859 |

| | | | |
|---|---|---|---|
| malonyl-CoA | C24H38N7O19P3S | C00083 | 7490.203039 |
| methionine | C5H11NO2S | C00073 | 20731924.99 |
| Methionine sulfoxide | C5H11NO3S | HMDB02005 | 3916614.45 |
| Methylcysteine | C4H9NO2S | HMDB0002108 | 3143633.349 |
| Methylmalonic acid | C4H6O4 | C02170 | 137215253.6 |
| methylnicotinamide | C7H9N2O | C02918 | 4726.301331 |
| myo-inositol | C6H12O6 | C00137 | 160646200.1 |
| N6-Acetyl-L-lysine | C8H16N2O3 | C02727 | 85853584.88 |
| N-acetyl spermidine | C9H21N3O | C00612 | 9185.227292 |
| N-acetyl spermine | C12H28N4O | C02567 | 58236.42563 |
| N-acetyl-aspartylglutamic acid | C11H16N2O8 | C12270 | 32576.27119 |
| N-acetyl-glucosamine | C8H15NO6 | C00140 | 24716.55423 |
| N-acetyl-glucosamine-1-phosphate | C6H14NO8P | C04256 | 151214.1839 |
| N-acetyl-glutamate | C7H11NO5 | C00624 | 611554.5672 |
| N-acetyl-glutamine | C7H12N2O4 | HMDB06029 | 74722.43766 |
| N-Acetyl-L-alanine | C5H9NO3 | C01073 | 44137604.46 |
| N-acetyl-L-aspartic acid | C6H9NO5 | C01042 | 2469.952237 |
| N-acetyl-L-ornithine | C7H14N2O3 | C00437 | 4030.859251 |
| N-Acetylputrescine | C6H14N2O | C02714 | 3178.36613 |
| NAD+ | C21H27N7O14P2 | C00003 | 39470966.5 |
| NADH | C21H29N7O14P2 | C00004 | 40990.22231 |
| NADP+ | C21H28N7O17P3 | C00006 | 4541053.924 |
| NADPH | C21H30N7O17P3 | C00005 | 47671.0383 |
| N-carbamoyl-L-aspartate | C5H8N2O5 | C00438 | 210626.705 |
| Ng,NG-dimethyl-L-arginine | C8H18N4O2 | C03626 | 27682219.69 |
| nicotinamide | C6H6N2O | C00153 | 1600399.598 |
| nicotinamide riboside | C11H15N2O5 | C03150 | 1656.64335 |
| Nicotinamide ribotide | C11H15N2O8P | C00455 | 1940598.857 |
| nicotinate | C6H5NO2 | C00253 | 635575.2495 |
| O8P-O1P (octulose-8-phosphate) | C5H11O8P | 0 | 945443.743 |
| O-acetyl-L-serine | C5H9NO4 | C00979 | 69742.10457 |
| octulose-1,8-bisphosphate (OBP) | C5H11O8P | 0 | 220220.5367 |
| ornithine | C5H12N2O2 | C00077 | 21665559.07 |
| orotate | C5H4N2O4 | C00295 | 323241.7317 |
| orotidine-5-phosphate | C10H13N2O11P | C01103 | 85998.357 |
| oxaloacetate | C4H4O5 | C00036 | 624785.496 |
| p-aminobenzoate | C7H7NO2 | C00568 | 534525.8132 |
| pantothenate | C9H17NO5 | C00864 | 18676015.53 |
| phenylalanine | C9H11NO2 | C00079 | 88982448.7 |
| Phenyllactic acid | C9H10O3 | C01479 | 62906.11705 |
| Phenylpropiolic acid | C9H6O2 | HMDB00563 | 249805.5352 |
| phenylpyruvate | C9H8O3 | C00166 | 30204.722 |
| Phosphocreatine | C4H10N3O5P | C02305 | 31045.55127 |
| phosphoenolpyruvate | C3H5O6P | C00074 | 1880445.423 |
| Phosphorylcholine | C5H15NO4P | C00588 | 56534886.19 |
| p-hydroxybenzoate | C7H6O3 | C00156 | 134501.2642 |
| prephenate | C10H10O6 | C00254 | 3343.177409 |
| proline | C5H9NO2 | C00148 | 113375736.8 |
| propionyl-CoA | C24H40N7O17P3S | C00100 | 5104.635089 |
| purine | C5H4N4 | C00465 | 1399033.628 |
| putrescine | C4H12N2 | C00134 | 30063.19897 |
| Pyridoxamine | C8H12N2O2 | C00534 | 2509.658572 |
| pyridoxine | C8H11NO3 | C00314 | 39503.02259 |
| Pyroglutamic acid | C5H7NO3 | C01879 | 16471266.02 |
| Pyrophosphate | P2H4O7 | C00013 | 75850366.01 |
| pyruvate | C3H4O3 | C00022 | 148191.2332 |
| quinolinate | C7H5NO4 | C03722 | 140899.8474 |
| retinoic acid | C20H28O2 | C00777 | 52666.3979 |
| riboflavin | C17H20N4O6 | C00255 | 1695092.917 |
| ribose-phosphate | C5H11O8P | C00117 | 23404996.29 |
| S-adenosyl-L-homocysteine | C14H20N6O5S | C00021 | 4563106.39 |
| S-adenosyl-L-methioninamine | C14H23N6O3S | C01137 | N/A |

| | | | |
|---|---|---|---|
| S-adenosyl-L-methionine | C15H22N6O5S | C00019 | 12261397.49 |
| sarcosine | C3H7NO2 | C00213 | 89042272.5 |
| sedoheptulose 1,7-bisphosphate (SBP) | C7H16O13P2 | C00447 | 379765.7463 |
| serine | C3H7NO3 | C00065 | 2879292.332 |
| shikimate | C7H10O5 | C00493 | 3587598.851 |
| shikimate-3-phosphate | C7H11O8P | C03175 | 97260.66311 |
| S-methyl-5-thioadenosine | C11H15N5O3S | C00170 | 1744730.793 |
| sn-glycerol-3-phosphate | C3H9O6P | C00093 | 19864587.45 |
| spermidine | C7H19N3 | C00315 | 194092.8396 |
| spermine | C10H26N4 | C00750 | 9246.709884 |
| S-ribosyl-L-homocysteine | C9H17NO6S | C03539 | 162308.8361 |
| succinate | C4H6O4 | C00042 | 137946853.8 |
| succinyl-CoA | C25H40N7O19P3S | C00091 | 85347.04792 |
| taurine | C2H7NO3S | C00245 | 6668792.866 |
| Taurodeoxycholic acid | C26H45NO6S | C05463 | 1008998.499 |
| thiamine | C12H16N4OS | C00378 | 13665.50945 |
| Thiamine pyrophosphate | C12H19N4O7P2S | C00068 | 354790.9089 |
| thiamine-phosphate | C12H17N4O4PS | C01081 | 300399.2922 |
| threonine | C4H9NO3 | C00188 | 3619483.214 |
| thymidine | C10H14N2O5 | C00214 | 59920.12815 |
| thymine | C5H6N2O2 | C00178 | 4703.229967 |
| trans, trans-farnesyl diphosphate | C15H28O7P2 | C00448 | 20894.8116 |
| trehalose-6-Phosphate | C12H23O14P | C00689 | 848332.1772 |
| trehalose-sucrose | C12H22O11 | C00089 | 12118.99669 |
| tryptophan | C11H12N2O2 | C00078 | 22472935.24 |
| tyrosine | C9H11NO3 | C00082 | 8333729.383 |
| UDP-D-glucose | C15H24N2O17P2 | C00029 | 2585634.789 |
| UDP-D-glucuronate | C15H22N2O18P2 | C00167 | 121197.8703 |
| UDP-N-acetyl-glucosamine | C17H27N3O17P2 | C00043 | 599776.0667 |
| UDP | C9H14N2O12P2 | C00015 | 13850972.37 |
| UMP | C9H13N2O9P | C00105 | 3467619.737 |
| uracil | C4H4N2O2 | C00106 | 92719284.03 |
| Urea | CH4N2O | C00086 | 3996583.273 |
| Uric acid | C5H4N4O3 | C00366 | 173117.0324 |
| uridine | C9H12N2O6 | C00299 | 142751.8021 |
| UTP | C9H15N2O15P3 | C00075 | 9336987.04 |
| valine | C5H11NO2 | C00183 | 9861926.246 |
| xanthine | C5H4N4O2 | C00385 | 52824048.58 |
| xanthosine | C10H12N4O6 | C01762 | 1976420.721 |
| xanthosine-5-phosphate | C10H13N4O9P | C00655 | 14219.20622 |
| Xanthurenic acid | C10H7NO4 | C02470 | 1122912.047 |

| WT | | elo1 | | | elo2 | | |
|---|---|---|---|---|---|---|---|
| 101509.4548 | 50168.87044 | N/A | N/A | N/A | N/A | 511593.4194 | N/A |
| N/A | 5019.267383 | 50310.26439 | 41834.95698 | 65286.19969 | 4200.928582 | 12568.3563 | 15929.52805 |
| 21142543.24 | 28208413.33 | 14176246.36 | 11172543.98 | 12123344.12 | 12096067.43 | 12605183.02 | 14523271.9 |
| 129973.5959 | 100551.6034 | 118565.6873 | 160987.2392 | 131648.0302 | 141245.3206 | 141314.5842 | 128651.7589 |
| 101987.468 | N/A | N/A | N/A | N/A | N/A | N/A | N/A |
| 377796.5205 | 312630.8638 | 476276.0776 | 563532.056 | 534559.483 | 526149.1398 | 434578.4355 | 410055.0075 |
| 169870.7567 | 149729.938 | 275071.9213 | 297010.5986 | 290750.3863 | 184510.5191 | 141787.1181 | 180643.7047 |
| 498640.1157 | 496740.1739 | 3282900.572 | 3514561.67 | 3396711.102 | 2440495.95 | 1988988.138 | 2456423.507 |
| 1213550.615 | 1364268.337 | 1519560.446 | 1710053.038 | 1557506.299 | 1900038.386 | 1773802.831 | 1853140.89 |
| 7843284.76 | 8473024.499 | 12482423.8 | 11093388.47 | 10132942.21 | 11306615.8 | 10284586.56 | 11317559.25 |
| 257163.3639 | 460268.2237 | 294095.9109 | 445992.8414 | 324526.2842 | 199731.1532 | 530985.0944 | 173558.6282 |
| 249821.5457 | 202244.1902 | 225010.3558 | 229387.5471 | 196492.2833 | 279104.0839 | 310745.4643 | 242214.0115 |
| 7398885.934 | 5391002.005 | 5971727.85 | 7086168.424 | 6042044.415 | 4629340.789 | 5526471.672 | 5278363.798 |
| 29767.08491 | 18876.87544 | 39303.01934 | 30320.78605 | 76547.30887 | 25932.00842 | 28551.15601 | 30678.50348 |
| 460092.6887 | 392552.0025 | 512961.4935 | 439137.997 | 464996.1934 | 326917.6486 | 353595.2586 | 330347.306 |
| 12889202.93 | 11910428.13 | 34378465.81 | 35284177.84 | 34018630.27 | 19706214.86 | 19428097.14 | 20166944.19 |
| 333273.2372 | 440958.9488 | 868569.5285 | 879594.3948 | 910003.0586 | 580005.1132 | 535119.2345 | 580736.2134 |
| 4358833.672 | 4944601.801 | 38766464.13 | 32641216.79 | 33104036.41 | 33481627.85 | 29663062.3 | 32731697.5 |
| 3350.326976 | 5875.869295 | 4284.057378 | 12511.24698 | 13976.9364 | 1622.366561 | N/A | N/A |
| 535813.4679 | 564602.7485 | 871811.8836 | 779368.0013 | 653556.9122 | 837143.514 | 14027.4155 | 754171.6774 |
| 314527.7556 | 304674.8451 | 391487.4147 | 529943.1576 | 374735.5199 | 229182.3481 | 221600.0985 | 348137.0708 |
| 301016.26 | 278217.9558 | 486233.698 | 415089.2662 | 575956.6302 | 340267.8883 | 306888.8869 | 351294.3915 |
| 147884.8374 | 106614.1878 | 156515.245 | 158887.5409 | 130373.1388 | 106093.8403 | 190864.122 | 140912.8443 |
| 10034.91764 | 21712.01089 | 41043.42837 | 13807.38417 | 24994.47695 | 22436.19954 | 15173.39432 | 22486.71511 |
| 19124.56146 | 32343.08768 | 17434.62033 | 28542.15851 | 37132.39631 | 16439.42766 | 11673.98817 | 24773.7771 |
| 169090.6351 | 86944.66285 | 280265.8101 | 223964.4787 | 198628.0219 | 159483.4469 | 95554.36152 | 217361.7232 |
| 36736.74196 | 82843.59598 | 184296.7352 | 218687.8583 | 176567.4628 | 147207.7924 | 100441.2413 | 109548.2325 |
| 1656.14874 | 9212.981123 | 3345.569488 | 5052.410236 | 4934.952408 | 1673.34073 | 3347.815936 | 2541.771266 |
| 293674.4174 | 248169.5185 | 360219.2285 | 400328.2092 | 523867.0288 | 261702.7501 | 316616.5772 | 339726.7704 |
| 151448.1131 | 142170.5554 | 194015.7036 | 207332.4386 | 200684.714 | 146908.654 | 195771.9282 | 184140.2314 |
| 439021.1401 | 316739.5433 | 360661.4144 | 341499.5742 | 323633.2913 | 292213.4491 | 313910.0042 | 313911.5884 |
| 300744.663 | 617121.7359 | 469560.1162 | 264324.9591 | 471085.6868 | 296774.1766 | 315216.6551 | 300951.461 |
| 57981585.89 | 67485442.09 | 109961143.4 | 112542201.6 | 104136548.2 | 90860732.2 | 94771256.43 | 89771789.14 |
| 347551.3245 | 777493.9794 | 3873495.408 | 3611668.74 | 5767207.484 | 2784633.818 | 3361866.612 | 3100647.784 |
| 71844970.73 | 83717571.23 | 177835083.6 | 198219885.4 | 176435453.6 | 113270626.6 | 112015133.7 | 100219720.7 |
| 1645086.541 | 1524772.543 | 2714370.79 | 2457307.396 | 2887795.604 | 1665222.952 | 1734592.188 | 1526564.17 |
| 658904.2369 | 543134.7213 | 780974.6525 | 810233.3053 | 654553.4072 | 648062.0129 | 712473.1435 | 618479.7381 |
| 1383219.453 | 1649200.709 | 2136640.567 | 8598999.588 | 8600452.472 | 3098459.317 | 3026692 | 4302282.178 |
| 1634989.439 | 1913058.23 | 1477449.118 | 1610145.373 | 2047171.338 | 2516638.426 | 2702350.489 | 2885759.558 |
| 282057.9735 | 259793.6687 | 227964.5198 | 347091.5942 | 262170.421 | 310282.5444 | 390772.6438 | 457128.6733 |
| 7526.709216 | 9174.525453 | 4182.221886 | 12643.0922 | 8479.760322 | 3313.507162 | 7530.565063 | 2508.46596 |
| 57728.64708 | 80306.12354 | 32629.26621 | 48518.78649 | 52734.58571 | 67779.91988 | 52707.74792 | 44339.20568 |
| 25359585.78 | 24030521.52 | 24093432.64 | 24050799.16 | 21423123.76 | 30366740.12 | 31665084.15 | 37941864.17 |
| 94494.15603 | 112914.5211 | 632309.8656 | 597309.7118 | 659550.6218 | 458873.6014 | 618098.3644 | 479048.4065 |
| 103293666.3 | 111248562.2 | 193369935.2 | 182949464.8 | 197799371.6 | 154879062.6 | 148894369.1 | 162029032.5 |
| 122016.4913 | 180307.6039 | 847309.623 | 783602.2683 | 832063.1055 | 2032693.632 | 3978377.537 | 1795302.306 |
| 2145320.78 | 2640501.79 | 3576616.045 | 3568888.774 | 3668044.696 | 2665252.134 | 3502129.766 | 3398034.46 |
| 4132983.795 | 3706756.322 | 20222231.45 | 17631100.05 | 19190100.42 | 12541427.38 | 12849270.98 | 13842275.52 |
| 9066.672576 | 161335.4842 | 137185.9455 | 157533.421 | 146643.6812 | 112891.8746 | 151757.6198 | 130581.1549 |
| 59333772.18 | 74966273.21 | 49685664.05 | 50146863.73 | 52027530.22 | 77953087.9 | 80853518.88 | 72213106.28 |
| 452799.9005 | 533789.0463 | 523022.1722 | 506430.691 | 500067.6709 | 396783.2386 | 427976.0607 | 475279.9257 |
| 156190821.1 | 153537666.2 | 220195835.3 | 233483767.5 | 210921287.7 | 180317890.1 | 207383346.9 | 208109565.1 |
| 90739.5946 | 156877.5621 | 81851.45569 | 112593.345 | 104578.0909 | 196066.4164 | 111444.6906 | 122421.0987 |
| 247419.463 | 267052.3586 | 1233025.716 | 1406714.382 | 1058651.005 | 411473.0222 | 246184.2679 | 270770.0417 |
| 3081354.718 | 3666524.673 | 8985704.821 | 9034751.061 | 9410384.491 | 5415043.104 | 4816010.273 | 5809441.277 |
| 4820700.909 | 4616901.479 | 7312204.245 | 9641600.019 | 8867062.939 | 5523576.724 | 5887369.074 | 7043397.296 |
| 18338851.8 | 24394071.21 | 29506091.32 | 29990753.19 | 32002712.04 | 51480032.05 | 46037734.02 | 48202280.51 |
| 219317.1986 | 27263.89759 | 516891.3388 | 104743.8171 | 362964.2219 | 204211.316 | 94461.6874 | 109165.9781 |
| 159120812.4 | 149645198.1 | 75604919.99 | 81896961.19 | 81858198.41 | 104945308.8 | 105776508.4 | 109012007.7 |
| 602570.6827 | 475035.2042 | 615579.952 | 541304.8966 | 572599.6998 | 493238.9532 | 532280.9697 | 632393.5111 |
| 487021.5399 | 1052493.486 | 722558.019 | 643546.4781 | 808091.1277 | 743389.3204 | 853904.1549 | 709644.4047 |

| | | | | | | | |
|---|---|---|---|---|---|---|---|
| 13185520.13 | 14695301.19 | 40753092.4 | 38309060.55 | 42454180.33 | 24115916.61 | 22064230.82 | 23122579.65 |
| 126939944.9 | 126260850 | 141375120.3 | 130954463.5 | 145764723.4 | 139985600.6 | 134967803.1 | 140995638 |
| 4980904.386 | 5137678.514 | 2290131.449 | 2700839.41 | 2718380.259 | 5973277.428 | 6274145.685 | 6099059.271 |
| 272879.0725 | 307449.7345 | 1569802.933 | 1276915.821 | 1403565.069 | 731027.0057 | 654551.6757 | 624151.9582 |
| 504800.5926 | 451261.3621 | 413787.8637 | 429754.8717 | 444744.1925 | 443256.9831 | 449621.1744 | 317807.1035 |
| 3328.629414 | 3344.906916 | 16767.07717 | 6857.395553 | 1662.304072 | 7478.17398 | 8346.968874 | 4437.447401 |
| 1942335.27 | 560782.0902 | 5133717.648 | 5160701.46 | 5191418.206 | 3818780.45 | 2923661.925 | 3270709.372 |
| 31384002.1 | 7401487.029 | 42425131.3 | 36776695.82 | 50360218.48 | 29307894.05 | 24614188.26 | 27269543.93 |
| 47442.80832 | 36494.2189 | 47797.26078 | 76909.68154 | 30961.64599 | 43330.74237 | 36043.52129 | 79542.88679 |
| 573087.2526 | 526054.3449 | 964985.7197 | 927416.316 | 1165081.979 | 1280179.578 | 1186095.263 | 1016299.957 |
| 1829189.36 | 1704736.084 | 1529856.973 | 1719562.654 | 1377791.851 | 2369532.295 | 2160025.46 | 1808883.582 |
| 17257005.24 | 16083744.08 | 10596974.91 | 11714979.91 | 11962806.22 | 21569775.11 | 21188650.5 | 22455937.49 |
| 38560521.26 | 34690514.92 | 25593697.37 | 28336674.79 | 28057309.97 | 51259992.59 | 44567019.8 | 41312989.13 |
| 4200969.602 | 4012754.286 | 5207652.564 | 5356485.84 | 4949755.764 | 4648812.478 | 4298910.628 | 4867947.347 |
| 3906495.197 | 5297377.837 | 7315680.992 | 9172468.506 | 7509889.677 | 5326730.888 | 6870044.375 | 6246338.221 |
| 11574.21149 | 72436.00261 | 90467.03856 | 64184.2172 | 107098.0408 | 38175.83878 | 85531.68494 | 41388.39225 |
| 64085383.82 | 65197960.14 | 24284862.99 | 29522039.83 | 27971197 | 45068388.76 | 45364476.87 | 45343439.38 |
| 24066234.24 | 33553587.97 | 53846795.16 | 38906099.2 | 56705395.9 | 38128933.58 | 38872521.69 | 42012069.19 |
| 127784.0547 | 159530.4169 | 201322.1358 | 217191.5551 | 156514.3915 | 234802.2062 | 239532.9114 | 222519.5007 |
| 5688.768083 | 10877.8816 | 15054.12931 | 8134.552434 | 18390.2628 | 5934.919292 | 3343.107913 | 3344.144545 |
| 110394.7539 | 171918.8517 | 521026.3025 | 424240.4482 | 455869.3391 | 291744.6233 | 269863.6724 | 322343.9474 |
| 14396755.67 | 18283845.97 | 13482472.05 | 12626746.13 | 11396946.77 | 17632400.92 | 18846847.01 | 20021518.31 |
| 9237.049702 | 10127.18411 | 5001.094089 | 14629.02869 | 29355.36956 | 7595.26126 | 4229.738224 | 16548.07195 |
| 6695.145093 | 9202.971448 | 8365.376857 | 3328.511848 | 2492.543916 | 5852.56539 | 5019.623712 | 2509.861389 |
| 14649.72392 | 37737.99072 | 14484.71288 | 26421.11368 | 686790.0838 | 3400.629154 | 28023.72734 | 20825.02508 |
| 34153.28365 | 106280.9526 | 460988.6747 | 432561.2411 | 320891.304 | 92960.68409 | 137184.7857 | 120283.043 |
| 114154.2487 | 126531.7122 | 189480.1668 | 193159.5104 | 169459.7354 | 98769.91034 | 74478.10161 | 84394.47151 |
| 1174380.497 | 1194261.144 | 1107928.312 | 1116567.229 | 1089186.583 | 1717177.185 | 2258953.345 | 2174850.856 |
| 116941.0155 | 148098.8744 | 113892.9282 | 92875.12572 | 112674.4969 | 553643.2007 | 446519.0042 | 458800.0403 |
| 156413.2439 | 166977.5242 | 194242.7789 | 228518.6183 | 234504.2564 | 192392.069 | 167895.1146 | 221607.8428 |
| 345348.4682 | 535426.0594 | 698362.7515 | 877924.4465 | 804778.557 | 459444.475 | 664427.3606 | 622705.7359 |
| 64586.87922 | 103738.752 | 90350.9078 | 102829.9314 | 81815.05603 | 151695.3548 | 168176.3978 | 148496.2482 |
| 135260.1147 | 95354.63914 | 197938.3728 | 170734.1345 | 159919.2879 | 132357.3939 | 137198.5207 | 129092.2006 |
| 14003.89204 | 56541.70565 | 53616.66816 | 66483.76325 | 39386.51945 | 187915.6095 | 230584.9102 | 218516.1517 |
| 5017.305064 | N/A | 5019.578136 | 2525.592097 | 4181.361201 | 3348.112723 | 5019.633191 | N/A |
| 82688.93369 | 175027.5947 | 148896.3988 | 134643.6439 | 177828.2513 | 191730.6583 | 238117.4701 | 283361.1704 |
| 453225.1643 | 484639.0157 | 1384022.84 | 1112091.708 | 1248884 | 694453.4165 | 903558.7614 | 900958.1913 |
| 306444.568 | 244772.1526 | 488260.1469 | 453768.7093 | 569451.5084 | 445066.4634 | 423091.0866 | 392632.5365 |
| 17075.05334 | 21233.58685 | 11991.15631 | 9218.969264 | 18231.43645 | 23965.53924 | 18032.27143 | 20123.9799 |
| 101200.4067 | 170559.9713 | 54201.20679 | 35908.70126 | 68349.89527 | 43086.60913 | 47602.4922 | 61878.56117 |
| 11898.70323 | 25895.03534 | 3345.030535 | 8329.930156 | 6086.810192 | 6653.584661 | 7497.280594 | 9206.326079 |
| 643030.5371 | 798678.2085 | 2146180.889 | 2445956.921 | 2322771.612 | 2463846.15 | 1932820.537 | 2172707.307 |
| 24829623.46 | 24918866.35 | 24251353.86 | 22005786.1 | 23124927.74 | 31076374.41 | 32146493.5 | 38195284.71 |
| 250400.1169 | 272864.9073 | 419326.9446 | 428195.5935 | 442318.6683 | 330119.9241 | 271873.3638 | 243130.6983 |
| 2320047.03 | 2282478.287 | 5207944.434 | 4433101.358 | 4454047.134 | 2800794.291 | 2508314.551 | 2199069.146 |
| 50968.28621 | 43961.24595 | 139869.4515 | 60135.23524 | 127490.7638 | 59346.51135 | 58315.40321 | 79439.27773 |
| 17261.716 | 5432.783472 | 13286.57869 | 31519.45844 | 15160.2732 | 10843.62622 | 9165.391714 | 13425.83798 |
| 10875.50566 | 20226.57582 | 143385.7586 | 132995.0954 | 106128.8439 | 66376.4634 | 90056.44356 | 79639.60243 |
| 3257603.656 | 3315891.927 | 4521551.591 | 4339121.136 | 3973773.128 | 2767938.506 | 2930349.594 | 2726617.993 |
| 41054.86114 | 42689.555 | 25081.19597 | 22479.33593 | 43500.08086 | 53056.44273 | 69627.74792 | 42457.79878 |
| 18885900.55 | 22190530.97 | 29292492.58 | 31845187.81 | 32387343.03 | 49535887.57 | 48711598.05 | 41927018.9 |
| 291655.1659 | 409798.5583 | 1364973.899 | 594286.7612 | 405720.0051 | 3361902.251 | 3580813.544 | 4103620.374 |
| 5128786.423 | 6515822.569 | 5213481.938 | 4913814.051 | 5115925.671 | 5126170.888 | 4570398.747 | 5003968.989 |
| N/A | N/A | 1673.20534 | 4181.469741 | 3344.175865 | 1674.078118 | N/A | N/A |
| 1215136.989 | 1395178.093 | 1167389.961 | 1226453.5 | 943665.3028 | 809988.0893 | 679057.1736 | 721260.8855 |
| 91646126.11 | 100339077.1 | 104643171.6 | 28306552 | 98631826.86 | 117658886 | 118809438.3 | 133973265.6 |
| 2559542.012 | 3594079.305 | 13413902.28 | 15385042.5 | 17203932.92 | 9735236.359 | 8275792.028 | 10622703.92 |
| 223230.9618 | 261826.9738 | 304807.868 | 285196.0299 | 312754.9542 | 457686.0088 | 418153.5034 | 490474.4627 |
| 90269.14533 | 160148.0564 | 91744.58746 | 150372.8525 | 119922.9203 | 140351.9368 | 174529.1748 | 221755.3615 |
| 2173005.859 | 2120641.493 | 1635141.832 | 1443013.571 | 1280271.539 | 2567903.42 | 3697624.795 | 2945840.828 |
| 74391.75262 | 96720.67511 | 256065.8118 | 231987.6319 | 185171.5999 | 472481.7944 | 287002.8722 | 429769.5735 |
| 93694.64321 | 81058.13959 | 125004.8466 | 113913.4734 | 143384.2158 | 108725.6291 | 90934.72523 | 111315.3529 |

| | | | | | | | |
|---|---|---|---|---|---|---|---|
| 21749.73288 | 11680.27413 | 12385.95351 | 7540.235858 | 14276.28848 | 18382.86998 | 27609.87845 | 25103.29868 |
| 79797.81556 | 73773.76483 | 63781.00201 | 56184.02972 | 69664.99729 | 67268.93435 | 33257.40995 | 49423.53775 |
| 214178.4616 | 540436.8598 | 481002.0388 | 445078.2392 | 653058.032 | 450112.681 | 509911.634 | 473514.1573 |
| 8983.422391 | 4871.48698 | 29071.75613 | 14343.7814 | 14045.00174 | 14420.54516 | 4653.186794 | 11419.09622 |
| 137282.6365 | 39447.08782 | 87191.92639 | 67513.40611 | 78767.45206 | 184137.453 | 303617.8729 | 204252.7704 |
| 5019.975756 | 3363.889624 | 10876.30957 | 5035.231904 | 5055.218887 | 6626.326558 | 10876.22726 | 5205.091238 |
| 8736818.183 | 9099200.516 | 10684488.6 | 14735044.22 | 14819449.74 | 6435498.122 | 5149064.705 | 5567901.723 |
| 421088.0843 | 521169.3301 | 5309939.45 | 6068189.787 | 6036294.446 | 3917768.716 | 4007785.98 | 4224102.03 |
| 7196529.815 | 5265909.037 | 6537243.351 | 7241769.701 | 7162184.846 | 4539945.765 | 5898136.407 | 5211325.343 |
| 1278977.186 | 1280239.677 | 605968.832 | 829276.6017 | 828894.2972 | 955910.6452 | 676854.1044 | 999884.8488 |
| N/A | 70921.57216 | 231039.3182 | 125471.6905 | 209600.7117 | 102990.0148 | 107958.4613 | 133448.4283 |
| 41264.95821 | 48825.76715 | 214333.2448 | 212100.0206 | 221097.7734 | 46158.3025 | 42647.09764 | 49369.54952 |
| 12990.55784 | 38370.99084 | 45965.99233 | 144836.4723 | 134934.4384 | 32577.099 | 26219.23289 | 63529.75086 |
| 5906079.407 | 7141453.667 | 23137956.6 | 26244469.69 | 29619984.87 | 13270603.5 | 12586207.81 | 13201863.22 |
| 503682.6609 | 545824.7974 | 4007983.306 | 4933316.164 | 4967009.024 | 3728339.134 | 3889462.627 | 3298062.615 |
| 50230427.54 | 62280671.75 | 64631465.03 | 64046926.66 | 61020744.98 | 67561000.52 | 70768352.86 | 65787777.96 |
| 2098281.404 | 2630180.055 | 16169748.19 | 14735056.88 | 19580992.85 | 11182887.73 | 12521843.27 | 10555668.6 |
| 116128.4367 | 306545.5576 | 257891.0511 | 175684.8993 | 213318.4169 | 247363.2151 | 252662.3416 | 268167.6741 |
| 3929488.497 | 5195576.719 | 1640486.815 | 1615464.974 | 1588873.38 | 4178738.849 | 4342672.863 | 4438287.331 |
| 226004.3942 | 278152.508 | 151878.4617 | 152001.5222 | 142677.6195 | 149794.3217 | 148951.0103 | 174878.0424 |
| 106225.5516 | 86837.56662 | 240295.2496 | 271143.667 | 222794.0671 | 127758.9581 | 129609.847 | 191793.6912 |
| 21163674.9 | 23659363.91 | 24642219.5 | 22547417.81 | 24327769.83 | 35414621.71 | 31299542.06 | 32922108.89 |
| 99215265.17 | 105768239.4 | 137620824 | 140424899.1 | 133103477.5 | 144880224.8 | 128846422.9 | 145249091.7 |
| 5854.406603 | 3346.73433 | 9201.586984 | 9201.15471 | 3356.218838 | 1674.137656 | 7508.598013 | N/A |
| 45238.15081 | 59242.23002 | 108976.4534 | 101664.01 | 110339.9091 | 28669.74205 | 76035.44166 | 59876.05463 |
| 175204.26 | 119501.0855 | 135555.1954 | 120954.5363 | 128689.3565 | 95040.59648 | 159532.8727 | 126305.0952 |
| 8062652.208 | 15272581.36 | 5734293.687 | 6487614.081 | 8631810.465 | 12359346.2 | 17031135.15 | 10370185.89 |
| 1215110.965 | 1240233.721 | 535277.0321 | 792640.8676 | 498587.9925 | 1780988.228 | 1928378.897 | 1928859.934 |
| 26729140.43 | 28838761.78 | 31338113.69 | 30844679.66 | 32791353.14 | 30786576.87 | 30672342.72 | 33749403.02 |
| 614527.035 | 1017170.076 | 1948183.212 | 2091884.749 | 2295411.253 | 1289586.382 | 1139167.683 | 1402526.532 |
| 234025.5527 | 254717.6087 | 182798.2729 | 222550.0241 | 217563.3978 | 478554.3498 | 500968.1908 | 629087.4374 |
| 354348.5589 | 116600.5233 | 75440.00206 | 107559.0599 | 163233.375 | 103509.4789 | 47888.89961 | 44322.01642 |
| 18063260.87 | 23553681.62 | 54377051 | 56014159.57 | 57102980.91 | 55475800.06 | 53917011.66 | 52590686.49 |
| 5559915.761 | 5992073.431 | 7524044.54 | 8243040.199 | 7752340.742 | 8837486.909 | 9293645.731 | 9583206.553 |
| 4180.952035 | 11712.99679 | 13391.37642 | 11714.76237 | 13385.12028 | 10876.53372 | 3345.706173 | 5864.871382 |
| 3343.940052 | 15049.27193 | 25968.37735 | 21566.98152 | 26633.87191 | 15047.67925 | 20687.03129 | 11639.2449 |
| 125853.9587 | 138818.9599 | 162906.7314 | 129443.0595 | 148035.2676 | 112610.1712 | 69211.38391 | 88943.42935 |
| 36801.69097 | 70166.50084 | 112341.0909 | 121713.05 | 72775.80301 | 73896.94168 | 55209.08335 | 73588.21634 |
| 73466.57519 | 81943.47867 | 115684.094 | 105357.9307 | 88516.25362 | 75311.69675 | 122790.6241 | 93016.79375 |
| 212597008.1 | 243299478.1 | N/A | 81903261.55 | N/A | 244970627.1 | 211877402.5 | 80468839.83 |
| 287244.5252 | 311741.598 | 424016.5999 | 385802.1151 | 380376.9477 | 225327.9356 | 284149.2265 | 274001.6764 |
| N/A | 12682.63591 | 15208.21742 | 2030.365282 | 2800.279953 | 5526.6619 | 33289.7374 | 13763.21662 |
| 35159885.73 | 41554714.09 | 21278008.21 | 19980741.32 | 23812654.1 | 31808776.5 | 41303589.81 | 32821768.79 |
| 49577106.7 | 45409128.5 | 75295073.55 | 67263746.52 | 73730136.41 | 45234961.24 | 50407570.13 | 48654245.25 |
| 3312229.816 | 3570419.572 | 3146050.643 | 3297587.598 | 2795140.621 | 4850768.842 | 4855957.856 | 5350729.587 |
| 116841.561 | 100744.9536 | 220382.3084 | 203548.7193 | 236645.2711 | 203549.3773 | 171385.7059 | 152954.1616 |
| 6335098.533 | 10111046.91 | 3880836.913 | 3854978.507 | 3277239.436 | 8171795.676 | 10093711.71 | 9170240.322 |
| 530422.4553 | 947039.7258 | 1102342.336 | 955398.551 | 1124885.347 | 882640.8189 | 929641.8681 | 1004383.11 |
| 2229400.432 | 1969809.721 | 10633354.98 | 10254306.9 | 10099788.49 | 7057392.734 | 7260700.22 | 7397922.54 |
| 647867.0523 | 822534.2953 | 584247.1092 | 565099.1529 | 697016.7464 | 623892.0868 | 584490.689 | 656379.6022 |
| 5901035.169 | 8346562.249 | 8593502.572 | 7864066.606 | 9183353.43 | 13021285.61 | 12331087.63 | 12327829.42 |
| 245267.4296 | 128441.4591 | 335506.9523 | 309552.1063 | 287818.555 | 171171.6423 | 128568.9216 | 169354.1209 |
| 699855.6771 | 727611.9435 | 900544.0539 | 821522.3134 | 1008743.286 | 986648.603 | 818195.9852 | 877892.7424 |
| 2069980.237 | 2239718.79 | 2483183.901 | 2180839.579 | 2372032.951 | 1929343.997 | 1958802.114 | 2021024.801 |
| 145436.8077 | 236900.8967 | 322269.5303 | 368513.2466 | 438228.7397 | 277663.8735 | 290855.4161 | 274506.6385 |
| 69417553.99 | 69402084.16 | 77305245.24 | 76752977.45 | 75371956.89 | 67379851.27 | 71214861.88 | 69345155.48 |
| 76291.45402 | 121437.9986 | 99778.67114 | 104325.4315 | 108923.2994 | 127856.2584 | 143630.2629 | 100489.3183 |
| 187152588 | 234449294.8 | 251467218.1 | 255391249.3 | 245486224.4 | 235153963.7 | 234208654.7 | 244094426.1 |
| 3397.909836 | 3344.978477 | 6693.13951 | 2578.146089 | 6694.362944 | 2411.342151 | N/A | 1673.800433 |
| 33645503.99 | 38426608.76 | 44293663.74 | 41194237.77 | 40240038.94 | 31466102.48 | 34084707.43 | 36967225.75 |
| 42998605.52 | 33052762.46 | 31705649.71 | 28520290.57 | 28449073.74 | 29489624.8 | 36531578.2 | 31717703.69 |
| 7688398.074 | 5156157 | 6763140.981 | 6711991.87 | 6357014.737 | 4927119.916 | 5523887.239 | 5305442.364 |

| | | | | | | | |
|---|---|---|---|---|---|---|---|
| 1576.867114 | 21749.55624 | 728654.0865 | 536676.2321 | 624294.7964 | 102002.3158 | 85269.66713 | 66937.90561 |
| 15252932.87 | 23738846.02 | 34656769.29 | 35000372.5 | 37489173.66 | 26538908.35 | 28494635.48 | 27210294.62 |
| 3682380.285 | 4121555.464 | 7799651.914 | 7585303.696 | 7244744.215 | 5883646.232 | 4555174.512 | 5421529.151 |
| 2697175.792 | 2642543.975 | 5922839.001 | 7117225.405 | 6157440.85 | 4133105.455 | 4192068.776 | 3839437.186 |
| 132633523.3 | 133540370.1 | 111056436.9 | 111852394.2 | 113889888.9 | 130333380.6 | 130648934.4 | 132134140.5 |
| 4165.408214 | 26779.14654 | 4336.978567 | 7879.610652 | 8504.324235 | 8322.027579 | 17363.1567 | 3019.873965 |
| 156953168.8 | 151657556.1 | 108406820.6 | 104312282.2 | 106691424.3 | 136447882.5 | 141781639.7 | 148249709 |
| 98919011.07 | 99239091.28 | 210689846.7 | 199514923.7 | 173963428.4 | 135830144 | 146282313.1 | 119668469.5 |
| 3328.640506 | 17556.64475 | 14221.12563 | 22340.51148 | 11452.62062 | 15016.80487 | 7433.062827 | 6944.899323 |
| 39051.43014 | 132907.9609 | 100732.5913 | 94588.24347 | 98523.99487 | 79557.97534 | 71304.99429 | 76856.36963 |
| 14099.1757 | 28385.55868 | 33467.01615 | 20083.91935 | 22558.21401 | 21752.29309 | 32646.79117 | 19241.09564 |
| 15623.3338 | 30088.11705 | 108778.9242 | 110330.4088 | 156923.2156 | 109133.6464 | 97923.8627 | 128110.9589 |
| 145499.6217 | 88621.59721 | 118489.9513 | 280096.9213 | 207897.5645 | 247686.3342 | 207090.4279 | 223225.0423 |
| 344490.2269 | 496243.9556 | 290493.4484 | 391200.4815 | 428327.4013 | 364886.0311 | 435715.2383 | 394291.83 |
| 60715.44952 | 59774.00056 | 40665.46266 | 33769.8605 | 18948.41871 | 51246.59814 | 50099.55893 | 52151.27295 |
| 53118970.61 | 45916760.49 | 19683891.9 | 21300039.27 | 19950364.42 | 33675953.29 | 34350499.39 | 33666933.07 |
| 4216.758729 | 6624.97617 | 16732.46002 | 14226.12554 | 17603.51746 | 10929.26151 | 13268.67602 | 8344.338041 |
| 2917.426849 | 4341.570109 | 3260.114355 | 2229.039117 | 7241.295982 | 8633.697504 | 2296.798709 | 3878.702352 |
| 8258.742649 | 11747.73348 | 7455.940781 | 8711.476018 | 6705.622635 | 14430.21746 | 6758.539145 | 2473.079276 |
| 34353583.35 | 39374585.49 | 11466018.97 | 10282149.93 | 9795657.267 | 45151498.79 | 40237112.37 | 44509458.39 |
| 56049.69048 | 57726.03714 | 15885.92713 | 6653.540341 | 11713.53711 | 47683.74856 | 59396.62806 | 62752.86282 |
| 3087459.195 | 3909096.369 | 3664006.986 | 3095264.967 | 3185142.591 | 5400760.386 | 5701720.149 | 6148000.966 |
| 29252.13922 | 28606.04125 | 32627.76238 | 20075.49612 | 31385.33375 | 35383.62205 | 47686.99942 | 56891.316 |
| 155550.8178 | 207015.096 | 2012729.135 | 2021435.717 | 2468665.804 | 526948.7286 | 643386.9317 | 549270.774 |
| 31958215.18 | 36292768.84 | 50011849.29 | 39962918.59 | 42722988.41 | 35730065.99 | 33367711.74 | 34045275.56 |
| 608714.6772 | 1827604.463 | 810376.2132 | 623707.252 | 439682.4548 | 5562610.172 | 6713454.921 | 6535267.484 |
| 10531.46637 | 39566.19141 | 4853.23838 | 6432.202279 | 33313.5804 | N/A | 9502.370132 | 14167.83059 |
| 2040234.968 | 2417949.728 | 229659.8187 | 185286.2945 | 247566.9207 | 951909.3428 | 1123693.239 | 891626.8432 |
| 531088.7801 | 628979.1675 | 664397.4185 | 632860.2606 | 638138.5508 | 525243.2675 | 579826.091 | 740452.6501 |
| 655469.8824 | 880493.1219 | 2023940.309 | 2932649.218 | 2557664.31 | 3864915.428 | 2631162.065 | 3604908.977 |
| 53489.39098 | 55555.72393 | 181674.2003 | 108665.897 | 166339.7846 | 207007.8754 | 97128.15917 | 95752.44665 |
| 247857.7954 | 249477.1484 | 1414222.799 | 1494925.964 | 1610956.095 | 388555.7088 | 328376.7605 | 380834.5446 |
| 28698583.41 | 27560696.61 | 28681056.23 | 38128094.91 | 33250901.44 | 16205967.55 | 25204144.98 | 22188684.85 |
| 244111.1611 | 290242.7135 | 161893.9428 | 151796.6368 | 232875.2544 | 254345.268 | 221175.9471 | 275053.6163 |
| 56118.83873 | 155806.4264 | 179094.201 | 156399.8553 | 158199.2046 | 105624.9804 | 127363.6936 | 141600.8667 |
| 580700.9248 | 492004.9883 | 687584.9969 | 1892554.697 | 2001663.154 | 618063.3065 | 721534.8672 | 609492.8837 |
| 393154.0801 | 500710.3748 | 527445.1423 | 497923.6896 | 544958.645 | 336478.3808 | 456739.7547 | 483756.3483 |
| 19973456.54 | 18551869.96 | 29327970.32 | 28702585.21 | 26306175.78 | 18365988.92 | 18207934.35 | 17488677.69 |
| 62577522.08 | 85658640.3 | 97035121.69 | 92670015.1 | 102643387.9 | 77446316.87 | 79456604.76 | 83268000.71 |
| 341771.4684 | 207008.0523 | 606898.9218 | 78381.53443 | 442749.0998 | 215689.5809 | 177258.9592 | 66266.44752 |
| 258611.6768 | 182364.7621 | 1249342.378 | 1161388.965 | 1295019.572 | 950284.2988 | 1107733.851 | 827152.7334 |
| 3103.733673 | 5061.692191 | 89754.91286 | 83144.1183 | 61795.5535 | 30608.26048 | 20516.12239 | 12180.0708 |
| 77709.60441 | 52600.23513 | 78953.16137 | 58410.76464 | 128712.964 | 99122.14074 | 105661.112 | 140026.5277 |
| 1709078.747 | 2205956.474 | 15834724.49 | 10891853.85 | 14007691.94 | 17425147.39 | 13959478.92 | 16243467.76 |
| 63770402.93 | 56766684.73 | 169687498.7 | 142825114.8 | 165152944.8 | 248743252.1 | 249530755.2 | 233983292.1 |
| 127951.4209 | 51696.1014 | 126496.0491 | 246507.7078 | 176992.7905 | 143246.1743 | 142187.892 | 118667.0296 |
| N/A | 1640.59535 | N/A | 1674.134664 | N/A | 1673.94281 | 1658.367082 | 2911.837564 |
| 140640211.5 | 136455949.2 | 93057090.47 | 91851872.1 | 89056868.59 | 129508227.9 | 198757119.1 | 145127543.9 |
| 7548.144965 | 4414.545401 | 15058.57552 | 16600.55146 | 16770.63483 | 7395.713294 | 10039.6656 | 6692.50968 |
| 709076.0253 | 1350701.626 | 1622056.123 | 1553204.31 | 1679751.641 | 1135308.476 | 1145497.247 | 1262052.735 |
| 14067.40195 | 57814.35003 | N/A | 47701.14293 | 89559.21749 | 121671.7901 | 65851.02341 | N/A |
| 2509.58198 | 4708.99305 | 2483.922248 | 4599.903346 | 1649.337686 | 1633.906466 | 5920.486591 | 4822.928069 |
| 68949.68508 | 134153.8065 | 901031.4196 | 807479.0573 | 1037690.972 | 1122268.02 | 1240418.026 | 1348231.842 |
| 15728287.1 | 15590580.92 | 20266028.89 | 20856676.82 | 20042589.36 | 14723970.03 | 15990794.67 | 16642680.37 |
| 70489127.63 | 74866800.54 | 108731506.3 | 118138735.5 | 38773769.14 | 77229337.11 | 80242076.82 | 75206914.51 |
| 168539.2787 | 155024.0314 | 211217.0423 | 188975.2229 | 179873.4954 | 181049.376 | 168851.3233 | 165290.3753 |
| 151021.9477 | 152965.3125 | 156515.3375 | 127632.2313 | 130231.3574 | 133317.8088 | 127682.7466 | 107674.1137 |
| 34284.22818 | 2973.753785 | 25246.39807 | 43306.22184 | 26995.51647 | 43846.77209 | 35926.45085 | 35008.29316 |
| 1280880.643 | 1342036.421 | 871138.3503 | 718059.8807 | 985113.6753 | 2770938.951 | 2999016.527 | 3384236.584 |
| 26328583.03 | 23968966.69 | 21510777.64 | 27997234.11 | 25963784.83 | 19775302.44 | 25491187.27 | 23315565.68 |
| 4050927.794 | 3942897.014 | 1128387.884 | 1158790.942 | 1160841.89 | 5030839.042 | 5102424.752 | 5658839.019 |
| N/A | N/A | 1641.230975 | N/A | N/A | 5001.973871 | 4172.252089 | N/A |

| | | | | | | | |
|---|---|---|---|---|---|---|---|
| 14200949.3 | 12499298.86 | 19593015.67 | 14149461.2 | 19504300.25 | 11723734.05 | 13475355.83 | 19598238.86 |
| 80259071.14 | 95000157.4 | 146819024.5 | 144689265.8 | 155010467.4 | 130910698.1 | 135550943.8 | 134765930.3 |
| 373923.9274 | 441533.9479 | 915580.4807 | 813274.782 | 740387.024 | 497241.3095 | 358481.8275 | 368012.5159 |
| 2241003.933 | 2293862.421 | 4644796.596 | 5166536.115 | 3674050.114 | 1923612.931 | 2325299.068 | 2463583.096 |
| 2418304.74 | 2683758.233 | 5402827.671 | 3224079.65 | 4842449.16 | 1535391.726 | 3561597.372 | 3673173.538 |
| 56828.89473 | 93307.93224 | 42051.87191 | 50808.26255 | 44695.70854 | 66772.45289 | 95274.18144 | 59835.41259 |
| 1017483.831 | 1033168.136 | 931724.91 | 774855.6699 | 804482.4876 | 945710.6531 | 966552.9516 | 802616.1332 |
| 18909035.97 | 22942485.36 | 13849952.92 | 14592125.2 | 12478915.16 | 24983463.59 | 30104759.39 | 27549152.49 |
| 89518.50853 | 197414.0903 | 58906.16753 | 57552.55709 | 136366.3032 | 132047.8887 | 138038.4908 | 177968.5015 |
| 20494.28663 | 5878.594882 | 47286.86836 | 45511.79799 | 40581.66067 | 30639.28456 | 5380.133418 | 29546.87239 |
| 175808.4306 | 138431.3929 | 67059.60385 | 52901.21253 | 96625.37992 | 102314.5508 | 108146.8386 | 149040.2831 |
| 132590943.6 | 135786737.1 | 113682706.6 | 101439745.9 | 113293922 | 128963969.4 | 126949125.4 | 133320022.6 |
| 36794.1755 | 91920.22684 | 38062.55674 | 20523.0643 | 23344.15285 | 20939.90186 | 21203.98485 | 28480.43768 |
| 6771421.208 | 5761141.531 | 10321166.19 | 11420671.6 | 11535489.27 | 7756483.572 | 7264594.34 | 7702878.696 |
| 1538322.189 | 567609.4313 | 2188454.904 | 1953446.658 | 1906774.536 | 814376.2827 | 840477.0513 | 774638.9175 |
| 10958.22862 | 61831.47896 | 15379.7734 | 495161.8551 | 597727.9932 | 561361.9993 | 121173.8933 | 140565.6568 |
| 383917.8601 | 427570.0691 | 2236663.019 | 2103484.491 | 2210064.207 | 1173309.544 | 1393944.377 | 1367570.195 |
| 378765.5392 | 373623.2643 | 1988495.272 | 1995184.507 | 2021206.309 | 783362.4021 | 822481.4235 | 482268.1248 |
| 3544726.465 | 4026656.445 | 28976209.65 | 29535122.47 | 25105573.28 | 15860694.62 | 15664233.5 | 17125298.3 |
| 31723.51009 | 63933.79287 | 68316.02871 | 33291.35954 | 81029.99086 | 40603.26401 | 60771.71763 | 68539.29059 |
| 26669.39467 | 71833.14498 | 27111.90588 | 27368.67244 | 41319.46016 | 25564.24539 | 38347.00421 | 25705.75702 |
| 72124.32217 | 68083.59405 | 140647.0822 | 41255.80845 | 66575.52173 | 34170.92986 | 107491.1144 | 53611.0262 |
| 557278.631 | 932803.4985 | 1246899.672 | 1406591.34 | 1267714.01 | 1046240.201 | 667156.0374 | 777825.346 |
| 18088.02642 | 32214.46619 | 30093.24932 | 44225.91734 | 43549.45076 | 44387.33293 | 40105.1085 | 58013.31073 |
| 15308051.08 | 25642349.97 | 24258253.83 | 21895654.12 | 23968136.84 | 21439665.94 | 22056881.24 | 22752458.43 |
| 8106098.721 | 10029717.18 | 17637361.33 | 17305887.33 | 21110090.39 | 10968356.11 | 14722590.99 | 12741268.77 |
| 2910629.025 | 2752971.829 | 54350451.48 | 49897317.74 | 53456890.71 | 28394917.84 | 32595993.17 | 33707907.7 |
| 133082.4629 | 174874.7136 | 86983.08318 | 121313.2843 | 132157.7877 | 77806.41313 | 204972.7533 | 155611.9865 |
| 614143.6907 | 548803.3107 | 15331060.7 | 11648168.66 | 12315135.22 | 8392905.518 | 8093686.993 | 10540834.01 |
| 16617622.92 | 15306893.61 | 20729562.52 | 17806635.12 | 17337257.88 | 13578662.42 | 17004593.33 | 15226349.34 |
| 2888809.75 | 4074990.116 | 2130201.871 | 2161382.264 | 1787389.758 | 2390638.756 | 2910246.168 | 2849550.736 |
| 78789837.21 | 88412593.78 | 166312498.4 | 174528174 | 172244937.7 | 247721287 | 251896259 | 254952569.8 |
| 2401831.447 | 3012293.966 | 2930551.945 | 2532779.826 | 2977148.867 | 3060364.898 | 3038212.072 | 3008296.692 |
| 148123.6584 | 138155.559 | 409393.5281 | 417490.7547 | 459979.2773 | 282217.1168 | 279098.2802 | 263647.8031 |
| 102607.9157 | 187999.2849 | 244643.1203 | 230576.7022 | 281728.2114 | 265245.6409 | 284609.4353 | 233104.5729 |
| 8985527.998 | 11051940.09 | 14438767.78 | 13989805.15 | 15460547.97 | 14327040.23 | 11884213.38 | 14919078.04 |
| 8071421.312 | 11000616.91 | 14609309.03 | 17114499.14 | 17297431.08 | 15955328.92 | 16816574.42 | 15120165.25 |
| 49369120.02 | 52988521.44 | 78182662.17 | 73548065.45 | 74946251.47 | 58553024.52 | 58324666.84 | 61325990.08 |
| 1497377.727 | 2045123.155 | 2014082.38 | 2260793 | 2371897.421 | 1768219.335 | 1728313.549 | 1984297.826 |
| 15060.18358 | 6693.67806 | 4195.645352 | N/A | 12549.33009 | 5083.825503 | 3326.674494 | 5855.337886 |
| 1170758.645 | 1167851.397 | 1789955.083 | 1544415.698 | 1687677.58 | 1442678.518 | 1277829.711 | 1319697.137 |

| elo3 | | | elo1::elo1-gfp | | | elo2::elo2-gfp | |
|---|---|---|---|---|---|---|---|
| N/A | 82077.07043 | N/A | 106381.0591 | N/A | N/A | N/A | N/A |
| 5863.640772 | 3320.966488 | 9560.145009 | 1658.453589 | 25114.48837 | 1674.077232 | 3350.915911 | 3427.052236 |
| 9811300.44 | 10036374.99 | 8627143.098 | 8095302.15 | 6951525.352 | 6950816.541 | 7515288.184 | 7174313.684 |
| 106209.2947 | 130901.0169 | 114383.8192 | 70636.5949 | 63488.78208 | 106467.3199 | 99978.57023 | 74650.04706 |
| N/A | 15172.74897 | N/A | N/A | 96556.21579 | N/A | 15462.39194 | N/A |
| 440821.8634 | 395763.4458 | 488647.2649 | 478702.1718 | 400854.4333 | 308429.3397 | 418932.484 | 446090.1015 |
| 253432.7689 | 203530.8323 | 268818.9566 | 187317.1886 | 159783.6588 | 163928.576 | 185943.223 | 136390.68 |
| 1098432.411 | 1074011.721 | 993741.6621 | 936216.5294 | 729222.2762 | 701664.1395 | 1056943.689 | 1163411.312 |
| 1376811.105 | 1403342.355 | 1559094.747 | 1230844.364 | 1261682.733 | 1279903.515 | 1216320.816 | 1130759.937 |
| 8391647.749 | 8726811.847 | 8323871.791 | 8181650.857 | 8169757.609 | 8245132.491 | 8553879.523 | 7450133.205 |
| 263137.6209 | 307198.9032 | 342985.8484 | 356085.7755 | 353719.3689 | 60844.16721 | 330609.2162 | 368813.8094 |
| 224057.8821 | 218256.0493 | 160515.875 | 224463.1318 | 191733.0776 | 184558.7531 | 202905.4532 | 187147.811 |
| 4567319.848 | 5109855.668 | 4719652.691 | 4739130.134 | 4051112.108 | 5048818.539 | 4555168.521 | 3848492.158 |
| 29426.99035 | 21735.34684 | 7275.577285 | 17076.19317 | 22028.08168 | 38601.01524 | 18074.30295 | 16518.73485 |
| 341275.904 | 303198.3471 | 423500.4453 | 261221.7747 | 376199.1548 | 233453.2715 | 458521.9423 | 332406.2236 |
| 13273581.22 | 13960572.8 | 12581880.23 | 17908502.01 | 16183861.86 | 16896154.23 | 14325816.4 | 14977895.99 |
| 635317.3049 | 376487.3273 | 392397.2111 | 516342.1202 | 521230.2654 | 444600.017 | 332493.9058 | 286116.9799 |
| 6205717.035 | 4882902.973 | 5700346.401 | 16908089.92 | 16751847.33 | 13696238 | 6641886.863 | 6311874.153 |
| 6621.409011 | 10226.8996 | 6899.39644 | 6530.374995 | 5210.904182 | 7954.812044 | 6114.681911 | 18259.70278 |
| 324285.3633 | 325675.8331 | 311976.7954 | 206743.177 | 200539.1214 | 201615.6592 | 437401.531 | 457520.8018 |
| 280254.3849 | 332023.0724 | 377316.8201 | 294084.8588 | 333824.0654 | 254978.323 | 315990.9315 | 285674.1259 |
| 266049.538 | 255302.4481 | 186083.2779 | 119306.6957 | 152119.3739 | 185532.9628 | 204850.4916 | 296840.6876 |
| 82896.85953 | 96769.17815 | 122476.0108 | 104708.0546 | 94879.43481 | 116609.7755 | 154747.0336 | 112195.9512 |
| 14220.44926 | 25535.16928 | 8144.07793 | 21753.18827 | 29279.72535 | 26190.87221 | 8365.108301 | 10064.12242 |
| 45247.29582 | 41639.44728 | 22938.95283 | 46194.4356 | 31102.13046 | 9118.831572 | 34069.49589 | 27622.0289 |
| 71473.35143 | 3930.374549 | 81883.85623 | 97762.1126 | 74281.82202 | 4945.438126 | 11617.04769 | 62839.27057 |
| 112710.9764 | 87071.92757 | 90364.45765 | 72679.2549 | 57012.46587 | 79996.31618 | 59701.99256 | 65547.53661 |
| 4791.081452 | 2524.787437 | 7481.503599 | 4149.536312 | 5002.629127 | 6712.546166 | 6693.733126 | 7591.402406 |
| 286986.7696 | 306103.112 | 492762.1698 | 237718.0007 | 218031.3344 | 282703.8659 | 401675.1504 | 158325.6886 |
| 177368.7926 | 150524.0881 | 184863.3623 | 188862.0497 | 170707.6519 | 124609.9736 | 129215.7896 | 146409.2376 |
| 410018.5032 | 309140.065 | 422544.6731 | 443044.5725 | 283798.5556 | 310666.9084 | 284136.5927 | 335102.2722 |
| 310746.2249 | 302809.3505 | 373524.7221 | 281879.4237 | 174811.8324 | 154765.9714 | 311188.0537 | 304777.5073 |
| 62638901.94 | 77793363.78 | 80054239.37 | 125278850.3 | 116197824.6 | 125396358.3 | 78306420.84 | 75906365.37 |
| 535502.6601 | 541587.4958 | 441052.4922 | 566801.5238 | 773167.2414 | 421253.2308 | 251197.6672 | 448450.288 |
| 82255001.56 | 91238241.63 | 97597246.27 | 98071874.14 | 111930790.3 | 97310329.49 | 107151934.3 | 87642077.07 |
| 1689542.45 | 615521.2037 | 1339361.781 | 1648226.725 | 1392682.256 | 1251343.15 | 1436691.93 | 1193351.018 |
| 903416.9655 | 795808.3464 | 973773.0971 | 700359.0292 | 679617.0126 | 674989.2218 | 501704.3614 | 589138.8948 |
| 5158035.154 | 1846185.601 | 4744254.229 | 3507452.262 | 3433347.126 | 2746902.019 | 1583700.281 | 1389045.086 |
| 2223517.361 | 2446316.194 | 2842741.251 | 1876111.441 | 2778970.515 | 1833804.145 | 1914836.356 | 1572976.312 |
| 206837.4338 | 246246.0054 | 305169.0788 | 405117.0528 | 418271.7071 | 323859.4021 | 251036.2698 | 351957.8953 |
| 7520.5232 | 7631.131145 | 5133.193008 | 9097.144447 | 5018.068154 | 10841.3657 | 9919.736535 | 2526.88596 |
| 71109.48099 | 91188.61768 | 82808.97417 | 51030.29947 | 98716.63077 | 117965.6059 | 55215.66572 | 73619.193 |
| 26323723.79 | 22750161.39 | 23390181 | 34408464.1 | 37611727.03 | 30235914.48 | 25579078.58 | 32890168.34 |
| 209057.2724 | 307704.8827 | 220544.7712 | 159573.2255 | 189416.7499 | 149809.9995 | 292757.0876 | 245569.6875 |
| 119388702.3 | 109384908.9 | 106042360.2 | 151882434 | 146413499.7 | 143246926.3 | 125004392.2 | 128529858.2 |
| 304149.359 | 223217.3082 | 233535.0758 | 1160914.618 | 1023520.014 | 1159612.162 | 214553.6969 | 192719.485 |
| 3042231.962 | 3011690.861 | 2723269.917 | 2783174.254 | 2734669.434 | 2549479.736 | 2628078.62 | 2605628.746 |
| 3088886.297 | 2999010.965 | 3093165.466 | 3834479.188 | 3454823.345 | 3131724.966 | 6334693.179 | 5995325.358 |
| 138194.8987 | 145476.7741 | 141391.4888 | 142794.4151 | 139120.1145 | 121327.2089 | 128975.4589 | 106473.5264 |
| 89547886.24 | 79825199.8 | 77810409.73 | 89415304.92 | 81395279.63 | 77172341.75 | 88934276.83 | 79587482.06 |
| 499122.0388 | 585230.3873 | 469207.476 | 527085.9941 | 383590.7732 | 388329.4198 | 411703.4954 | 442258.7007 |
| 182326420.7 | 163022133.4 | 193695759.2 | 182592225.3 | 178371764.4 | 196340690 | 92271514.72 | 197570946.4 |
| 121980.1109 | 142097.542 | 108308.8999 | 150421.7017 | 119563.6429 | 80116.12417 | 108646.002 | 62937.24088 |
| 271309.9504 | 325088.0799 | 377820.1939 | 637117.3359 | 503018.1646 | 585384.9897 | 329836.7353 | 304654.1825 |
| 3887445.022 | 3207702.442 | 3280376.305 | 2580867.356 | 2714633.367 | 2321715.798 | 2866421.698 | 2761132.668 |
| 7059484.688 | 7046985.155 | 6163986.981 | 6972610.9 | 7169102.475 | 5411842.509 | 7217796.427 | 4929360.109 |
| 24195713.59 | 20361987.75 | 25886156.19 | 38106783.47 | 44254923.66 | 39346909.13 | 24870940.09 | 24515068.67 |
| 109955.6537 | 190367.6403 | 238553.4443 | 36665.74586 | 58224.13086 | 185128.9849 | 104458.238 | N/A |
| 140678095.2 | 136292425.7 | 137272887.3 | 109229826.3 | 116850146.3 | 104682315.5 | 145132104.4 | 137503362.1 |
| 606514.2992 | 559258.3378 | 532056.5559 | 397289.3557 | 440831.8801 | 438623.3326 | 563265.0318 | 515095.0911 |
| 753373.2781 | 799783.4739 | 736087.5056 | 790963.6152 | 614465.3573 | 505118.2534 | 1059547.188 | 815660.5944 |

| | | | | | | | |
|---|---|---|---|---|---|---|---|
| 16401705.49 | 15999940.52 | 14519851.53 | 16289175.65 | 17641986.29 | 15765374.86 | 18892783.52 | 19301171.67 |
| 134233897.6 | 130535320.3 | 129688940.8 | 120343606.3 | 118685009.7 | 126543694 | 122928723.9 | 139078658.8 |
| 5419781.072 | 5610248.85 | 6323060.264 | 5501631.659 | 6511387.681 | 5513202.328 | 5298124.466 | 7366080.649 |
| 391870.5937 | 279590.5047 | 374139.3786 | 946109.8224 | 756201.8614 | 766169.858 | 439282.2675 | 479175.0733 |
| 482429.1575 | 579519.6567 | 513282.552 | 735714.3851 | 508856.246 | 707433.4226 | 492014.8579 | 497082.2461 |
| 7630.140777 | 1841.863932 | 7529.450429 | 10006.60876 | 3579.713578 | 7444.876864 | 2492.895078 | 10000.97595 |
| 423905.561 | 567932.0678 | 558002.8254 | 2584978.93 | 2810947.506 | 4562092.51 | 1013557.112 | 788510.1469 |
| 5894226.051 | 7220494.518 | 8902249.145 | 13995118.99 | 13682619.43 | 17564083.4 | 18141448.03 | 10261072.15 |
| 36804.20043 | 35419.93956 | 35751.03807 | 50294.01071 | 19394.82737 | 63488.51122 | 5686.087389 | 30586.12808 |
| 977187.9215 | 738732.5975 | 858706.507 | 1461883.822 | 1299886.456 | 1236865.616 | 1132318.401 | 1140743.881 |
| 2214838.453 | 2441033.074 | 2443889.826 | 2002125.859 | 1868152.015 | 1781007.326 | 1660492.614 | 1510835.714 |
| 27634297.33 | 27474769.78 | 30462746.09 | 18903898.61 | 21583040.19 | 16831077.47 | 20378114.86 | 17722114.67 |
| 49149320.49 | 56358080.09 | 63775651.94 | 50472061.72 | 44566138.31 | 49853741.04 | 49171078.33 | 33347736.09 |
| 3775633.239 | 4183998.345 | 4136495.129 | 3983196.195 | 4028151.601 | 4430349.179 | 4728670.767 | 4951106.342 |
| 6152326.696 | 6540263.084 | 6365335.559 | 5412880.915 | 6702491.474 | 4903988.991 | 5453198.98 | 6101667.948 |
| 22732.82332 | 14090.55771 | 22720.78359 | 47528.67406 | 55598.17946 | 23873.69763 | 12846.90357 | 35155.69603 |
| 54646137.07 | 52627627.17 | 54363318.51 | 28429280.54 | 29002477.54 | 33171439.35 | 40864325.88 | 44316892.79 |
| 43585634.38 | 45026387.42 | 41808134.92 | 40751808.86 | 37574492.43 | 35529953.06 | 32944555.06 | 31820813.09 |
| 162571.3801 | 216899.8979 | 225762.893 | 228482.3376 | 272988.6189 | 197751.6572 | 131813.7071 | 125050.1554 |
| 2758.425745 | 3361.211146 | 13168.26836 | 8362.125744 | 3294.194187 | 3328.299417 | 2509.885245 | 4181.266584 |
| 166827.1139 | 275631.2455 | 272480.5625 | 211336.6354 | 196297.84 | 233438.0215 | 173088.2501 | 179602.761 |
| 13626613.86 | 20481605.34 | 18890227.37 | 11048532.14 | 7542533.259 | 6707347.418 | 10836127.04 | 9378736.971 |
| 13559.76619 | 8065.595051 | 53446.30162 | 5760.267365 | 12448.02336 | 11681.48779 | 10498.24487 | 4291.105059 |
| 2511.304745 | 2509.640215 | 4186.140117 | N/A | 1689.739319 | 9206.104927 | 3348.903363 | 1674.156715 |
| 1035836.286 | 37287.76125 | 33126.76215 | 29561.67717 | 32173.67888 | 25312.79612 | 37236.6474 | 21134.80831 |
| 116509.6049 | 120197.9191 | 119575.5025 | 119635.7587 | 198264.9137 | 72716.48872 | 107922.6435 | 73603.6579 |
| 90439.85869 | 150189.6769 | 104013.1973 | 55737.01662 | 105093.711 | 114890.6359 | 45256.46518 | 246747.1143 |
| 1897675.552 | 2370474.836 | 2090146.81 | 2339868.51 | 2103716.384 | 1745763.207 | 2111906.137 | 1249597.73 |
| 165687.937 | 179880.6767 | 193517.0629 | 372029.2324 | 424088.6864 | 394830.9398 | 234273.0268 | 231879.7564 |
| 213849.7793 | 238967.1431 | 218691.3019 | 295846.8923 | 313505.4093 | 270155.1141 | 179194.3779 | 147605.2529 |
| 644271.0935 | 639272.6468 | 570853.7643 | 596393.4693 | 487599.2859 | 493752.7662 | 569065.9345 | 496334.7106 |
| 160498.295 | 142109.549 | 171430.8717 | 190729.1879 | 185508.7765 | 193093.2982 | 83524.41109 | 74361.04371 |
| 108327.6834 | 141535.5165 | 128296.9535 | 130143.2146 | 128882.0977 | 149548.8695 | 133343.8273 | 166830.8963 |
| 33221.70355 | 37449.57958 | 21425.91335 | 66863.85837 | 98770.73649 | 56666.36946 | 33517.65404 | 11280.22165 |
| 7753.117068 | N/A | N/A | 1674.077232 | 2509.901485 | 6691.337013 | 1691.202303 | N/A |
| 190019.425 | 159402.6312 | 149009.3694 | 167696.5307 | 166484.808 | 97254.01725 | 146982.7531 | 131260.7716 |
| 596837.2294 | 689570.6127 | 753609.2884 | 492149.1548 | 494491.6325 | 521969.8213 | 486802.5977 | 491155.6036 |
| 256129.1165 | 304914.993 | 368647.4695 | 332427.2708 | 360103.3297 | 330694.2007 | 269051.4763 | 238208.3737 |
| 13075.58414 | 7935.245328 | 13101.49921 | 20701.37281 | 5810.480088 | 8405.926589 | 19020.59369 | 6913.632256 |
| 355633.178 | 258626.137 | 265690.7913 | 41874.14929 | 32906.7692 | 28780.95276 | 82683.9015 | 101041.7503 |
| 52013.93769 | 76111.34452 | 38698.17224 | 10038.78132 | 7528.274114 | 6709.096842 | 14223.88006 | 21750.61175 |
| 850570.7559 | 882920.8498 | 1027709.887 | 1472299.629 | 1255960.646 | 1381091.915 | 914293.9401 | 856891.6257 |
| 24654569.4 | 20606237 | 25387040.32 | 33222736.74 | 37704389.49 | 30327857.91 | 25489791.18 | 32495941.22 |
| 381860.3356 | 500825.1128 | 562212.5336 | 327370.7279 | 231938.8253 | 414964.5161 | 282523.6539 | 192505.2443 |
| 3118227.965 | 3843242.437 | 3501472.876 | 2864124.846 | 2210450.534 | 2417648.349 | 2524179.967 | 1757771.16 |
| 66219.84693 | 82283.37944 | 52917.18331 | 48697.93264 | 77593.21975 | 159726.7283 | 42134.3759 | 41106.48268 |
| 12371.70445 | 6706.796924 | 7134.720152 | 9165.411109 | 6607.588646 | 4180.971379 | 21708.50348 | 5837.238719 |
| 27740.79777 | 32203.9223 | 22979.84018 | 99164.08567 | 65236.99268 | 64487.56209 | 65289.34568 | 40359.06534 |
| 1807163.285 | 1473533.84 | 1773712.279 | 2356467.826 | 2356953.356 | 2256081.187 | 1537717.914 | 1251768.249 |
| 79280.18258 | 66552.97584 | 60204.70158 | 66651.56202 | 61586.06589 | 62739.19462 | 37072.51811 | 64444.41616 |
| 22192619.59 | 22044727.64 | 24743800.5 | 37904488.46 | 40714942.6 | 41679660.03 | 27929823.63 | 25269156.3 |
| 403685.8682 | 374833.1173 | 372086.025 | 363424.3073 | 405463.0897 | 1277572.87 | 1035864.054 | 1042085.43 |
| 7258566.233 | 6426303.421 | 6427387.869 | 5372798.431 | 5878222.966 | 5432701.064 | 4525071.644 | 4522324.215 |
| N/A | N/A | 1656.892457 | N/A | N/A | 5855.519379 | 2475.679821 | 1674.157287 |
| 1455637.02 | 1334156.894 | 1248821.018 | 898316.8001 | 640168.4632 | 789757.5035 | 845266.4431 | 592855.747 |
| 96082951.91 | 94028872.18 | 95835112.78 | 102452863 | 91158516.69 | 78801374.37 | 127095820.8 | 115189732 |
| 4244940.963 | 4486462.237 | 4036953.327 | 5631523.721 | 4740247.498 | 6008812.101 | 5826214.973 | 5212136.476 |
| 324489.0721 | 486139.9878 | 328673.5019 | 652631.2876 | 618847.867 | 510132.8098 | 370445.7843 | 294188.2664 |
| 150160.9818 | 228912.1886 | 221602.7632 | 282659.775 | 232913.0252 | 296159.8388 | 160102.8781 | 107088.3299 |
| 1940658.737 | 1711203.287 | 1859916.956 | 1632302.731 | 1721136.447 | 1716329.428 | 1320807.225 | 1970406.978 |
| 139149.8968 | 169815.7249 | 154858.7765 | 405171.4857 | 405392.3181 | 454613.2014 | 143386.7317 | 172362.7372 |
| 82783.17986 | 111178.1781 | 108297.8885 | 141559.3101 | 139859.793 | 149048.3354 | 102541.436 | 86216.13798 |

| | | | | | | | |
|---|---|---|---|---|---|---|---|
| 19231.12767 | 19262.99976 | 19227.55906 | 24940.53554 | 29320.1451 | 20912.42661 | 9982.377391 | 5050.681728 |
| 79591.58454 | 51823.39195 | 56590.3878 | 99641.00211 | 114527.466 | 77092.12925 | 132077.3776 | 114699.2767 |
| 553768.2719 | 470194.7381 | 462694.2377 | 732022.3252 | 683498.8446 | 346360.5184 | 356406.8952 | 353986.1877 |
| 11115.06162 | 11655.32941 | 2518.500084 | 16252.0457 | 15642.66895 | 6431.201917 | 16233.90186 | 11209.86199 |
| 116137.1246 | 75812.31571 | 105836.3973 | 96415.3906 | 128179.0086 | 84420.39583 | 157697.1756 | 124396.5416 |
| 4184.578175 | 15477.06966 | N/A | 5742.945086 | 7543.613589 | 6709.521698 | 2525.575412 | 3345.754556 |
| 4640166.986 | 4311107.436 | 4230115.122 | 4253188.615 | 4623188.004 | 4226589.516 | 1361846.297 | 1365480.368 |
| 355151.0026 | 460193.3765 | 452012.2984 | 2106652.306 | 2297683.993 | 2296549.023 | 729616.2863 | 721609.9195 |
| 4409801.907 | 5285329.403 | 4950859.295 | 5263177.893 | 4440161.775 | 5446801.086 | 5297635.779 | 4014025.326 |
| 765323.7439 | 923832.5275 | 1040693.332 | 1529869.264 | 1188830.657 | 1138094.902 | 1128854.716 | 1230023.722 |
| 85856.76633 | 92353.08212 | 146983.1757 | 119796.0054 | 160018.7957 | 86354.1105 | 60084.21862 | 54593.45056 |
| 61858.64344 | 53664.4817 | 42467.80869 | 53111.33451 | 53143.5605 | 43836.13712 | 36438.85356 | 38448.06871 |
| 11634.31837 | 8064.47976 | 8736.784212 | 42158.53443 | 11976.49567 | 7490.31684 | 10618.34119 | 17458.59516 |
| 9244816.691 | 10722366.07 | 8815283.408 | 13369131.34 | 14898883.68 | 11340571.47 | 8405299.848 | 8217423.609 |
| 352346.5017 | 402122.0044 | 496676.4314 | 1936820.169 | 2253981.193 | 2091723.003 | 804728.3365 | 664218.0164 |
| 60333424.6 | 56969286.87 | 57970443.25 | 55063204.01 | 57673501.04 | 52818887.86 | 61843235.8 | 60688165.29 |
| 5500622.539 | 5132204.477 | 4740836.955 | 3821807.53 | 3824515.941 | 3285056.294 | 5984886.76 | 5473470.89 |
| 244196.4942 | 238362.8217 | 191291.0718 | 194540.8129 | 205700.7275 | 175998.2753 | 216070.5363 | 167289.3006 |
| 2220703.904 | 2392660.945 | 2269503.561 | 2537936.671 | 2375297.792 | 2150064.403 | 2115339.681 | 2431631.539 |
| 148531.9432 | 114790.6969 | 141184.4716 | 131120.8958 | 112760.6762 | 142998.4685 | 110944.9839 | 151239.7864 |
| 118060.6482 | 93299.81657 | 75094.69234 | 111586.8552 | 100338.8274 | 144349.4487 | 148570.7165 | 76797.6657 |
| 26961242.69 | 29433488.78 | 27161599.65 | 36880436.31 | 31852428.89 | 36246726.11 | 21882600.71 | 17306338.85 |
| 132003349.2 | 129220504.3 | 126083603.1 | 161711956.8 | 159696122.8 | 159486447.7 | 159714834.2 | 155417491.1 |
| 5019.922663 | 7526.777574 | 6742.895567 | 1671.900687 | 1672.761436 | 7529.474543 | 5855.102605 | N/A |
| 59172.66922 | 53336.79267 | 57919.1674 | 59254.66811 | 67249.75419 | 66709.64096 | 46779.74088 | 62144.7838 |
| 126806.0617 | 129736.0947 | 112635.1368 | 130998.9327 | 92201.50168 | 137592.1951 | 92902.11568 | 111107.5069 |
| 15040740.3 | 16297377.97 | 13526735.49 | 15385462.64 | 14290868.35 | 10697365.24 | 18671980.23 | 19536423.64 |
| 962648.4342 | 894106.0147 | 950944.7096 | 1582911.871 | 1650418.019 | 1320282.822 | 1023591.866 | 1139118.462 |
| 34986657.51 | 33689433.25 | 31061485.52 | 37305346.21 | 33427331.92 | 37400278.05 | 30641557.83 | 30573612.91 |
| 1533227.783 | 1439152.068 | 1249594.954 | 1655729.354 | 1659278.261 | 1327078.915 | 1404565.867 | 1358171.304 |
| 494634.4487 | 603121.0345 | 415853.7009 | 409221.3187 | 378003.5671 | 354127.9181 | 447597.0872 | 391320.4947 |
| 78299.4321 | 66822.30128 | 74002.92962 | 74955.91497 | 87769.28907 | 91725.6764 | 34445.93778 | 38191.2913 |
| 26331356.93 | 21320609.41 | 33082575.36 | 48649588.86 | 47526664.82 | 45897371.42 | 21393203.4 | 25308110.47 |
| 6385800.317 | 7481389.855 | 6722288.856 | 7902904.46 | 8199043.273 | 4923442.385 | 6520896.496 | 5862915.256 |
| 8365.105331 | N/A | 15860.34491 | 11555.60329 | 10068.17984 | 2495.060967 | 13397.70647 | 6556.766908 |
| 14240.17259 | 10840.9264 | 8367.599053 | 13384.2073 | 15874.50599 | 5858.000725 | 3314.459844 | 6639.765123 |
| 134055.4967 | 178233.468 | 161970.5506 | 98080.56385 | 95711.18576 | 75403.64033 | 124367.8186 | 97367.21567 |
| 63254.89348 | 62893.32603 | 78223.09369 | 55904.11593 | 69640.62572 | 50588.11358 | 67038.67574 | 84773.09761 |
| 65428.93083 | 71447.02326 | 49617.62775 | 63969.53175 | 80601.95293 | 67709.53671 | 57967.97291 | 46743.50663 |
| 212195368 | 197951728.5 | 196801110.4 | 238254217.5 | 75663678.02 | 183128587 | 223741256.3 | 226510537.3 |
| 302187.3182 | 318808.6473 | 258051.9907 | 281131.3336 | 276891.4702 | 282622.4902 | 272136.4084 | 301397.1359 |
| 2121.992513 | 25719.54126 | 4480.239686 | 2805.274573 | 14788.01875 | N/A | 14416.16602 | 12935.35748 |
| 37264025.65 | 36768498.42 | 41294631.12 | 33525262.11 | 30722947.43 | 30360348.87 | 33598769.35 | 34418779.79 |
| 52011181.77 | 55401872.97 | 56326496.04 | 45011671.68 | 44421502.7 | 40846559.71 | 38495680.69 | 34340770.77 |
| 3111282.493 | 2824624.263 | 3572717.243 | 4843005.525 | 5432163.758 | 3542597.709 | 3379962.534 | 3956550.814 |
| 218268.2602 | 269685.1661 | 169486.0839 | 243123.3743 | 196887.6394 | 214073.4525 | 149574.8373 | 147423.675 |
| 9558129.647 | 8955955.182 | 8964035.704 | 10595223.88 | 10820222.32 | 8181695.921 | 10424727.3 | 11617268.07 |
| 1033489.207 | 912993.4053 | 910206.8148 | 824193.3203 | 832812.5809 | 648923.736 | 858686.6592 | 827259.345 |
| 1758217.492 | 1561360.057 | 1644379.799 | 1936610.349 | 1870280.321 | 1946188.887 | 3368827.712 | 3880431.557 |
| 547733.9487 | 658128.75 | 794298.5709 | 984426.7861 | 924036.7833 | 923613.924 | 708533.5487 | 849263.1618 |
| 9390630.651 | 10428452.03 | 10391842.28 | 8255415.625 | 6558545.59 | 7148938.852 | 8010363.408 | 8294191.412 |
| 397980.6756 | 441046.3885 | 570990.6637 | 246156.2069 | 256540.267 | 309889.6892 | 189733.37 | 158880.8031 |
| 957934.979 | 993764.1144 | 903985.0257 | 899386.7321 | 810241.452 | 953050.5896 | 899413.1528 | 720990.3296 |
| 2149506.37 | 2379017.526 | 1994416.645 | 2225582.769 | 2111480.045 | 1834021.817 | 2328863.58 | 2075505.271 |
| 302361.0844 | 364237.0584 | 247085.8001 | 207489.4048 | 286109.7115 | 209090.8663 | 182817.6135 | 154283.7665 |
| 69844993.51 | 69991928.08 | 70089486.14 | 62281904.58 | 65116764.92 | 62564683.87 | 66016225.56 | 62689979.05 |
| 105045.1369 | 135324.9216 | 89014.32298 | 141299.2413 | 127583.6914 | 92959.43991 | 123077.0512 | 103909.328 |
| 237473790.7 | 241445533.1 | 243354810.5 | 237904647.6 | 229403937.9 | 223659433.3 | 232845176.7 | 233168895.4 |
| 2511.11287 | 4228.92612 | 11627.42752 | 2509.794118 | 9183.318424 | 3361.977191 | 6687.635767 | N/A |
| 41688363.43 | 34776958.64 | 37017604.07 | 33959329.81 | 32139651.97 | 30020179.87 | 31514930.17 | 32085367.11 |
| 29512633.04 | 27581130.57 | 26205364.43 | 25792135.39 | 25761102.88 | 24993571.38 | 22595767.45 | 25270148.19 |
| 4398218.327 | 4713835.054 | 4897800.83 | 5019581.828 | 4029557.445 | 4881401.855 | 4647343.885 | 4112672.939 |

| | | | | | | | |
|---|---|---|---|---|---|---|---|
| 5148.074487 | 3332.194285 | 22651.18105 | 26882.27885 | 22375.32713 | 12948.55323 | 4167.522951 | 21758.81451 |
| 26040748.14 | 28633695.17 | 27325591.27 | 23825095.1 | 24184104.63 | 21641350.36 | 28950703.75 | 29019729.74 |
| 5804049.294 | 4390971.354 | 5835866.12 | 7106946.453 | 4807056.881 | 5010869.823 | 4516294.442 | 3812253.266 |
| 3886980.546 | 3317451.837 | 3457121.935 | 3541677.606 | 3033281.71 | 2566092.302 | 3309921.549 | 3498310.505 |
| 135861138.2 | 128614201.2 | 134003840.5 | 132681241.9 | 133325065.3 | 121389053.6 | 127726373.2 | 135031287.2 |
| 16885.57611 | 6239.58942 | 11892.26079 | 21062.12006 | 5233.151948 | 5374.434718 | 12683.6202 | 9603.849966 |
| 152054347.2 | 157760488.4 | 151763761.4 | 119796645.8 | 128595621.7 | 123338034 | 115341505.2 | 118466288 |
| 91869928.84 | 114338110.6 | 108486401.5 | 99905700.28 | 115660509.9 | 126473360.8 | 119873178.6 | 110330435.7 |
| 33230.73758 | 23913.53176 | 9356.119709 | 11547.23188 | 9728.175034 | 4842.317841 | 9382.176174 | 34930.69025 |
| 86931.79 | 95994.04354 | 97817.73579 | 136238.1327 | 78543.55813 | 76563.06354 | 117918.2181 | 74224.24736 |
| 29145.59608 | 37730.28706 | 12140.38631 | 33057.14835 | 19242.22743 | 13385.49948 | 28444.81652 | 13251.08058 |
| 42780.04634 | 21007.29849 | 29274.13939 | 41634.19749 | 57788.85519 | 36171.67294 | 32994.32997 | 27570.84914 |
| 345317.6635 | 273455.5245 | 273530.4035 | 341947.4525 | 155514.9112 | 117903.0861 | 240937.5754 | 638313.737 |
| 488975.7252 | 451198.1234 | 370261.9102 | 424077.5019 | 410232.0912 | 356145.3285 | 413387.7094 | 413298.3409 |
| 49106.95961 | 52532.90835 | 49771.34371 | 46706.87561 | 47301.95173 | 68995.8801 | 56204.60052 | 14736.40084 |
| 43500280.84 | 38734973.1 | 40784195.16 | 22502369.17 | 21923486.85 | 24621951.25 | 29818136.6 | 32553540.46 |
| 6639.957121 | 12585.02887 | 15741.24905 | 5937.460353 | 10871.95027 | 6689.920692 | 3330.946625 | 4159.526978 |
| 4781.049366 | 6521.39644 | N/A | 6109.409459 | 6653.4866 | 10957.04748 | 6184.728382 | 17270.79703 |
| 10644.81002 | 30635.5089 | 16868.48831 | 13720.24499 | 16260.65391 | 29698.60324 | 6023.079253 | 12349.91825 |
| 46067906.56 | 48322010.58 | 49602946.86 | 53072838.22 | 53306739.33 | 52329426.92 | 50358403.77 | 50877578.49 |
| 66925.30335 | 60247.48729 | 60245.81137 | 87863.71335 | 63581.39345 | 60228.97481 | 2422.601866 | 65293.80493 |
| 3374304.391 | 3125128.08 | 3157320.493 | 3811786.516 | 3350027.36 | 2907428.673 | 2613290.657 | 3004371.827 |
| 26768.29731 | 24244.43374 | 9297.67246 | 35133.62271 | 39317.8628 | 30115.34508 | 30951.73427 | 16732.32354 |
| 430605.5367 | 271157.4428 | 371668.4497 | 717975.5316 | 698464.2822 | 644014.1314 | 296206.3191 | 313021.1431 |
| 42256101.42 | 43366281.25 | 41684564.84 | 43438539.97 | 35418422.03 | 48362610.45 | 43702382.03 | 38743223.49 |
| 2179825.29 | 2074277.91 | 1588389.932 | 950547.861 | 1024667.999 | 427425.6094 | 1283485.999 | 2491874.621 |
| 1714.119554 | 34366.22385 | 12900.87971 | 18211.10126 | 3851.039958 | 57669.61082 | 16769.99104 | 25868.48217 |
| 6107524.929 | 6459428.252 | 7273772.877 | 737642.144 | 691812.1615 | 618617.7481 | 1509542.843 | 1412747.396 |
| 701994.2744 | 684817.7181 | 660567.1891 | 803059.9038 | 682730.53 | 698752.8952 | 708446.409 | 606730.0983 |
| 994900.7057 | 786194.0939 | 898550.782 | 2301364.41 | 1873600.439 | 2678835.579 | 1412442.423 | 1125421.459 |
| 61788.29182 | 37765.15524 | 108036.5376 | 65455.02233 | 78551.23881 | 49680.26316 | 56523.31242 | 47682.3175 |
| 107445.0047 | 158157.4566 | 165407.311 | 291487.8098 | 286679.6779 | 238287.5063 | 103517.2478 | 129238.1061 |
| 21895146.62 | 16910548.68 | 24236095.46 | 22347603.58 | 18065686.79 | 18180114.25 | 16159868.34 | 13453631.19 |
| 461482.0007 | 415678.1765 | 430089.6617 | 243802.3472 | 349230.544 | 313011.079 | 230451.2103 | 166287.6683 |
| 78878.86036 | 141884.5974 | 111000.9043 | 76957.9495 | 119543.5987 | 104065.5629 | 93501.16001 | 114849.4004 |
| 668379.9846 | 669172.7267 | 825158.8685 | 790155.6985 | 1023259.333 | 717069.8935 | 735677.5863 | 995643.0043 |
| 580248.6993 | 465956.5247 | 531732.0684 | 368892.8766 | 468853.2524 | 392367.6202 | 381087.6123 | 480258.5541 |
| 20677377.26 | 20287715.64 | 20198912.61 | 19317936.15 | 17498641.4 | 21798116.2 | 20002150.61 | 16134721.21 |
| 91310366.9 | 87915028 | 85748937.95 | 85206848.35 | 83723180.44 | 81203434.37 | 83431551.99 | 81921729.43 |
| 248729.4495 | 111205.7229 | 299714.2107 | 242250.9199 | 155683.2237 | 115790.6784 | 144757.3105 | 170309.8803 |
| 362248.6086 | 464167.5076 | 336086.0734 | 373545.7037 | 349668.9798 | 304954.9885 | 511862.488 | 441857.5295 |
| 9442.188577 | 22043.45705 | 30961.63851 | 10088.33093 | 28627.0548 | 17676.36318 | 6616.127646 | 29179.14867 |
| 86636.6708 | 137140.3398 | 47346.31683 | 47247.94412 | 66651.94748 | 49708.8236 | 28442.05566 | 81684.675 |
| 2262622.467 | 1911142.295 | 1953827.429 | 5595375.31 | 7957031.968 | 6475554.074 | 1915467.937 | 2638898.324 |
| 68499141.25 | 79145887.18 | 74667498.38 | 49435038.34 | 50259750.22 | 42848848.8 | 40075068.11 | 54406709.58 |
| 149947.5572 | 123718.2326 | 182154.4487 | 199452.3858 | 153150.4266 | 95996.50021 | 131336.5581 | 120968.5241 |
| 3345.375146 | N/A | N/A | 5019.585364 | N/A | N/A | 1668.819952 | 1656.892457 |
| 118227435.8 | 85536636.83 | 78698414.08 | 79472359.26 | 78118554.16 | 73572019.52 | 79256163.33 | 193522459.4 |
| N/A | 1673.599019 | 4179.852067 | 16092.6701 | 14352.66195 | 10039.43862 | 3196.932273 | 3348.085088 |
| 1616404.297 | 1586312.467 | 1287846.892 | 1168975.106 | 1243228.805 | 922102.8214 | 1100013.547 | 1342504.125 |
| 45363.13381 | 92793.73019 | 100950.3729 | 216481.1745 | 119420.1671 | 74690.18393 | 134581.8997 | 22126.12528 |
| 3317.958596 | N/A | N/A | 1674.144208 | 4942.39741 | 3320.882584 | 7520.499936 | 8337.193814 |
| 140836.7689 | 128273.8306 | 172853.4692 | 259404.7419 | 387947.5622 | 320053.5767 | 490603.7631 | 458937.37 |
| 15107921.17 | 16487154.61 | 14744569.9 | 15872760.39 | 13548316.51 | 14981308.72 | 13555830.89 | 13592851.41 |
| 78482317.68 | 76723365.73 | 76868011.87 | 69714319.41 | 70987375.01 | 72951175.14 | 57030047.8 | 61280216.08 |
| 150279.7958 | 175518.6574 | 167477.7335 | 115515.2145 | 145436.6364 | 172346.8127 | 150678.0411 | 149155.3211 |
| 136587.1841 | 74038.86915 | 132935.7412 | 95944.81485 | 109497.8371 | 87030.39207 | 70205.39525 | 99120.88748 |
| 16896.92907 | 26339.05672 | 45066.36738 | 48309.06721 | 35889.61126 | 18939.61634 | 26895.50466 | 9466.452507 |
| 2078091.511 | 1878634.688 | 1979491.98 | 3180735.963 | 2366666.819 | 2781876.656 | 2260287.526 | 1862645.066 |
| 25915002.58 | 23336447.26 | 28752697.81 | 17546184.12 | 19096736.53 | 18977624.22 | 21132445.99 | 21779771.12 |
| 3888970.403 | 3436866.209 | 3613686.473 | 4225096.31 | 4213092.346 | 3559434.688 | 3699631.76 | 3615885.279 |
| 3592.902641 | 8286.498896 | 2476.417349 | 5954.275636 | 4183.931758 | 3361.296533 | N/A | 4180.837942 |

| | | | | | | | |
|---|---|---|---|---|---|---|---|
| 16421635 | 11120849.14 | 12468040.63 | 17017215.84 | 17529181.39 | 14815479.42 | 9931112.66 | 8801119.969 |
| 103789734.4 | 93066192.56 | 95532526.18 | 140314166.2 | 122717193.6 | 121388033.9 | 115624853.7 | 99067827.91 |
| 272243.7038 | 246299.1395 | 219461.9404 | 326459.8453 | 334518.1338 | 283458.9633 | 182417.6464 | 157078.2149 |
| 2461664.386 | 1943625.361 | 2260062.488 | 1944965.347 | 1902636.975 | 1316456.897 | 1873834.775 | 1607184.548 |
| 1998361.994 | 3357034.577 | 3601785.639 | 3201429.537 | 3149726.885 | 2956713.994 | 2422031.275 | 2554126.487 |
| 127654.9271 | 102111.7137 | 101957.4855 | 98473.51641 | 74033.78397 | 69899.96601 | 86553.18794 | 158410.349 |
| 1560392.48 | 1335692.499 | 1002668.206 | 1398461.282 | 1304896.378 | 1407071.771 | 1563027.094 | 1316551.73 |
| 18883150.84 | 17107529.21 | 18423674.95 | 22809585.4 | 25036551.16 | 21829197.55 | 15516374.98 | 14548592.85 |
| 180846.7035 | 61272.90048 | 193056.6723 | 140906.3035 | 147750.4145 | 168450.4844 | 156244.0234 | 183498.2769 |
| 7697.382682 | 9987.847711 | 2989.304026 | 28765.34146 | 35297.67695 | 21427.88221 | 2878.41116 | 28886.34743 |
| 134018.9919 | 167037.6334 | 153735.1394 | 91497.53164 | 107731.3381 | 100283.6638 | 168792.0428 | 145422.192 |
| 136179091.7 | 131634044.2 | 133819518.2 | 129818833.3 | 132134703.8 | 120399246.8 | 127343024.5 | 133067365.2 |
| 31594.29287 | 28357.44849 | 34300.83518 | 66662.57172 | 43350.56281 | 22058.66731 | 59294.28173 | 47669.79202 |
| 7375035.746 | 7181819.236 | 7967037.405 | 7173320.618 | 8071026.963 | 6558108.407 | 5002276.037 | 5702442.765 |
| 811407.3738 | 886440.703 | 793402.4901 | 529329.0646 | 611086.5119 | 865952.2809 | 514761.5645 | 434290.9216 |
| 258067.214 | 140415.8207 | 73893.10593 | 27946.97097 | 48567.79906 | 63521.57939 | 31537.45498 | 34422.57039 |
| 532685.846 | 548487.3901 | 448212.0954 | 908934.1317 | 903118.2334 | 863459.857 | 531990.6508 | 678170.5776 |
| 340304.6116 | 499960.8546 | 570838.476 | 1123070.85 | 795022.3726 | 366846.4656 | 452854.0473 | 693872.7096 |
| 2314301.441 | 1925583.533 | 1999928.862 | 2696924.727 | 2570671.181 | 2167021.537 | 5668467.897 | 5807621.177 |
| 43571.49164 | 50767.79867 | 56206.93651 | 49603.9111 | 47918.475 | 38718.51546 | 22331.47014 | 23103.7904 |
| 33012.07208 | 30748.65774 | 15299.23228 | 123593.5267 | 22966.75793 | 68557.43444 | 24649.22551 | 7617.323257 |
| 33421.23354 | N/A | 80053.8881 | 65546.5786 | 46239.29122 | 121914.7828 | 55534.44606 | 57668.16544 |
| 905906.7674 | 1087248.495 | 863956.3179 | 2137879.738 | 2242003.799 | 984335.0421 | 452411.2617 | 1150952.458 |
| 34769.13115 | 25861.27052 | 11712.37061 | 40612.24967 | 26357.39374 | 40135.48513 | 19058.61876 | 19000.84712 |
| 25089008.74 | 24441853.15 | 23079805.48 | 20658312.62 | 18826224.01 | 18390954.86 | 18484202.57 | 21359690.62 |
| 13189881.11 | 12552474.06 | 9390768.091 | 11946537.98 | 9984056.474 | 8102201.869 | 13529976.14 | 11657135.3 |
| 3268084.361 | 3485129.122 | 3377846.301 | 23189204.93 | 26638910.04 | 22900863.33 | 4034378.332 | 6067088.32 |
| 87003.97801 | 72791.86642 | 86991.92967 | 174843.7819 | 193183.8727 | 162153.6937 | 51032.90647 | 47685.51982 |
| 1305018.128 | 1189659.031 | 1201105.805 | 9057840.886 | 10779253.9 | 6369783.012 | 1058041.003 | 2445578.463 |
| 15766235.44 | 20625544.39 | 18483338.67 | 40254583.59 | 37907681.06 | 34582930.84 | 22432622.07 | 27585843.69 |
| 4029553.307 | 3739414.122 | 3607884.661 | 6566615.994 | 7976071.948 | 4773268.774 | 5778543.899 | 5248764.057 |
| 105651372.4 | 105589073.4 | 102362258 | 142435511.6 | 136185127 | 131784895.1 | 59580157.09 | 54343846.79 |
| 3921484.718 | 3694836.485 | 3532358.874 | 4323625.348 | 3946530.39 | 3316307.102 | 3752352.932 | 3754927.093 |
| 236424.4267 | 327314.5889 | 265401.0688 | 204858.0389 | 191641.6955 | 202757.1059 | 185328.8305 | 154985.1691 |
| 202288.1253 | 248707.5684 | 242916.0374 | 192438.9175 | 236265.2906 | 187852.3994 | 136959.2179 | 158886.767 |
| 13440560.78 | 15403816.82 | 17120909.07 | 32563307.06 | 38299295.37 | 30275986.3 | 16284371.87 | 19435631.01 |
| 12214771.39 | 11086414.01 | 10507017.87 | 12093232.17 | 9323443.08 | 7830613.33 | 11371647.75 | 11940503.96 |
| 54647188.12 | 59754804.68 | 57688013.97 | 53363223.34 | 47537935.79 | 58357390.62 | 47417320.13 | 47131429.74 |
| 1558419.565 | 1964216.051 | 1819993.505 | 2195199.373 | 1797925.371 | 1908851.708 | 1552836.648 | 1604293.491 |
| 14255.87266 | 5019.639468 | 7527.284233 | 24274.84481 | 22621.65557 | 6690.422232 | 9201.114292 | 17566.3275 |
| 1412801.636 | 1618148.488 | 1304026.911 | 1197349.956 | 1371248.639 | 1343712.002 | 1165802.2 | 1135296.033 |

| | elo3::elo3-gfp | | |
|---|---|---|---|
| 60632.1279 | 115335.3001 | 123883.7264 | N/A |
| 3347.158844 | 2510.394647 | 5381.007876 | 8496.338695 |
| 7492190.001 | 5881919.042 | 6136789.389 | 5515733.541 |
| 95653.87334 | 97190.11151 | 181742.9169 | 99706.03152 |
| 29220.73107 | 71521.59142 | N/A | 35815.5852 |
| 365575.9564 | 323355.9696 | 429463.0882 | 249069.9026 |
| 142212.8757 | 147216.0647 | 158031.4077 | 155931.612 |
| 1047460.485 | 1574015.306 | 1418079.764 | 1503200.321 |
| 1356454.694 | 1189558.679 | 1209574.452 | 1374831.159 |
| 7838593.574 | 7655451.736 | 9493978.511 | 7180098.216 |
| 340622.2031 | 331016.5708 | 332383.1167 | 320306.827 |
| 238851.6331 | 169937.8201 | 231590.0925 | 271391.6749 |
| 3803841.511 | 4274140.534 | 4446757.831 | 4972885.423 |
| 19476.25979 | 13629.31589 | 13020.33566 | 32379.98492 |
| 351956.7388 | 369781.6232 | 361825.9843 | 314133.0016 |
| 14640620.12 | 12182993.02 | 11398787.25 | 11828023.45 |
| 318269.7671 | 371254.9673 | 535495.7611 | 754104.477 |
| 5461390.125 | 6316093.75 | 7016391.67 | 7240051.605 |
| 8899.552522 | 9305.373042 | 10598.43127 | 6677.665948 |
| 7441.937968 | 606693.375 | 602175.5209 | 13056.13927 |
| 220831.0607 | 220827.9087 | 370096.8107 | 223004.9929 |
| 225962.6619 | 164863.8435 | 176803.3591 | 174730.8626 |
| 83866.06729 | 115712.5058 | 186709.8118 | 76034.51927 |
| 10824.59023 | 11695.86017 | 18025.36837 | 10822.43833 |
| 19241.32802 | 33427.94767 | 24911.45421 | 8479.907769 |
| 10947.29438 | 124693.8486 | 133195.9959 | 5808.511434 |
| 67347.75958 | 73834.91879 | 77397.44231 | 79772.92083 |
| N/A | N/A | 1674.155371 | 3344.869138 |
| 372122.2066 | 269476.2865 | 275969.9527 | 229503.0821 |
| 118701.0608 | 156418.8612 | 93294.49471 | 128835.8908 |
| 296229.3002 | 363784.15 | 286129.8368 | 295461.4264 |
| 260553.2143 | 377565.5632 | 356325.1299 | 231434.2985 |
| 79536280.98 | 74022810.88 | 89626462.04 | 86102866.67 |
| 329085.6615 | 395047.5447 | 412712.9685 | 293085.5757 |
| 91129420.33 | 110856621.8 | 99919113.21 | 101403287.1 |
| 519627.4263 | 651337.5791 | 634139.3573 | 624996.4442 |
| 409052.9253 | 587664.3785 | 632455.5463 | 528098.297 |
| 1466867.455 | 2743775.767 | 2859131.218 | 2016911.504 |
| 1501308.83 | 2005946.97 | 1896206.786 | 1572740.314 |
| 347943.9809 | 457153.4805 | 420300.2975 | 431439.0943 |
| 4216.23702 | 12548.74185 | 1644.897597 | 1676.179673 |
| 59398.46561 | 56920.79791 | 92046.49631 | 68190.38523 |
| 28487223.71 | 36215918.98 | 34461031.12 | 37784260.32 |
| 226399.0522 | 301635.6748 | 325540.5797 | 292739.501 |
| 114703923.2 | 140933557.8 | 137163279.6 | 136848969.8 |
| 186883.7887 | 173560.2826 | 183405.519 | 144128.6621 |
| 2149872.927 | 2528180.059 | 2736472.075 | 2235867.984 |
| 5661118.177 | 7194570.271 | 6744045.725 | 6314406.325 |
| 124163.3397 | 117143.9333 | 6934.649602 | 80537.33091 |
| 79379184.46 | 69294860 | 91794847.78 | 69425667.14 |
| 418399.5454 | 428520.5024 | 441277.6391 | 348138.5707 |
| 209146228.4 | 201355537.2 | 209626661 | 205189355.5 |
| 110881.1522 | 66720.44126 | 71471.88457 | 81720.72515 |
| 287729.1107 | 396414.2255 | 411459.041 | 387387.1322 |
| 2722745.6 | 2680248.607 | 2969719.911 | 2644201.885 |
| 5943168.106 | 4707288.219 | 6421524.7 | 6520091.105 |
| 24406274.58 | 32065515.11 | 31586625.87 | 33294571.81 |
| 33782.65533 | 208847.688 | 89702.97595 | 163310.7641 |
| 130830716 | 135692798.1 | 134431208.5 | 124637927.8 |
| 363677.9761 | 479441.4517 | 521687.0374 | 498875.6559 |
| 976112.4832 | 667034.5273 | 911433.5356 | 681836.3924 |

| | | | |
|---|---|---|---|
| 17508808.34 | 20422622.68 | 19450206.07 | 18854960.31 |
| 136399327.3 | 139173791.3 | 130650124.7 | 129296739.3 |
| 6558115.922 | 5043955.713 | 5220343.182 | 4762242.795 |
| 399010.5194 | 562717.2504 | 588724.4889 | 452097.2566 |
| 465314.0114 | 595822.1279 | 684340.8701 | 592274.8029 |
| 3346.33821 | 7531.200012 | 4100.084246 | 1672.615601 |
| 1139492.956 | 598182.5316 | 684003.9704 | 951523.7726 |
| 18662458.79 | 15379027.52 | 14798016.39 | 20379233.83 |
| 12899.74403 | 36744.44726 | 41973.30316 | 35836.95241 |
| 1203321.595 | 1759909.396 | 1763024.958 | 1769662.469 |
| 1478865.524 | 1641532.171 | 1669989.14 | 1952627.302 |
| 13588038.61 | 22430945.03 | 20756551.19 | 23261220.61 |
| 41516804.61 | 46445094.82 | 44521141.59 | 50190977.37 |
| 3939084.381 | 4740845.286 | 5110512.546 | 5012339.646 |
| 5424203.128 | 3911215.503 | 4728754.585 | 4057915.488 |
| 18385.57976 | 54309.2299 | 42672.36316 | 23328.04576 |
| 41982967.14 | 29878670.02 | 34083523.04 | 33400937.62 |
| 24984547.96 | 32303167.92 | 35348456.74 | 30280245.01 |
| 122385.7357 | 207272.2847 | 176241.626 | 198971.1677 |
| 1675.414955 | N/A | 7531.71788 | 3312.77124 |
| 173431.9463 | 269849.3903 | 294961.3533 | 209971.3842 |
| 9186615.268 | 6190811.435 | 6908293.382 | 6534782.803 |
| 5780.322804 | 8978.50188 | 72684.77414 | 4241.00474 |
| 5019.379301 | 3346.351396 | 4113.323196 | 5854.026875 |
| N/A | 16577.64854 | 32725.30696 | 2787.382957 |
| 69435.74034 | 46014.67162 | 61065.46439 | 51884.45987 |
| 150870.6052 | 59144.15584 | 78198.99531 | 39581.65887 |
| 1727177.091 | 1606216.488 | 2658022.707 | 1483572.039 |
| 227997.26 | 228473.9545 | 263253.225 | 268858.7782 |
| 135035.7389 | 254358.8905 | 221979.9247 | 268172.1803 |
| 497018.5162 | 361535.6546 | 423444.6723 | 350178.6916 |
| 67059.18141 | 127888.0983 | 157899.2482 | 124591.8335 |
| 105134.4353 | 141291.1603 | 164541.3732 | 164361.509 |
| 34148.26056 | 33213.95461 | 26158.16909 | 43983.33015 |
| 2510.510172 | 5020.316998 | 1674.931052 | 1691.096268 |
| 133837.6529 | 160560.3772 | 212354.1232 | 110478.9384 |
| 494624.6771 | 717988.4234 | 777955.2329 | 654420.0531 |
| 269774.954 | 283489.1943 | 302652.6925 | 330239.6239 |
| 4889.31101 | 11762.90258 | 11641.69353 | 21382.87103 |
| 85212.76777 | 319427.3224 | 372133.9731 | 393633.5395 |
| 10877.6888 | 23520.45396 | 47659.42479 | 20821.28945 |
| 903487.8105 | 915380.0308 | 1144914.251 | 1040826.082 |
| 25699543.39 | 39770035.87 | 38004668.69 | 36922111.66 |
| 270677.7134 | 180254.9441 | 314846.6415 | 255656.1929 |
| 2441422.783 | 1851959.958 | 2337875.313 | 2217075.786 |
| 53245.77538 | 53316.13486 | 78607.14747 | 56459.7167 |
| 6799.208497 | 9566.403253 | 1712.820108 | 4116.971128 |
| 39618.97335 | 21741.0242 | 53368.04698 | 21994.33581 |
| 1176512.881 | 1592552.731 | 1485612.377 | 1779472.374 |
| 59381.7132 | 31777.23994 | 40900.5749 | 47666.60102 |
| 23940439.85 | 32354896.18 | 31960420.08 | 36859746.37 |
| 300626.7188 | 306286.8979 | 449784.2017 | 1466397.372 |
| 5310044.424 | 6268505.922 | 6198417.543 | 6933192.898 |
| N/A | 3951.525449 | 10877.88966 | N/A |
| 841951.9489 | 822430.5142 | 855845.7273 | 762815.1651 |
| 118366843.1 | 120090785.6 | 126388460.8 | 115014032.6 |
| 5476287.607 | 6022181.625 | 8620924.54 | 5379269.698 |
| 339147.6021 | 426768.8794 | 525438.7524 | 486615.2981 |
| 144718.6458 | 73613.78285 | 200057.049 | 116993.7761 |
| 1714618.926 | 2096627.95 | 1361665.188 | 1815831.575 |
| 144753.1656 | 207062.008 | 302448.5188 | 175942.1065 |
| 76068.94178 | 93544.09563 | 85095.89907 | 90932.57227 |

| | | | |
|---|---|---|---|
| 15059.49936 | 21745.66041 | 24677.64204 | 13370.37723 |
| 125514.6372 | 133935.7033 | 114269.9612 | 94860.32453 |
| 240082.4616 | 456734.2034 | 437485.9324 | 299508.7332 |
| 7905.323686 | 18797.5627 | 18056.9642 | 9464.263228 |
| 144535.3864 | 154241.1321 | 98774.33385 | 195724.1015 |
| N/A | 2508.339202 | 3745.810736 | 1641.96538 |
| 1170899.764 | 3489804.026 | 4713396.491 | 3200302.525 |
| 699576.4618 | 780046.99 | 834703.0881 | 652876.1757 |
| 4462754.855 | 5031929.567 | 5140040.04 | 4844085.212 |
| 1031560.67 | 1572652.896 | 1441708.456 | 1691281.183 |
| 7033.071891 | 15678.38472 | 193225.1828 | N/A |
| 28977.34376 | 28717.82418 | 36963.80882 | 29449.16225 |
| 25374.39276 | 40048.69904 | 46079.20268 | 18569.46043 |
| 7364632.569 | 10559845.91 | 9535390.055 | 10404663.41 |
| 681511.5159 | 588035.6341 | 768423.307 | 570048.0328 |
| 59020480.91 | 61041582.22 | 61004851.96 | 61562871.11 |
| 5680827.835 | 6067448.671 | 6916539.107 | 6529222.77 |
| 193099.411 | 140788.2898 | 209266.5189 | 112216.2615 |
| 2344025.373 | 1572454.444 | 1924653.344 | 1587045.848 |
| 159045.2791 | 130639.6905 | 113382.67 | 110584.782 |
| 108431.6133 | 159804.946 | 116578.8674 | 134290.5074 |
| 19546517.31 | 28027451.18 | 30388318.42 | 25869201 |
| 153908638.9 | 178109614.7 | 184042279.4 | 166034896.2 |
| 5019.554043 | 4181.267166 | 11711.14764 | 3346.411687 |
| 66786.99954 | 38670.67681 | 62398.75605 | 50738.53328 |
| 99547.34624 | 117070.5363 | 113693.0012 | 111623.4993 |
| 19384703.08 | 12240429.72 | 16643802.18 | 10477792.52 |
| 1133217.672 | 1000517.865 | 1192619.224 | 1270728.059 |
| 27698989.98 | 30058897.46 | 32774439.98 | 23789632.87 |
| 1433637.869 | 1087657.86 | 1457084.133 | 1177194.783 |
| 633693.3262 | 363120.9813 | 494881.8879 | 276950.3647 |
| 159802.8268 | 54578.98939 | N/A | 75213.90621 |
| 23824409.45 | 27790827.65 | 31942106.27 | 23023478.42 |
| 6069365.315 | 7584581.696 | 8011609.435 | 6810420.908 |
| 9151.987331 | 4181.293027 | 16727.55031 | 6724.57264 |
| 5520.659916 | 10019.83419 | 9195.164512 | 8466.789573 |
| 79429.14688 | 81549.60374 | 120283.2024 | 77236.8678 |
| 56500.87541 | 35575.19154 | 45747.60967 | 51606.26555 |
| 71783.75428 | 53984.89503 | 66398.97784 | 65886.86171 |
| 256881285 | 189626801.2 | 305651927.1 | 185882811.2 |
| 253046.5202 | 311862.6885 | 214776.8229 | 205858.7845 |
| 3335.956906 | 6941.082812 | 13559.92263 | 5263.007589 |
| 36386712.16 | 31121287.55 | 34527542.88 | 27043274.21 |
| 33357179.95 | 39380466.84 | 43685832.74 | 41115796.08 |
| 2700582.57 | 5493019.939 | 5369217.269 | 4442120.491 |
| 197038.8965 | 154479.6793 | 201433.527 | 127359.9622 |
| 10854901.97 | 6872930.58 | 9427017.351 | 8808311.256 |
| 806064.6406 | 799400.2724 | 874003.9825 | 606758.8018 |
| 3429349.431 | 4030704.449 | 3595708.657 | 3972440.807 |
| 806099.4293 | 949134.5393 | 970657.5889 | 1173227.761 |
| 7538359.683 | 10784876.86 | 9464313.097 | 9139274.076 |
| 263209.1987 | 162667.2354 | 244283.4788 | 224609.7243 |
| 795149.3921 | 919322.5037 | 875989.6537 | 949285.4226 |
| 3079262.837 | 2233124.818 | 2566268.035 | 2196362.224 |
| 190117.2485 | 189165.1123 | 169047.9921 | 163312.9657 |
| 60817170.82 | 68983239.87 | 66363930.08 | 68080069.68 |
| 140876.5479 | 58727.8309 | 85343.12982 | 45890.01237 |
| 231617823.5 | 207621581.8 | 240769954.9 | 204646411.4 |
| 6787.233745 | 4177.982471 | 2508.225035 | 4183.951405 |
| 30373404.89 | 35576289.85 | 32145595.78 | 36540605.82 |
| 24073672.55 | 32174058.08 | 26359293 | 32054256.77 |
| 4096521.679 | 4536139.574 | 5189645.669 | 4437264.712 |

| | | | |
|---|---|---|---|
| 4566.41442 | 16732.33613 | 9115.139189 | 12562.35369 |
| 27526737.41 | 21153096.49 | 25092850.59 | 16781340.1 |
| 5044545.119 | 3796902.647 | 5891403.537 | 3995141.682 |
| 3006441.215 | 2508915.33 | 3707640.878 | 2029225.619 |
| 123827382.5 | 135440095.5 | 130262830 | 133934048.9 |
| 5632.286668 | 7784.955597 | 18312.79744 | 9161.987862 |
| 111263299.4 | 113405026.3 | 119729213.1 | 118049422.7 |
| 115922050.3 | 106408475.8 | 145139273.9 | 105099926.6 |
| 14281.73638 | 7135.445595 | 16768.97212 | 3166.56625 |
| 62243.76951 | 68251.53464 | 67174.47148 | 40496.29046 |
| 10874.67538 | 20930.24 | 25102.70539 | 7991.986635 |
| 31361.77482 | 26729.74497 | 29304.19624 | 10808.15518 |
| 186773.9039 | 140945.117 | 215656.5291 | 291093.3287 |
| 441270.227 | 615328.0586 | 397361.9065 | 458114.48 |
| 50557.87552 | 52869.30449 | 47499.81133 | 29872.47592 |
| 27118438.87 | 26980397.22 | 29652897.1 | 25013983.82 |
| 7499.600674 | 6708.856642 | 3328.551696 | 4132.71541 |
| 3103.67774 | 2142.747888 | 5261.766153 | 3931.591069 |
| 10133.07551 | 17919.96109 | 13641.38553 | 7448.230351 |
| 45957286.58 | 60632944.57 | 60918266.92 | 55902611.15 |
| 88681.34292 | 111270.3699 | 86163.56202 | 93697.70249 |
| 2740511.493 | 3386867.759 | 4716853.067 | 4411861.689 |
| 17344.32373 | 25093.498 | 51887.57531 | 27574.6053 |
| 257667.5051 | 632463.5871 | 687006.9159 | 471010.6872 |
| 36031079.31 | 42838206.76 | 49894662.54 | 39391106.86 |
| 1281267.796 | 1469911.255 | 1093388.807 | 891205.2586 |
| 7772.500386 | 12932.73439 | 10179.80362 | 9565.936095 |
| 1508570.419 | 1851374.404 | 2549278.93 | 2385367.69 |
| 632049.1113 | 773180.3361 | 903999.983 | 764423.2142 |
| 1462076.136 | 1515557.426 | 1896391.306 | 1269940.785 |
| 70151.43649 | 56080.99399 | 58986.68056 | 51982.07798 |
| 105618.3989 | 122939.9283 | 118062.4162 | 87108.83923 |
| 17537090.43 | 21749591.97 | 3566630.305 | 17654811.69 |
| 205491.5552 | 238955.5794 | 192406.547 | 424844.9737 |
| 83856.50224 | 67133.12218 | 81616.08308 | 50203.16286 |
| 841910.7567 | 944229.967 | 633387.0182 | 879795.8883 |
| 345290.5441 | 441035.6644 | 469174.4243 | 320696.643 |
| 16584350.57 | 18177510.95 | 20490755.86 | 19697433.12 |
| 74466026.35 | 84137657.19 | 86524718.56 | 68394721.88 |
| 232290.8555 | 97968.51699 | 125175.1165 | 319519.8629 |
| 464558.6486 | 523465.2675 | 599430.4652 | 500581.0062 |
| 7364.570965 | 12423.97022 | 35408.06233 | 29511.09676 |
| 89389.94686 | 72068.41531 | 46903.91695 | 21544.01816 |
| 2088600.823 | 2458500.259 | 2640742.042 | 2923239.507 |
| 41505182.71 | 46309828.26 | 39339270.83 | 44753131.49 |
| 164146.9866 | 185299.2137 | 129071.592 | 164173.6309 |
| 3345.578245 | 4212.459574 | 2492.765397 | 1689.528335 |
| 192301838.8 | 202148164.8 | 77549846.71 | 208323489 |
| 5020.862397 | 7530.535531 | 9340.925159 | 6694.558271 |
| 1112083.706 | 1065981.045 | 1337143.371 | 849874.1118 |
| 18170.90656 | 12092.35027 | 26247.79605 | 20606.02541 |
| N/A | 9173.460915 | 2980.972805 | 1673.889206 |
| 511004.4524 | 445865.0411 | 406596.9079 | 316725.9664 |
| 13773011.03 | 14129399.73 | 15276132.87 | 13671469.28 |
| 57014018.52 | 61694653.16 | 66893522.16 | 64902536.67 |
| 140222.8233 | 152571.6698 | 140939.3091 | 159779.7516 |
| 100187.4807 | 84769.62162 | 86696.84905 | 137040.1494 |
| 35763.38129 | 22763.47591 | 48965.0593 | 48314.47218 |
| 1161135.425 | 1876899.245 | 2295476.602 | 1772744.662 |
| 22102148.1 | 24787223.17 | 18136770.5 | 23872264.21 |
| 3358830.158 | 6274134.803 | 5153250.732 | 5757683.469 |
| N/A | 4943.450848 | 6226.920514 | 2399.384583 |

| | | | |
|---|---|---|---|
| 13281382.9 | 14481621.83 | 20443263.02 | 17461015.8 |
| 100008149 | 114783069.9 | 125166849.9 | 114163644.4 |
| 208712.0952 | 166135.1274 | 206526.0233 | 218399.8313 |
| 1809064.983 | 2062621.558 | 1876783.848 | 1902248.444 |
| 2510665.721 | 3050474.523 | 1814042.471 | 1486261.881 |
| 55155.07662 | 91999.84175 | 125659.5003 | 61283.25276 |
| 813781.0932 | 1173645.622 | 1127000.696 | 973256.5306 |
| 18131259.56 | 24271996.84 | 18463321.13 | 22810715.25 |
| 134057.9374 | 175198.4928 | 136647.5059 | 122207.9104 |
| 20810.96128 | 6974.194041 | 11380.17086 | 10397.5527 |
| 175232.933 | 177027.4162 | 124161.1094 | 175355.4173 |
| 129062777.2 | 134304953.3 | 126984791.3 | 134727369.4 |
| 33426.10176 | 35187.29758 | 46224.68381 | 36828.31978 |
| 5788335.734 | 7269650.226 | 6369981.757 | 7340547.431 |
| 569446.7002 | 748747.551 | 662176.6078 | 1155655.622 |
| 48337.14425 | 38382.98519 | 145803.1612 | 60769.33494 |
| 605270.1764 | 710001.0236 | 655948.4028 | 674872.9795 |
| 678215.0853 | 837586.0253 | 679110.1407 | 524215.6168 |
| 4852789.984 | 5812320.988 | 6899368.836 | 6163531.023 |
| 42443.81173 | 73330.03208 | 50740.61353 | 28620.65429 |
| 17527.18924 | 23313.25242 | 21011.36924 | 8556.829039 |
| 30115.96116 | 60950.13821 | 64541.39713 | 102535.9998 |
| 351936.7582 | 1361194.941 | 976348.5224 | 1360052.589 |
| 16685.80018 | 25195.42276 | 28004.01448 | 11556.20647 |
| 20269371.27 | 19273805.16 | 18620063.76 | 16436762.12 |
| 12272628.25 | 8980965.095 | 9468923.593 | 8324993.595 |
| 4585237.099 | 6000513.115 | 6123376.933 | 6566543.639 |
| 45210.04439 | 111269.8895 | 81976.74266 | 144689.6895 |
| 1562857.258 | 2517275.821 | 1562565.843 | 3694257.164 |
| 26241219.77 | 40645276.21 | 32993473.32 | 33415513.54 |
| 5767301.212 | 5023779.783 | 4614279.951 | 5414636.204 |
| 58817407.71 | 73838653.87 | 83532914.04 | 75224238.74 |
| 3258025.818 | 3371555.145 | 3507006.793 | 3368480.785 |
| 130988.3864 | 132947.2051 | 212050.0281 | 102134.1988 |
| 175734.2135 | 183049.2526 | 227881.7278 | 196816.0282 |
| 15181777.96 | 21928427.05 | 19584002.52 | 21351818.54 |
| 11691391.31 | 8311806.049 | 10102198.04 | 7427254.199 |
| 46359153.39 | 49825840.41 | 53473307.58 | 54545012.99 |
| 1225741.194 | 1587186.591 | 1632216.88 | 1189214.648 |
| 11710.31515 | 7529.48508 | 12551.98258 | 7529.775084 |
| 1437173.32 | 1104054.768 | 1105664.815 | 1144708.868 |
