## Supplemental Table 1 for "Fatty acid elongases 1-3 have distinct roles in mitochondrial function, growth and lipid homeostasis in *Trypanosoma cruzi*"

| | fold change | | | | | |
|---|---|---|---|---|---|---|
| | *elo1* | *elo2* | *elo3* | *elo1::elo1-gfp* | *elo2::elo2-gfp* | *elo3::elo3-gfp* |
| CE(13:0) | 3.657228164 | 5.306867138 | 2.857843111 | 1.54806918 | 1.959560713 | 1.709550356 |
| CE(22:5) | 1.36791876 | 2.083044205 | 0.99750988 | 0.772470138 | 1.104826835 | 0.812774914 |
| Cer-ADS(d23:0/19:0) | 1.328710253 | 6.220181974 | 0.086624785 | 2.877149036 | 0.961719986 | 0.719677885 |
| Cer-AS(d18:1/16:1) | 3.131639157 | 3.01509695 | 0.153090112 | 2.566602067 | 1.962069368 | 1.780329159 |
| Cer-AS(d23:1/19:0) | 3.977696876 | 1.307000591 | 0.286698551 | 1.067203393 | 0.39389353 | 0.484024118 |
| Cer-AS(d26:1/16:0) | 0.908271673 | 0.08622212 | 0.426205231 | 1.194348763 | 0.486426092 | 0.903888581 |
| Cer-NDS(d16:0/24:0) | 0.772546235 | 1.21196946 | 8.891372943 | 2.716944346 | 0.713632409 | 0.877892674 |
| Cer-NDS(d16:0/24:1) | 1.444300508 | 3.402161206 | 3.333472373 | 1.002615974 | 0.591803556 | 0.637578019 |
| Cer-NDS(d17:0/22:0) | 1.27460724 | 2.594880559 | 4.537276264 | 1.322205198 | 0.996271873 | 0.886979246 |
| Cer-NDS(d18:0/18:1) | 1.483765049 | 12.87564895 | 2.116611361 | 2.476962544 | 1.561750861 | 1.655340139 |
| Cer-NDS(d18:0/20:0) | 1.324154598 | 2.270463861 | 1.994436181 | 0.985206209 | 1.257174945 | 0.786725533 |
| Cer-NDS(d18:0/23:0) | 1.067568142 | 1.593791226 | 5.663697329 | 3.362186262 | 0.726001856 | 0.890736306 |
| Cer-NDS(d18:0/24:1) | 1.90282375 | 2.243131006 | 2.749197885 | 1.175291921 | 0.452657419 | 0.26578635 |
| Cer-NDS(d18:0/25:1) | 1.815899374 | 2.352825284 | 5.399780786 | 3.056823863 | 0.923374276 | 1.09213816 |
| Cer-NDS(d18:0/26:1) | 1.350740424 | 2.263770306 | 6.250956249 | 3.41380853 | 0.987605028 | 1.042641 |
| Cer-NDS(d18:0/28:1) | 1.366246162 | 10.77246836 | 15.6678972 | 2.372279808 | 0.830077693 | 0.525421382 |
| Cer-NS(d16:1/16:0) | 1.939506069 | 1.902858894 | 22.13728635 | 3.032132799 | 1.175214529 | 1.47264571 |
| Cer-NS(d17:1/14:0) | 3.177780436 | 3.261503354 | 51.03367413 | 2.269115915 | 1.273874516 | 1.178192874 |
| Cer-NS(d17:1/16:0) | 2.172191853 | 1.82273913 | 6.318828545 | 2.394142203 | 0.897075762 | 1.164039631 |
| Cer-NS(d17:1/22:0) | 1.697387489 | 2.956411757 | 2.179312471 | 1.040640521 | 1.313660208 | 1.280013899 |
| Cer-NS(d17:1/24:1) | 2.676538876 | 3.610720449 | 3.443910828 | 1.821674951 | 2.180609846 | 1.630661764 |
| Cer-NS(d17:1/40:2) | 1.174149001 | 1.82909504 | 0.989377695 | 0.580989118 | 0.826101395 | 0.820506296 |
| Cer-NS(d18:1/20:0) | 1.890411762 | 2.691805846 | 2.608587495 | 1.175178776 | 1.220658683 | 1.211473847 |
| Cer-NS(d18:1/22:0) | 1.247190479 | 2.121416711 | 1.954894725 | 1.027254563 | 1.091538467 | 0.976403903 |
| Cer-NS(d18:1/25:1) | 1.830649062 | 2.195006364 | 2.223989234 | 1.890579297 | 0.973359997 | 0.930058099 |
| Cer-NS(d18:1/26:0) | 0.24115814 | 0.493409061 | 1.180571161 | 2.727441778 | 0.656646821 | 1.16161708 |
| Cer-NS(d18:1/26:1) | 0.508479434 | 3.016257478 | 1.838969563 | 2.739890945 | 0.822588079 | 1.099725693 |
| Cer-NS(d18:1/32:2) | 0.511935675 | 0.419259923 | 1.199330817 | 2.520411714 | 0.695170952 | 1.036140323 |
| Cer-NS(d18:2/16:1) | 2.096507481 | 5.468374583 | 2.680861369 | 1.894801755 | 1.200283352 | 0.876939171 |
| Cer-NS(d18:2/17:0) | 3.02437943 | 2.925552885 | 1.978340282 | 1.9992731 | 0.805514718 | 1.090472172 |
| Cer-NS(d18:2/18:0) | 2.168862068 | 3.15446809 | 2.243484301 | 1.595276582 | 1.043559962 | 1.076438481 |
| Cer-NS(d18:2/20:0) | 2.256906182 | 2.712316804 | 2.32724577 | 1.350285321 | 1.291992264 | 1.212869589 |
| Cer-NS(d18:2/21:0) | 2.015224573 | 2.99535735 | 2.314664369 | 1.149250616 | 1.306775839 | 1.410013954 |
| Cer-NS(d18:2/22:0) | 2.224553409 | 2.886514653 | 2.203501845 | 1.242405925 | 1.45192225 | 1.445684678 |
| Cer-NS(d18:2/22:1) | 3.024309234 | 5.010870409 | 3.481649181 | 1.609442203 | 2.009934664 | 1.837484749 |
| Cer-NS(d18:2/23:1) | 2.619938035 | 4.188640144 | 2.998214942 | 1.475775401 | 1.743664229 | 1.578279259 |
| Cer-NS(d18:2/24:1) | 2.614296639 | 3.687537468 | 2.788743922 | 1.513290127 | 1.631033376 | 1.475946609 |
| Cer-NS(d18:2/24:2) | 3.056963442 | 4.812524327 | 3.319306194 | 1.674945367 | 1.683832182 | 1.452565532 |
| Cer-NS(d18:2/25:1) | 2.796461799 | 4.40665707 | 2.519567781 | 1.608931458 | 1.528418386 | 1.346901907 |
| Cer-NS(d18:2/26:0) | 0.621043682 | 3.10031086 | 1.524540602 | 2.794906767 | 0.809007056 | 0.911683802 |
| Cer-NS(d18:2/26:1) | 1.871604486 | 1.981493334 | 2.947928078 | 2.317843246 | 0.92393709 | 0.881349752 |
| Cer-NS(d18:2/26:2) | 2.977393566 | 4.685977953 | 2.639666816 | 2.136109524 | 1.493271692 | 1.162892492 |
| Cer-NS(d19:2/16:0) | 4.166632185 | 2.902435885 | 1.84085497 | 4.024037554 | 2.320237967 | 2.68226618 |
| Cer-NS(d19:2/24:2) | 0.821756525 | 1.903977546 | 3.325859176 | 0.724778531 | 0.371057819 | 0.040798455 |
| Cer-NS(d20:1/24:1) | 2.529735003 | 5.677837499 | 5.540395844 | 2.595218757 | 1.018913867 | 0.769000896 |
| DG(14:0_16:0) | 0.784107245 | 1.726014118 | 6.632926495 | 3.79099419 | 1.717573686 | 2.119479193 |
| DG(14:0_18:1) | 0.428845937 | 1.08244101 | 4.8403836 | 1.678747971 | 1.466358567 | 3.014357209 |
| DG(14:0_18:2) | 0.251620945 | 0.489294289 | 12.46037461 | 1.263668183 | 1.040179131 | 2.527235045 |
| DG(16:0_18:1) | 0.2530986 | 0.51642252 | 1.01302077 | 0.963045374 | 1.584658029 | 2.761549724 |
| DG(16:0_18:2) | 0.531099555 | 0.818344508 | 2.046797689 | 0.557776412 | 0.83659415 | 1.849267132 |
| DG(16:0_20:4) | 0.767550597 | 2.377887313 | 1.631150436 | 1.831303467 | 2.086773306 | 2.623074459 |
| DG(16:1_18:2) | 0.357483289 | 0.94287491 | 3.883138371 | 1.09698792 | 1.8901024 | 4.28106977 |
| DG(18:0_18:0) | 1.302139617 | 1.299005619 | 2.228373758 | 1.435226213 | 0.716871671 | 0.639729003 |
| DG(18:0_20:0) | 1.090744741 | 1.305347849 | 2.747998734 | 1.51600285 | 0.700874173 | 0.579131322 |
| DG(18:0_22:6) | 1.350161025 | 3.119530003 | 11.23840281 | 1.146955695 | 2.676789964 | 2.147742214 |
| DG(18:0_24:0) | 0.364876395 | 0.384622084 | 0.451819774 | 0.834511036 | 0.405474025 | 0.73079209 |
| DG(18:1_22:6) | 3.541820518 | 6.536429794 | 22.25740362 | 3.910500975 | 11.08723466 | 17.22049626 |
| DG(18:1_23:0) | 0.199031973 | 0 | 1.93901872 | 1.209837503 | 1.857553654 | 1.978094789 |
| DG(18:2_20:3) | 1.240519624 | 4.01872066 | 2.110096026 | 0.929899562 | 2.549037573 | 3.323703026 |

| | | | | | | |
|---|---|---|---|---|---|---|
| DG(18:2_20:4) | 0.614148057 | 0.926917761 | 2.124202325 | 0.664422528 | 1.720857918 | 2.399190913 |
| DG(18:2_22:0) | 0.700845287 | 1.986026315 | 1.790818347 | 0.566266542 | 1.599097708 | 4.28691266 |
| DG(18:2_22:4) | #DIV/0! | #DIV/0! | #DIV/0! | #DIV/0! | #DIV/0! | #DIV/0! |
| DG(18:2_26:0) | 0.303870076 | 1.185063167 | 1.578058561 | 0.695088044 | 3.874195522 | 7.89074383 |
| DMPE(18:0_18:1) | 0.99456147 | 2.401666627 | 0.395553284 | 1.317880901 | 4.621219779 | 3.913037147 |
| DMPE(18:2_18:2) | 2.183131155 | 12.11858108 | 1.442256918 | 1.067647342 | 3.816641397 | 2.881026563 |
| DMPE(18:2_18:3) | 0.744557403 | 1.43215104 | 0.659973996 | 1.09858121 | 1.477705688 | 1.431204848 |
| FAHFA(18:1/19:0) | 3.810442241 | 0.019305632 | 0.217068121 | 1.192895074 | 1.552407759 | 6.544913075 |
| FAHFA(18:1/25:0) | 3.654961432 | 0.023022003 | 0.419479458 | 0.99227172 | 1.476746691 | 3.870225372 |
| FAHFA(18:1/27:0) | 3.515132901 | 0.04671339 | 0.184639581 | 0.678173159 | 1.479229554 | 2.982677677 |
| FAHFA(18:2/17:0) | 6.276957907 | 0.054208728 | 0.36069874 | 1.928995739 | 0.924348846 | 3.929181484 |
| FAHFA(18:2/17:1) | 12.77330791 | 0.008182846 | 0.405024013 | 0.871183589 | 0.728513772 | 2.544654246 |
| FAHFA(18:2/18:1) | 4.356091263 | 0.140483157 | 0.377062169 | 0.424826764 | 0.502051359 | 1.361818342 |
| FAHFA(18:2/18:1) | 4.356091263 | 0.140483157 | 0.377062169 | 0.424826764 | 0.502051359 | 1.361818342 |
| FAHFA(18:2/25:0) | 3.438035373 | 0.045227047 | 0.085255008 | 1.003111554 | 1.015795185 | 2.618769878 |
| FAHFA(18:2/26:0) | 3.994853605 | 0.287316365 | 0.233578877 | 1.15914401 | 0.846636869 | 1.690179665 |
| FAHFA(18:2/27:0) | 3.947276149 | 0.040297208 | 0.109489549 | 0.838250794 | 1.132643976 | 2.19317057 |
| FAHFA(20:1/27:0) | 2.745348875 | 0.059031949 | 0.217392955 | 0.612974115 | 1.74443026 | 2.995115565 |
| HexCer-NS(d18:1/16:0) | 1.48044054 | 2.856745249 | 1.125592807 | 0.451702064 | 0.619254389 | 0.385329469 |
| HexCer-NS(d18:1/22:0) | 1.146967582 | 1.969513371 | 1.561780926 | 0.814437558 | 0.925249721 | 1.024097353 |
| lanosteryl | 0.59113781 | 0.31387154 | 0.286131975 | 1.044702579 | 0.838549418 | 1.680433144 |
| LPC(12:0) | 1.355474211 | 1.877058322 | 0.821798462 | 2.445537971 | 0.756703517 | 0.550114237 |
| LPC(14:1) | 1.476241089 | 0.329518618 | 10.57217094 | 1.641795943 | 0.358180424 | 0.814147599 |
| LPC(16:0) | 0.664280477 | 0.267757288 | 0.632683826 | 0.934046975 | 0.456630627 | 0.864076527 |
| LPC(17:2) | 0.926568649 | 0.416293606 | 0.582074034 | 1.168920247 | 0.504044898 | 0.716974326 |
| LPC(18:1) | 0.815465674 | 0.684220031 | 0.428095861 | 0.836464454 | 1.02230691 | 0.81929391 |
| LPC(18:2) | 0.686321718 | 0.534527041 | 0.437262472 | 0.845296026 | 0.619618087 | 0.723757471 |
| LPC(18:3) | 0.973935287 | 7.076755318 | 0.868370266 | 0.780251245 | 0.773548957 | 0.676251251 |
| LPC(19:0) | 2.173753342 | 0.791147011 | 0.257053541 | 1.207649328 | 0.613511253 | 0.92405805 |
| LPC(20:0) | 1.734722559 | 1.095954666 | 0.36337162 | 1.222961586 | 0.650077057 | 1.065064418 |
| LPC(20:1) | 1.209222262 | 0.935776837 | 0.285315516 | 0.758240518 | 0.849986706 | 0.957818265 |
| LPC(20:2) | 1.515698811 | 2.148387431 | 0.585950852 | 0.84180689 | 0.823512344 | 0.782035474 |
| LPC(22:2) | 3.662973895 | 3.075924529 | 1.41056109 | 1.505815191 | 1.295598968 | 1.367324115 |
| LPC(O-20:1) | 1.522168252 | 1.475522986 | 1.398099443 | 1.673080102 | 0.767280938 | 1.420428809 |
| LPE(12:0) | 1.033060585 | 10.55377081 | 3.192119216 | 3.67797454 | 1.383937947 | 0.653328596 |
| LPE(16:1) | 0.707490747 | 0.199807032 | 14.21670898 | 1.878799922 | 0.795914825 | 0.841095061 |
| LPE(20:3) | 2.532461973 | 2.837287101 | 3.090780779 | 1.743435221 | 1.259244944 | 1.058519919 |
| LPE(20:4) | 1.831893697 | 3.354878956 | 2.835057722 | 1.060689718 | 1.161630188 | 1.222372225 |
| LPE(22:4) | 4.070146881 | 5.334679952 | 2.230771359 | 1.009552511 | 2.600940423 | 1.314989096 |
| LPE(22:5) | 4.293662323 | 3.616677407 | 3.716146706 | 2.899049149 | 1.467108046 | 1.590893353 |
| LPE(26:0) | 1.832466567 | 5.782424257 | 2.943209356 | 11.3298095 | 0.761867612 | 0.659472629 |
| LPE(P-18:1) | 0.299260742 | 0.201670304 | 0.488516341 | 0.588329493 | 0.810161588 | 0.991336346 |
| LPE(P-20:0) | 1.367708078 | 1.520133847 | 1.61929124 | 2.823614244 | 0.911706097 | 1.82040869 |
| MGDG(14:0_18:1) | 0.102767209 | 2.533759702 | 0 | 0.679792729 | 0.895899682 | 0.741873809 |
| MMPE(18:2_18:2) | 3.271547642 | 12.23141598 | 1.876408438 | 2.235917527 | 1.495201107 | 1.601079883 |
| OxLPC(15:1(COOH)) | 0.820907128 | 0.320117705 | 5.703789563 | 1.442175743 | 0.467719409 | 0.623546314 |
| OxLPC(15:2(CHO)) | 1.002807216 | 0.05626033 | 7.200764832 | 1.12825317 | 0.399337317 | 0.639585384 |
| OxLPC(15:3(CHO)) | 0.97687962 | 0 | 19.12782353 | 1.10878479 | 0.077072536 | 0.14987305 |
| OxLPC(17:2(Ke)) | 0.744801678 | 0.325724669 | 0.207759832 | 0.657713324 | 0.820065727 | 0.916154875 |
| OxLPC(17:2(OO)) | 0.748889815 | 0.380603612 | 0.213417745 | 0.668618466 | 0.816727589 | 0.901596955 |
| OxLPC(17:4(CHO)) | 2.499260313 | 9.538680388 | 0.677936412 | 0.559512114 | 0.496761552 | 0.101625337 |
| OxLPC(19:4(COOH)) | 6.488317259 | 8.055323672 | 0 | 1.367170079 | 1.497818226 | 1.851600223 |
| OxLPC(20:3(OH)) | 1.282385773 | 1.267998321 | 0.714599242 | 1.112069539 | 0.577724782 | 0.901063412 |
| OxLPC(4:0(CHO)) | 4.212471977 | 4.896288914 | 4.99613009 | 0.711858859 | 1.566809592 | 2.398608402 |
| OxPE(18:0_18:1(1O)) | 2.527559146 | 3.051632308 | 0.322891105 | 0.492685327 | 2.332746949 | 1.647050072 |
| OxPE(18:1_18:0(1O)) | 1.678093436 | 1.499745836 | 1.769398153 | 0.826500531 | 1.67563853 | 1.70911719 |
| OxPE(18:1(OH)_18:1(OH)) | 1.848630873 | 1.127260875 | 0.529971029 | 1.623649387 | 1.089591156 | 1.280161915 |
| OxTG(14:0_16:0_18:3(OO)) | 0 | 0 | 8.577660211 | 1.091492315 | 0.362801975 | 2.198292582 |
| OxTG(16:0_16:0_18:3(OO)) | 0.108514604 | 0.198089759 | 1.287649076 | 0.527014593 | 0.510974681 | 1.769108753 |
| OxTG(16:0_16:1_18:2(OH)) | 0.29370619 | 0.746916767 | 1.590055955 | 0.510299513 | 0.846032384 | 2.880029425 |
| OxTG(16:0_18:0_18:2(OH)) | 0.611929367 | 0.369626604 | 0.653178152 | 0.22090031 | 0.470610879 | 1.580347836 |
| OxTG(16:0_18:1_18:1(OH)) | 0.517074837 | 0.41264582 | 0.508759006 | 0.390756164 | 0.704056689 | 2.302675454 |

| | | | | | | |
|---|---|---|---|---|---|---|
| PA(18:2_22:5) | 6.924394123 | 14.92079796 | 3.083115084 | 1.581320476 | 0.586995493 | 1.124259724 |
| PC(10:0_18:2) | 0.91510974 | 111.7155129 | 7.011309842 | 4.176216233 | 3.674303698 | 2.904991482 |
| PC(12:0_18:2) | 1.293776134 | 6.929359724 | 20.23246553 | 2.37357897 | 3.556485358 | 1.580080428 |
| PC(14:0_14:0) | 0.860224439 | 13.57818085 | 91.97219394 | 3.26348285 | 0.998332083 | 1.060838641 |
| PC(14:0_16:1) | 0.982155295 | 5.653674905 | 114.1427354 | 2.005136705 | 3.756825844 | 2.177967158 |
| PC(14:0_18:2) | 0.481672876 | 0.419450151 | 19.78484036 | 1.782759827 | 1.684328049 | 1.467566078 |
| PC(16:0_16:1) | 0.375561248 | 0.318409281 | 11.34949943 | 1.283920689 | 1.379741488 | 1.459437158 |
| PC(16:0_26:0) | 0.399954579 | 1.726319435 | 1.010259601 | 5.877364263 | 0.465638353 | 0.376318261 |
| PC(17:0_18:0) | 1.363060219 | 0.888748185 | 0.032821015 | 3.341135009 | 1.046989777 | 1.114618084 |
| PC(17:0_20:3) | 2.546855072 | 4.963796972 | 1.933645244 | 3.015151477 | 6.263309686 | 3.529154464 |
| PC(17:1_18:1) | 0.687005141 | 1.372086937 | 0.444172142 | 0.909576129 | 1.825254007 | 1.433083269 |
| PC(17:2_22:5) | 2.971883271 | 3.432918343 | 2.269902191 | 3.712957535 | 4.623489787 | 3.697451859 |
| PC(17:2_22:6) | 2.734136414 | 2.768199864 | 2.325214625 | 4.551358583 | 3.941944718 | 3.498697889 |
| PC(18:0_18:0) | 0.693101751 | 0.41604625 | 0.183456535 | 1.666576082 | 0.379822518 | 0.590891999 |
| PC(18:0_20:3) | 1.835293975 | 6.136073527 | 1.225104538 | 1.854162394 | 4.596393902 | 2.606213075 |
| PC(18:0_20:4) | 0.975513922 | 5.58508059 | 2.083346084 | 1.300504888 | 2.974970845 | 2.074347843 |
| PC(18:0_22:0) | 0.17253119 | 0.427691883 | 0.678547409 | 3.94936571 | 0.192991095 | 0.212688979 |
| PC(18:0_22:4) | 3.432951559 | 12.39540461 | 2.423567294 | 1.457571002 | 5.04266793 | 3.871678371 |
| PC(18:0_22:5) | 1.584335603 | 2.700178593 | 1.219740317 | 1.111113432 | 2.413916046 | 2.169748646 |
| PC(18:0_22:6) | 1.644626792 | 3.745931468 | 1.63817179 | 1.484317041 | 3.614996118 | 2.535182828 |
| PC(18:1_22:6) | 2.719119722 | 3.781971792 | 3.027195922 | 4.377629738 | 6.09383211 | 4.24033905 |
| PC(18:2_18:3) | 0.796052528 | 4.9040882 | 2.609096917 | 1.231539796 | 1.575158576 | 1.117488251 |
| PC(18:2_20:1) | 1.979303009 | 8.721963194 | 0.76687516 | 0.641748615 | 3.338233572 | 2.077083627 |
| PC(18:2_22:1) | 2.105107796 | 6.431711103 | 2.026344792 | 1.418522364 | 1.897637784 | 2.360091611 |
| PC(18:2_26:0) | 0.556513064 | 4.534303144 | 2.104353951 | 0.929603444 | 3.773109656 | 4.808842124 |
| PC(18:3_22:6) | 3.222337734 | 13.20075011 | 3.863459322 | 3.174518031 | 3.494930224 | 2.195937567 |
| PC(20:1_22:6) | 4.084364331 | 8.384694179 | 2.231796992 | 3.239344739 | 4.422486407 | 3.379278619 |
| PC(20:3_22:5) | 9.906164533 | 9.114842197 | 4.566832227 | 6.966737902 | 5.125409749 | 5.483184758 |
| PC(20:3_22:6) | 5.731099734 | 4.759225857 | 2.425232554 | 4.113371486 | 3.059686427 | 2.955501932 |
| PC(20:4_22:6) | 3.121986604 | 3.840368717 | 2.366291976 | 2.505664071 | 2.054419431 | 2.028089366 |
| PC(22:5_22:5) | 5.424887498 | 2.979925069 | 2.576239916 | 5.307881028 | 2.183998546 | 2.44970325 |
| PC(22:6_22:6) | 5.988992911 | 3.976728272 | 2.316645842 | 6.383510817 | 2.649960698 | 2.977024996 |
| PC(O-16:0/18:2) | 0.639006668 | 0.569808918 | 4.225644361 | 0.793222616 | 1.028179638 | 0.992227121 |
| PC(O-16:1/22:6) | 1.794171355 | 1.680089413 | 3.176381832 | 1.706091769 | 1.483396486 | 1.556001741 |
| PC(O-18:0/17:2) | 0.966961866 | 1.706566631 | 2.312242406 | 1.834701748 | 2.26378436 | 1.392788463 |
| PC(O-18:1/20:4) | 1.191942261 | 1.54361436 | 1.683171099 | 1.05067504 | 1.37277341 | 0.854798811 |
| PC(O-18:1/22:5) | 1.335145228 | 1.351560236 | 1.992138569 | 1.04640351 | 1.374617237 | 1.324827713 |
| PC(O-20:0/16:0) | 1.143017342 | 1.847384165 | 2.834128528 | 0.94434242 | 0.778761863 | 0.922979025 |
| PC(O-20:0/18:3) | 1.102748952 | 1.512611988 | 2.594452905 | 1.210041143 | 0.858264747 | 0.71649603 |
| PC(P-16:0/16:1) | 2.056194721 | 2.566973091 | 5.80125569 | 1.682181789 | 1.407504096 | 2.5553218 |
| PC(P-16:0/17:2) | 0.661711618 | 0.596982929 | 4.967824024 | 0.820844601 | 1.142549311 | 0.820467845 |
| PC(P-16:0/20:3) | 1.807438077 | 2.494853952 | 2.039800574 | 2.20939082 | 2.787829351 | 1.862567736 |
| PC(P-16:0/22:6) | 1.403882039 | 1.417603227 | 2.808871831 | 1.347451049 | 1.392655497 | 1.211718958 |
| PC(P-16:1/22:4) | 2.382384526 | 0.200253905 | 0.598974755 | 1.740713171 | 0.997773075 | 1.269231944 |
| PC(P-18:0/22:6) | 1.946226577 | 1.817266257 | 3.211057246 | 1.85259932 | 1.473230491 | 1.203756866 |
| PC(P-18:1/18:1) | 0.991302092 | 1.695667022 | 1.010483213 | 0.736694518 | 0.721619184 | 0.646691832 |
| PC(P-18:1/20:1) | 1.232382472 | 1.387469943 | 2.130892638 | 1.302208597 | 0.743580045 | 0.608723816 |
| PC(P-18:1/22:4) | 1.435843165 | 1.685659453 | 0.888822305 | 0.583677983 | 0.609458111 | 0.621117708 |
| PE(12:0_18:2) | 1.120160983 | 11.19557226 | 18.02631306 | 2.48904868 | 3.461302274 | 1.581996358 |
| PE(14:0_17:1) | 0.548160097 | 0.234769639 | 36.18681336 | 2.712200347 | 2.526158213 | 2.225169386 |
| PE(14:0_18:2) | 0.319196999 | 0.463095865 | 27.03515632 | 1.873363991 | 1.812535842 | 1.528347183 |
| PE(14:1_18:2) | 0.742846704 | 0.579007945 | 3.355858239 | 1.092054476 | 0.575792277 | 0.662837272 |
| PE(16:0_18:1) | 0.197447192 | 0.338729346 | 0.771863891 | 1.050114887 | 2.762961294 | 2.124160599 |
| PE(16:0_18:2) | 0.386476382 | 0.644754833 | 1.445506809 | 1.352772643 | 1.997818262 | 1.603664488 |
| PE(16:0_22:6) | 2.769881748 | 7.834469058 | 4.987017461 | 1.078191681 | 1.580188972 | 1.271545813 |
| PE(16:1_16:1) | 0.293925226 | 0.296288065 | 17.69888463 | 1.682143398 | 0.155855893 | 0.507810618 |
| PE(16:1_17:2) | 1.179826712 | 0.611079116 | 11.56127592 | 1.632203699 | 1.151225769 | 0.930177874 |
| PE(16:1_18:3) | 0.511794883 | 1.545972512 | 13.17838155 | 1.139345212 | 0.386808514 | 0.476581599 |
| PE(17:0_18:1) | 0.791114351 | 1.847583541 | 0.479784964 | 0.814362654 | 3.505773805 | 2.807967783 |
| PE(18:0_18:1) | 0.292268382 | 0.829102279 | 0.254238984 | 0.863647906 | 2.947559108 | 2.553508555 |
| PE(18:0_22:6) | 6.545908442 | 30.46829494 | 2.761952792 | 0.815888943 | 7.646345376 | 4.50121364 |
| PE(18:1_18:2) | 0.352286213 | 0.973078802 | 0.555244421 | 0.702461684 | 1.033509058 | 0.978885883 |

| | | | | | | |
|---|---|---|---|---|---|---|
| PE(18:1_19:0) | 0.408553 | 1.664872663 | 0.280047901 | 0.973660699 | 3.720580634 | 3.023849798 |
| PE(18:1_22:4) | 1.132261832 | 3.717710335 | 0.883668818 | 0.940446892 | 7.442617177 | 4.408754587 |
| PE(18:1_24:0) | 0.42502528 | 5.607751508 | 0.691875274 | 1.289867796 | 4.700443272 | 4.545329684 |
| PE(18:1_26:0) | 0.309946569 | 3.342110216 | 0.680865109 | 0.670660685 | 4.841601215 | 5.673892057 |
| PE(18:2_18:3) | 2.488448092 | 1.356159481 | 1.130841229 | 3.254078048 | 1.526191619 | 1.572710756 |
| PE(18:2_20:3) | 2.231473298 | 2.292910827 | 1.515971982 | 2.764314218 | 1.923848337 | 1.922007636 |
| PE(18:2_22:4) | 2.482216222 | 2.922281299 | 1.998317686 | 1.273628638 | 2.691808924 | 2.277655794 |
| PE(18:2_22:5) | 2.442536567 | 3.351181174 | 1.676720158 | 3.644205903 | 1.957684932 | 2.325332931 |
| PE(18:2_22:5) | 2.442536567 | 3.351181174 | 1.676720158 | 3.644205903 | 1.957684932 | 2.325332931 |
| PE(18:2_22:6) | 2.843477079 | 2.514991438 | 1.07878731 | 3.581581076 | 1.584846486 | 1.811173953 |
| PE(18:2_26:0) | 0.704539184 | 11.51400262 | 1.043507191 | 1.30002854 | 5.507322355 | 5.152748787 |
| PE(30:2) | 1.244824732 | 13.43325278 | 26.04054672 | 3.359539305 | 4.590110758 | 1.731233088 |
| PE(32:2) | 4.666809931 | 34.38766421 | 7.914007301 | 4.585197171 | 8.685321841 | 3.27646361 |
| PE(32:5) | 0.306460327 | 14.17976172 | 24.21048329 | 7.052727858 | 14.00112741 | 3.561724295 |
| PE(34:0) | 2.418282493 | 3.27476159 | 6.635254843 | 4.658012221 | 2.200872794 | 2.588268726 |
| PE(39:8) | 3.878913164 | 6.532717475 | 2.784050536 | 5.501712174 | 3.456904639 | 2.178031568 |
| PE(42:3) | 1.230830666 | 7.602418239 | 1.721806353 | 1.383749486 | 2.333625126 | 1.654961067 |
| PE(42:9) | 2.446520412 | 1.605922716 | 0.439422593 | 1.039342731 | 0.807062323 | 0.636589833 |
| PE(O-16:0/12:0) | 3.430569874 | 77.72607855 | 8.52847452 | 7.11769682 | 3.41315558 | 1.012900191 |
| PE(O-16:0/14:1) | 11.29798177 | 68.98118194 | 5854.00856 | 12.05487843 | 5.425233685 | 5.531857482 |
| PE(O-16:0/16:1) | 1.678124781 | 0.710744322 | 62.46535107 | 1.613244888 | 1.229425867 | 1.364897513 |
| PE(O-16:0/17:1) | 3.363263167 | 1.574174243 | 8.438476741 | 2.228731804 | 1.968352286 | 2.513713577 |
| PE(O-16:0/18:1) | 1.443034486 | 1.200753808 | 2.628167653 | 1.1938825 | 1.651993743 | 1.905082186 |
| PE(O-16:0/22:5) | 1.114727126 | 3.039670533 | 0.279955613 | 0.641165103 | 0.663286162 | 0.862655762 |
| PE(O-18:0/17:1) | 3.370368837 | 3.914396148 | 0.791911497 | 1.165919757 | 2.850298603 | 2.829187862 |
| PE(O-18:0/17:2) | 7.475909962 | 5.100317852 | 1.766734533 | 2.02036097 | 2.018131056 | 1.71891046 |
| PE(O-18:0/17:2) | 7.475909962 | 5.100317852 | 1.766734533 | 2.02036097 | 2.018131056 | 1.71891046 |
| PE(O-18:0/18:1) | 1.799530574 | 4.705225403 | 0.658296701 | 0.769577245 | 2.401128344 | 2.206605575 |
| PE(O-18:0/18:2) | 1.690887392 | 1.764753159 | 0.634713968 | 0.881896213 | 0.752281358 | 0.752501108 |
| PE(O-28:0) | 2.768723955 | 33.68004416 | 4.180708905 | 7.623237029 | 1.974227032 | 1.25130755 |
| PE(O-33:1) | 3.340016028 | 1.697232482 | 8.217951792 | 2.076554332 | 2.293502289 | 2.613324814 |
| PE(O-35:1) | 2.9168045 | 3.113664506 | 2.659637741 | 2.088130292 | 3.478135693 | 3.162651444 |
| PE(O-38:2) | 1.551499182 | 1.725749138 | 1.475458799 | 1.687147886 | 1.650155962 | 1.66789116 |
| PE(P-16:0/12:0) | 2.01743776 | 6.595736989 | 7.763466822 | 4.736816551 | 1.23981529 | 0.8191892 |
| PE(P-16:0/14:0) | 3.397086058 | 3.554414177 | 133.6133986 | 4.371009736 | 1.567656115 | 1.612744023 |
| PE(P-16:0/14:1) | 4.03681254 | 14.0488713 | 334.139429 | 4.615221786 | 4.288743848 | 1.525000433 |
| PE(P-16:0/16:0) | 1.161253128 | 0.39159632 | 3.803707922 | 1.340038381 | 0.682542156 | 1.361796847 |
| PE(P-16:0/16:1) | 0.822469015 | 0.282188318 | 13.71371929 | 1.565354916 | 0.816112889 | 1.001224264 |
| PE(P-16:0/17:1) | 1.204802192 | 0.4963694 | 1.95304219 | 2.22813861 | 1.530361234 | 1.855371334 |
| PE(P-16:0/18:1) | 0.725732821 | 0.520691187 | 0.78629452 | 1.219009183 | 1.383115044 | 1.412278205 |
| PE(P-16:0/18:2) | 0.651013646 | 0.487597233 | 1.399910767 | 1.516455744 | 1.093632109 | 1.226010633 |
| PE(P-16:0/22:3) | 1.913734975 | 5.056475669 | 0.903883908 | 1.239162615 | 4.092903312 | 2.170569756 |
| PE(P-16:0/22:4) | 1.504405193 | 2.237956793 | 1.458106975 | 1.219296211 | 0.754978977 | 1.425347403 |
| PE(P-16:1/18:2) | 1.832431768 | 4.188697093 | 6.198385958 | 1.367740872 | 1.502711049 | 1.147578757 |
| PE(P-18:0/17:2) | 1.53601379 | 1.608674518 | 0.460680209 | 1.747392471 | 1.972521111 | 1.837219256 |
| PE(P-18:0/18:2) | 1.163307005 | 1.756016859 | 0.460724135 | 1.151523028 | 1.306488324 | 1.095730353 |
| PE(P-18:0/20:4) | 1.099815777 | 2.099101429 | 1.677925126 | 0.828424175 | 0.621171277 | 0.659000379 |
| PE(P-18:0/22:3) | 4.354405775 | 3.763105003 | 0.718365545 | 1.856147029 | 1.917638272 | 1.334781839 |
| PE(P-18:0/22:4) | 1.498971658 | 2.036861946 | 1.20003148 | 1.333916856 | 1.200279823 | 1.194826915 |
| PE(P-18:0/22:6) | 1.678645922 | 1.80017291 | 1.564213836 | 1.250329527 | 0.952216165 | 1.327666801 |
| PE(P-18:1/17:1) | 1.012534106 | 0.953456323 | 0.333221393 | 1.058352506 | 0.973310635 | 0.978719738 |
| PE(P-18:1/22:6) | 2.201529629 | 2.159384788 | 1.771320333 | 0.93294109 | 1.530478694 | 1.903945528 |
| PE(P-20:0/18:1) | 2.208424669 | 1.303642204 | 0.275829349 | 1.453195195 | 2.05125139 | 2.00385512 |
| PE(P-20:0/18:2) | 2.276309649 | 1.42746345 | 0.444915942 | 1.634925297 | 1.185747432 | 1.238331841 |
| PE(P-20:1/18:1) | 1.34377576 | 1.593087905 | 0.430077499 | 0.971727854 | 1.421703434 | 0.712119392 |
| PEtOH(18:1_18:1) | 4.355616756 | 3.547956661 | 5.007856328 | 7.394189784 | 11.55507286 | 10.35865825 |
| PG(16:0_18:2) | 0.26661597 | 0.467140373 | 1.402995055 | 1.551174032 | 0.920347223 | 1.38391453 |
| PG(18:1_18:2) | 0.410225492 | 0.266842492 | 0.340354844 | 0.950261222 | 1.566257298 | 1.780221134 |
| PI(16:0_17:1) | 2.230199099 | 2.888570551 | 4.188032899 | 2.320015869 | 3.840876513 | 5.189694135 |
| PI(16:1_18:2) | 0.751188493 | 6.432658017 | 7.25346812 | 0.57598345 | 2.721015331 | 6.216445387 |
| PI(17:0_18:1) | 1.1379248 | 2.089676408 | 4.999891804 | 0.893760338 | 2.430337805 | 2.029058962 |
| PI(18:0_18:1) | 0.480398322 | 2.179341315 | 0.373049953 | 0.370126702 | 4.425519957 | 4.377060347 |

| | | | | | | |
|---|---|---|---|---|---|---|
| PI(18:0_18:2) | 0.719614981 | 3.252437092 | 0.992786864 | 0.440305596 | 3.339371375 | 3.506947249 |
| PI(18:0_20:3) | 1.452250619 | 58.04301844 | 4.646359308 | 0 | 12.60715049 | 13.95889153 |
| PI(18:2_18:2) | 1.492753907 | 25.29035197 | 3.602291008 | 0.363260073 | 12.10373099 | 20.37121364 |
| PI(18:2_22:4) | 2.490727574 | 11.85750498 | 2.394440703 | 0.18183169 | 5.291631403 | 10.20668346 |
| PS(15:0_18:1) | 0.100623513 | 0.135990823 | 7.847921267 | 1.025058884 | 4.870327324 | 1.817812948 |
| PS(15:0_18:2) | 0.444934002 | 0.414935875 | 28.9430047 | 2.330744485 | 3.53334903 | 1.336988666 |
| PS(16:0_18:1) | 0.225327622 | 0.241388336 | 0.864110528 | 0.52465453 | 1.819459944 | 0.823241531 |
| PS(16:0_18:2) | 0.287882465 | 0.516813268 | 1.245431581 | 0.718806249 | 1.555452268 | 0.825569777 |
| PS(18:0_18:1) | 0.437490957 | 1.239722959 | 0.343586403 | 0.686600187 | 3.751504715 | 1.49239114 |
| PS(18:1_22:5) | 0.515298893 | 1.363788913 | 0.50958705 | 1.40836821 | 2.91792352 | 2.44356025 |
| PS(18:2_18:2) | 0.486147022 | 0.385523241 | 0.604975184 | 0.731148004 | 1.225633004 | 0.567062488 |
| PS(18:2_19:0) | 1.09147847 | 2.886760045 | 0.405151715 | 0.760387543 | 5.967391948 | 2.345828239 |
| PS(18:2_22:5) | 1.752998099 | 3.025711611 | 1.663390535 | 2.29769378 | 1.739418143 | 1.803847627 |
| PS(22:5_22:6) | 7.545883675 | 17.27244019 | 2.8488151 | 3.25202154 | 2.582452302 | 2.542990323 |
| PS(34:2) | 0.681073438 | 0.38069648 | 0.832412969 | 1.104150676 | 2.3677071 | 1.416877791 |
| PS(34:3) | 0.562904689 | 0.319155043 | 1.772415995 | 2.444602834 | 2.186013972 | 1.793559525 |
| PS(35:1) | 2.064345656 | 2.16890286 | 1.820477944 | 0.824966863 | 5.402551931 | 2.074892537 |
| PS(35:2) | 2.443309768 | 1.45576891 | 2.558075345 | 1.038125347 | 3.720000789 | 1.078679241 |
| PS(36:5) | 1.244646627 | 0.492464925 | 3.368779888 | 2.849696777 | 1.146320653 | 1.595741968 |
| PS(O-16:0/18:2) | 1.110003762 | 0.60197499 | 3.143560321 | 2.761743494 | 1.480874551 | 1.526307069 |
| PS(P-16:0/17:1) | 1.114943146 | 0.310340813 | 11.52946386 | 1.220528581 | 2.340071312 | 0.812861608 |
| PS(P-16:0/18:1) | 0.887409785 | 0.675337338 | 3.07150759 | 1.921527275 | 1.28738529 | 1.502117444 |
| PS(P-16:1/18:3) | 3.732183567 | 1.895403536 | 116.1790935 | 4.390511484 | 8.106514266 | 1.424783419 |
| PS(P-16:1/20:4) | 4.968703604 | 2.851228304 | 2.780633231 | 1.770934169 | 7.953580432 | 1.656302873 |
| PS(P-16:1/22:4) | 3.744562791 | 16.88162991 | 1.291028386 | 0.620989862 | 6.134948405 | 1.117145421 |
| PS(P-18:0/16:1) | 1.691041087 | 0.238905592 | 0.397854983 | 0.181656567 | 0.961184869 | 0.425337521 |
| PS(P-18:0/18:0) | 1.906970034 | 2.265182823 | 1.159192638 | 0.878300004 | 3.823789758 | 1.768038449 |
| PS(P-18:1/18:1) | 1.799893637 | 1.038051966 | 6.950581161 | 1.429461503 | 5.837040323 | 1.934385876 |
| PS(P-20:0/18:2) | 2.388883307 | 3.08333287 | 0.497021266 | 1.247203835 | 3.129078755 | 1.621965102 |
| PS(P-20:1/17:1) | 1.546803731 | 2.856274338 | 0.430908664 | 0.738270893 | 4.416161632 | 1.282485627 |
| PS(P-20:1/18:1) | 2.00696466 | 5.286686461 | 1.058524975 | 0.86666805 | 7.855341883 | 2.223162361 |
| PS(P-20:1/18:2) | 1.187534957 | 1.614319746 | 0.308558414 | 0.940423687 | 4.704174516 | 1.126719489 |
| SM(d18:1/22:0) | 1.244227203 | 1.489098542 | 1.658942066 | 0.84896907 | 0.671485534 | 1.067724729 |
| SM(d18:1/22:1) | 1.426535792 | 1.984036375 | 1.008323789 | 0.777767945 | 0.40393799 | 0.790029972 |
| SM(d18:1/23:0) | 1.243275376 | 2.52546663 | 1.489938284 | 0.558414774 | 0.471380334 | 1.072161403 |
| SM(d18:1/24:1) | 1.19162358 | 1.499882063 | 1.085048025 | 0.313142172 | 0.664181098 | 1.000123042 |
| SM(d18:2/24:0) | 1.04486331 | 2.475266863 | 1.81662457 | 1.663100186 | 1.550288425 | 1.691694677 |
| SM(d20:0/16:1) | 1.552615236 | 2.116607347 | 1.778653888 | 2.09801721 | 1.196290822 | 1.915636925 |
| So(d18:0) | 0.144963124 | 0.178642463 | 0.794201942 | 0.655876304 | 0.93162607 | 0.816878554 |
| TG(10:0_14:0_18:1) | 0.280876454 | 0.375339161 | 16.37602528 | 0.426725814 | 0.51000415 | 1.902820114 |
| TG(10:0_16:1_18:1) | 0.207648534 | 1.275149206 | 19.64650851 | 0.663912052 | 1.290886274 | 3.899168665 |
| TG(12:0_14:0_16:0) | 0.338330764 | 0.741249589 | 26.32010742 | 1.980399593 | 0.684568387 | 0.969751292 |
| TG(13:0_13:0_15:0) | 1.619358007 | 3.713586041 | 3.33084561 | 1.649469876 | 1.537819324 | 1.695350759 |
| TG(14:0_14:0_16:1) | 0.391844604 | 1.77265131 | 37.16001509 | 0.824640874 | 1.04527069 | 2.165114804 |
| TG(14:0_14:0_18:2) | 0.386530356 | 2.086856359 | 24.40437152 | 0.684388385 | 1.224535323 | 2.251826182 |
| TG(14:0_16:0_16:0) | 0.395267445 | 2.179748502 | 4.538569147 | 2.575511363 | 0.931649353 | 1.170668111 |
| TG(14:0_16:0_18:1) | 0.492506409 | 2.509578794 | 3.52507849 | 1.034311914 | 1.03877003 | 1.708590127 |
| TG(14:0_16:0_18:2) | 0.430606526 | 1.968073581 | 5.522772513 | 0.804706554 | 1.09173944 | 1.989099787 |
| TG(14:0_18:2_18:2) | 0.246774371 | 1.007042206 | 4.297702755 | 0.389079432 | 0.690028661 | 1.508736688 |
| TG(14:0_18:2_22:6) | 0.651518463 | 3.172336107 | 3.356085545 | 0.650991903 | 1.168068116 | 1.44069852 |
| TG(14:1_16:0_18:1) | 0.630909501 | 3.103888922 | 2.317887135 | 0.910937051 | 1.581885051 | 2.6699257 |
| TG(14:1_16:1_18:1) | 0.64609704 | 2.540864469 | 3.292654468 | 0.749508732 | 1.317572406 | 2.100089374 |
| TG(14:1_18:2_18:2) | 0.688032152 | 1.991309322 | 3.237579376 | 0.423290809 | 0.95234294 | 1.242211847 |
| TG(15:0_16:0_16:0) | 0.541796613 | 3.021170523 | 3.64310418 | 2.690024754 | 1.415322503 | 1.463185796 |
| TG(15:0_16:0_18:1) | 0.586211293 | 2.961746209 | 2.876537609 | 1.078439292 | 1.248292241 | 1.925674897 |
| TG(15:0_18:1_18:1) | 0.477205408 | 2.552079712 | 1.829153444 | 0.632632672 | 1.363650869 | 2.882210196 |
| TG(15:0_18:2_22:5) | 0.792125308 | 2.767403894 | 1.70912339 | 0.420519328 | 0.988147168 | 1.442328125 |
| TG(15:1_16:1_18:2) | 0.995399104 | 1.002938805 | 2.353291613 | 0.43910355 | 0.726697451 | 1.176335379 |
| TG(16:0_16:0_22:6) | 0.839926354 | 3.690115494 | 2.119934948 | 1.2286166 | 2.741151511 | 3.315144214 |
| TG(16:0_16:1_18:2) | 0.353091603 | 1.302718526 | 2.566038915 | 0.439829266 | 0.87013496 | 1.856399978 |
| TG(16:0_16:1_22:6) | 0.673184009 | 2.862594993 | 2.830118912 | 0.473178501 | 1.143016952 | 1.037388324 |
| TG(16:0_17:0_18:1) | 0.546555347 | 3.022034559 | 1.713942394 | 0.797186267 | 1.099135392 | 2.04919034 |

| | | | | | | |
|---|---|---|---|---|---|---|
| TG(16:0_17:2_18:2) | 0.799490341 | 2.610895108 | 1.622365203 | 0.452454539 | 1.870454666 | 4.366039406 |
| TG(16:0_18:0_18:1) | 0.317603652 | 1.295556393 | 1.067760481 | 0.591040452 | 0.933687293 | 2.166653841 |
| TG(16:0_18:0_20:0) | 0.297269116 | 0.934137412 | 2.638244727 | 1.1786544 | 0.345666964 | 0.412031687 |
| TG(16:0_18:1_18:1) | 0.349900109 | 1.63220541 | 0.920745493 | 0.411413307 | 1.089765292 | 2.581261872 |
| TG(16:1_18:1_18:1) | 0.596387889 | 0.463554836 | 1.868227003 | 0.863393019 | 0.675566977 | 1.027692818 |
| TG(16:1_18:1_8:0) | 0.058646115 | 0.233598868 | 9.69818727 | 0.85327838 | 1.641231647 | 11.28635829 |
| TG(16:1_18:2_18:2) | 0.355196887 | 2.383936817 | 1.934977985 | 0.440613597 | 0.982741805 | 2.459661427 |
| TG(16:1_18:2_18:3) | 0.611626296 | 1.769033165 | 2.543788614 | 0.32758752 | 0.559653811 | 1.023811889 |
| TG(16:1_18:2_24:1) | 0.544991364 | 2.226016852 | 1.973885297 | 0.389477152 | 1.708044102 | 3.240541015 |
| TG(17:0_18:0_18:1) | 0.702448098 | 2.503273644 | 2.041946966 | 0.826357172 | 1.267703181 | 2.556423717 |
| TG(17:0_18:1_18:1) | 0.62321104 | 3.057951007 | 1.474127613 | 0.545954271 | 1.470718026 | 3.013854607 |
| TG(17:1_18:1_18:1) | 0.402393891 | 2.337371742 | 1.055160078 | 0.37212278 | 1.765865585 | 3.87241443 |
| TG(17:1_18:1_20:4) | 0.5436509 | 3.644403884 | 1.969356865 | 0.601389718 | 1.371247087 | 1.557245419 |
| TG(17:1_18:2_22:6) | 1.264342448 | 2.25282604 | 1.212933263 | 0.864857426 | 1.50306803 | 2.416281548 |
| TG(17:2_18:1_18:2) | 0.948906364 | 4.249101588 | 2.388301356 | 1.029664854 | 2.43365283 | 4.556182329 |
| TG(18:0_18:1_18:2) | 0.514575813 | 1.287359678 | 0.713477479 | 0.350178569 | 1.712908029 | 3.134254943 |
| TG(18:0_18:1_22:5) | 1.31541057 | 3.42717375 | 1.637631026 | 0.304036455 | 1.844499418 | 1.319572796 |
| TG(18:1_18:1_18:1) | 1.086146877 | 0.855819679 | 2.54764496 | 1.374654383 | 0.593756471 | 0.835010221 |
| TG(18:1_18:1_20:4) | 0.825397727 | 3.454657066 | 1.897263429 | 1.436940692 | 4.274351365 | 6.361392766 |
| TG(18:1_18:1_22:5) | 1.607335993 | 2.034456228 | 1.866822354 | 0.97216817 | 2.414315306 | 2.834862969 |
| TG(18:1_18:2_18:2) | 0.464261151 | 0.4328885 | 0.367622957 | 0.492298076 | 1.573095891 | 2.821902281 |
| TG(18:1_18:2_18:3) | 0.499650502 | 0.146670421 | 0.363720712 | 0.51070831 | 1.115858787 | 1.986438382 |
| TG(18:1_18:2_19:0) | 1.021377136 | 3.790625349 | 1.84447442 | 0.778286041 | 1.713657222 | 2.301664221 |
| TG(18:1_18:2_20:1) | 0.928667367 | 4.045729149 | 1.396938242 | 0.528011638 | 2.272511238 | 2.953805995 |
| TG(18:1_18:2_20:2) | 0.754933777 | 4.376033173 | 1.789145614 | 0.509498837 | 2.075392494 | 2.347269295 |
| TG(18:1_18:2_20:4) | 0.71436408 | 3.00777796 | 2.226634932 | 0.566510473 | 1.505545098 | 2.122914972 |
| TG(18:1_18:2_21:0) | 0.487312269 | 2.158540121 | 2.38248956 | 0.600540288 | 1.827519909 | 4.468006972 |
| TG(18:1_20:4_22:6) | 1.534667445 | 3.266856254 | 1.704693924 | 0.617210549 | 1.354868071 | 1.826613817 |
| TG(O-16:0_10:0_18:2) | 0.353845979 | 10.17894948 | 7.453610279 | 4.725066117 | 0.397579029 | 1.173065469 |
| TG(O-16:0_14:0_18:1) | 0.457654772 | 0.437113288 | 23.30439904 | 1.575931044 | 0.458867178 | 1.299095808 |
| TG(O-16:0_14:1_18:1) | 0.176169688 | 0.44362926 | 11.64811292 | 1.933515991 | 2.160327892 | 4.807083665 |
| TG(O-16:0_16:0_18:1) | 0.505111082 | 0.258637308 | 2.621197966 | 1.49126493 | 0.531685882 | 1.234467335 |
| TG(O-16:0_16:0_24:0) | 0.38441453 | 1.940570399 | 4.246251339 | 0.898893809 | 0.540504476 | 0.659220006 |
| TG(O-16:0_16:1_18:2) | 1.168120629 | 0.315054329 | 13.84629106 | 2.646255391 | 0.313846957 | 0.51652002 |
| TG(O-16:0_18:0_18:2) | 0.28737391 | 0.140431741 | 0.478970726 | 2.12998027 | 1.19595821 | 2.476585636 |
| TG(O-16:1_18:1_20:2) | 1.149830499 | 0.489269619 | 2.303510213 | 2.354809191 | 0.589358094 | 0.513608022 |
| TG(O-18:0_18:2_22:6) | 2.437650247 | 3.11970761 | 7.310604158 | 1.726169853 | 1.013740479 | 1.330163457 |
| TG(O-18:1_10:0_18:2) | 3.056010161 | 36.93164678 | 61.69400517 | 2.444979696 | 0.275525676 | 0.642975917 |
| TG(O-18:1_18:2_22:6) | 5.51923395 | 15.15050273 | 29.30086445 | 4.854922434 | 0.749789426 | 1.315570286 |
| TG(O-18:1_18:2_24:0) | 0.952356183 | 2.582632616 | 4.388080665 | 1.905096757 | 0.875303245 | 1.213296694 |
| TG(O-20:0_16:0_20:4) | 1.384950699 | 1.295246915 | 4.360996141 | 2.245983856 | 0.594284426 | 0.769766293 |
| TG(O-20:0_17:1_18:1) | 1.188242415 | 1.853535252 | 1.631279259 | 0.90485516 | 1.033081246 | 1.429160659 |
| TG(O-20:0_18:1_22:4) | 0.707042178 | 1.648964299 | 1.305981576 | 0.549113295 | 1.021801857 | 1.21637792 |

| q value | | | | | |
|---|---|---|---|---|---|
| *elo1* | *elo2* | *elo3* | *elo1::elo1-gfp* | *elo2::elo2-gfp* | *elo3::elo3-gfp* |
| 0.218115 | 0.011614 | 0.084636 | 0.076709 | 0.018082 | 0.029948 |
| 0.479642 | 0.006442 | 0.910616 | 0.309445 | 0.705277 | 0.263996 |
| 0.018824 | 0.008294 | 0.021295 | 0.397899 | 0.777344 | 0.041201 |
| 0.025653 | 0.024041 | 0.844626 | 0.808226 | 0.261521 | 0.040699 |
| 0.441716 | 0.025958 | 0.103173 | 0.063919 | 0.04311 | 0.019762 |
| 0.895669 | 0.055424 | 0.06191 | 0.025967 | 0.022796 | 0.111385 |
| 0.652437 | 0.359192 | 0.048033 | 0.342852 | 0.449725 | 0.349247 |
| 0.092969 | 0.049261 | 0.09043 | 0.492086 | 0.087768 | 0.316927 |
| 0.59019 | 0.055424 | 0.01117 | 0.581424 | 0.037887 | 0.100218 |
| 0.058082 | 0.388258 | 0.197183 | 0.193035 | 0.188037 | 0.042779 |
| 0.085916 | 0.015286 | 0.630034 | 0.545064 | 0.092118 | 0.154324 |
| 0.031039 | 0.493176 | 0.732658 | 0.067733 | 0.128821 | 0.029948 |
| 0.341902 | 0.781765 | 0.025789 | 0.283867 | 0.124103 | 0.012009 |
| 0.071054 | 0.829047 | 0.134252 | 0.283281 | 0.196752 | 0.03224 |
| 0.13825 | 0.285764 | 0.007333 | 0.17782 | 0.518458 | 0.03224 |
| 0.125692 | 0.000892 | 0.051182 | 0.00258 | 0.213208 | 0.101463 |
| 0.137309 | 0.015496 | 0.025239 | 0.301093 | 0.033133 | 0.020493 |
| 0.313508 | 0.011121 | 0.029798 | 0.028939 | 0.882568 | 0.757145 |
| 0.347166 | 0.53245 | 0.142272 | 0.072091 | 0.89962 | 0.626414 |
| 0.503953 | 0.005013 | 0.287467 | 0.056983 | 0.250101 | 0.222908 |
| 0.44991 | 0.0063 | 0.053929 | 0.853513 | 0.108789 | 0.126362 |
| 0.627296 | 0.040405 | 0.611451 | 0.095227 | 0.862867 | 0.360909 |
| 0.050726 | 0.015915 | 0.061403 | 0.369311 | 0.207086 | 0.159563 |
| 0.395717 | 0.011121 | 0.098988 | 0.750398 | 0.543174 | 0.725478 |
| 0.030523 | 0.024633 | 0.39983 | 0.029229 | 0.124103 | 0.425657 |
| 0.062209 | 0.003133 | 0.021652 | 0.048212 | 0.011481 | 0.063564 |
| 0.356876 | 0.117748 | 0.602742 | 0.099271 | 0.518458 | 0.463089 |
| 0.167677 | 0.007701 | 0.103719 | 0.300891 | 0.287044 | 0.376413 |
| 0.291473 | 0.047245 | 0.140638 | 0.076644 | 0.657211 | 0.652502 |
| 0.503374 | 0.061939 | 0.072333 | 0.045049 | 0.124103 | 0.108 |
| 0.138796 | 0.003257 | 0.090723 | 0.020431 | 0.06632 | 0.425946 |
| 0.084565 | 0.00661 | 0.033714 | 0.14873 | 0.101329 | 0.095014 |
| 0.785877 | 0.093182 | 0.391984 | 0.066169 | 0.056283 | 0.167283 |
| 0.123067 | 0.020499 | 0.036743 | 0.055164 | 0.843348 | 0.66026 |
| 0.074536 | 0.019967 | 0.030122 | 0.020431 | 0.632673 | 0.370131 |
| 0.599274 | 0.603352 | 0.389178 | 0.352933 | 0.882568 | 0.739663 |
| 0.306559 | 0.62223 | 0.369926 | 0.12232 | 0.043621 | 0.641568 |
| 0.202342 | 0.013812 | 0.090723 | 0.553888 | 0.35549 | 0.228626 |
| 0.86579 | 0.065093 | 0.252452 | 0.075374 | 0.147621 | 0.179637 |
| 0.310044 | 0.041392 | 0.871457 | 0.04003 | 0.20779 | 0.192052 |
| 0.139382 | 0.53245 | 0.113045 | 0.060815 | 0.19415 | 0.321596 |
| 0.082367 | 0.01756 | 0.675029 | 0.038612 | 0.082814 | 0.769347 |
| 0.875892 | 0.092638 | 0.085865 | 0.050503 | 0.120592 | 0.165444 |
| 0.570668 | 0.127159 | 0.26683 | 0.139388 | 0.338434 | 0.434586 |
| 0.097377 | 0.00858 | 0.415328 | 0.058513 | 0.092213 | 0.036886 |
| 0.726999 | 0.443135 | 0.234993 | 0.782926 | 0.520882 | 0.372505 |
| 0.219091 | 0.004845 | 0.073775 | 0.071382 | 0.074826 | 0.71541 |
| 0.2365 | 0.014327 | 0.015933 | 0.020431 | 0.805367 | 0.220644 |
| 0.180956 | 0.015286 | 0.35488 | 0.136845 | 0.761549 | 0.833073 |
| 0.618995 | 0.004235 | 0.019813 | 0.255709 | 0.032077 | 0.538957 |
| 0.662149 | 0.115827 | 0.333773 | 0.317475 | 0.518458 | 0.532885 |
| 0.341902 | 0.108457 | 0.63156 | 0.581742 | 0.499831 | 0.848417 |
| 0.641363 | 0.025958 | 0.430684 | 0.409629 | 0.862618 | 0.318164 |
| 0.149838 | 0.020499 | 0.067113 | 0.762627 | 0.400882 | 0.139493 |
| 0.502564 | 0.038886 | 0.051134 | 0.174805 | 0.89962 | 0.288235 |
| 0.268125 | 0.035356 | 0.026389 | 0.045049 | 0.746383 | 0.697066 |
| 0.156413 | 0.104316 | 0.111626 | 0.254523 | 0.804668 | 0.510249 |
| 0.542518 | 0.165607 | 0.822611 | 0.144173 | 0.511212 | 0.767063 |
| 0.517771 | 0.214923 | 0.265662 | 0.224866 | 0.334544 | 0.096425 |

| | | | | | |
|---|---|---|---|---|---|
| 0.370182 | 0.52796 | 0.365337 | 0.043683 | 0.854893 | 0.334192 |
| 0.276108 | 0.022873 | 0.247196 | 0.644958 | 0.841728 | 0.870765 |
| 0.076468 | 0.01026 | 0.060541 | 0.442812 | 0.272776 | 0.280843 |
| 0.918638 | 0.002219 | 0.858729 | 0.051499 | 0.862618 | 0.178557 |
| 0.05396 | 0.002695 | 0.018891 | 0.028939 | 0.864193 | 0.436373 |
| 0.738529 | 0.210281 | 0.22954 | 0.185546 | 0.301289 | 0.686319 |
| 0.627814 | 0.753139 | 0.408154 | 0.062431 | 0.087156 | 0.109879 |
| 0.761999 | 0.026853 | 0.074295 | 0.364102 | 0.06632 | 0.186046 |
| 0.07963 | 0.004845 | 0.000464 | 0.020431 | 0.43799 | 0.446403 |
| 0.886848 | 0.003688 | 0.456206 | 0.266806 | 0.24405 | 0.151906 |
| 0.087027 | 0.260569 | 0.01843 | 0.28844 | 0.083098 | 0.583164 |
| 0.074536 | 0.534123 | 0.015933 | 0.624754 | 0.071038 | 0.566911 |
| 0.147261 | 0.009735 | 0.061326 | 0.317455 | 0.308558 | 0.240302 |
| 0.147261 | 0.009735 | 0.061326 | 0.317455 | 0.308558 | 0.240302 |
| 0.349575 | 0.365646 | 0.081897 | 0.063783 | 0.041379 | 0.079485 |
| 0.187662 | 0.638168 | 0.082539 | 0.027239 | 0.022182 | 0.029948 |
| 0.318934 | 0.801348 | 0.547728 | 0.455099 | 0.211941 | 0.66026 |
| 0.07963 | 0.285886 | 0.52185 | 0.733209 | 0.814806 | 0.711812 |
| 0.942321 | 0.005013 | 0.009182 | 0.52588 | 0.037887 | 0.089454 |
| 0.156197 | 0.026555 | 0.022764 | 0.507146 | 0.059482 | 0.03224 |
| 0.186684 | 0.013812 | 0.012691 | 0.136369 | 0.297341 | 0.592892 |
| 0.254968 | 0.034784 | 0.220409 | 0.227338 | 0.221416 | 0.199278 |
| 0.698286 | 0.447916 | 0.262716 | 0.806355 | 0.174687 | 0.344472 |
| 0.942321 | 0.30489 | 0.033149 | 0.534254 | 0.169326 | 0.528259 |
| 0.167958 | 0.00981 | 0.038724 | 0.492756 | 0.488836 | 0.258278 |
| 0.937997 | 0.628511 | 0.215366 | 0.809448 | 0.047574 | 0.095014 |
| 0.452583 | 0.021513 | 0.542419 | 0.565848 | 0.042217 | 0.164498 |
| 0.606559 | 0.078831 | 0.526149 | 0.729068 | 0.335264 | 0.711812 |
| 0.138296 | 0.032115 | 0.251831 | 0.414391 | 0.081304 | 0.538177 |
| 0.251269 | 0.01836 | 0.437146 | 0.446997 | 0.23191 | 0.656353 |
| 0.505878 | 0.152651 | 0.654103 | 0.806289 | 0.062549 | 0.524509 |
| 0.086055 | 0.136046 | 0.775783 | 0.372175 | 0.406862 | 0.397402 |
| 0.34276 | 0.037759 | 0.32565 | 0.55694 | 0.335586 | 0.142428 |
| 0.740029 | 0.100535 | 0.047107 | 0.043941 | 0.270942 | 0.494834 |
| 0.092969 | 0.001588 | 0.020234 | 0.02875 | 0.045942 | 0.691288 |
| 0.038548 | 0.00211 | 0.002187 | 0.024801 | 0.052445 | 0.037925 |
| 0.106002 | 0.005693 | 0.52515 | 0.853513 | 0.052499 | 0.725062 |
| 0.59019 | 0.078567 | 0.816366 | 0.01605 | 0.13803 | 0.044576 |
| 0.223321 | 0.71933 | 0.084636 | 0.347641 | 0.016928 | 0.138008 |
| 0.138296 | 0.257733 | 0.261637 | 0.33546 | 0.288916 | 0.652502 |
| 0.23898 | 0.00858 | 0.192294 | 0.00258 | 0.690782 | 0.122586 |
| 0.061444 | 0.026434 | 0.002137 | 0.817518 | 0.177756 | 0.372101 |
| 0.452601 | 0.285521 | 0.675029 | 0.254599 | 0.178017 | 0.589818 |
| 0.080788 | 0.049293 | 0.584513 | 0.283281 | 0.287044 | 0.096731 |
| 0.716809 | 0.105795 | 0.387471 | 0.047623 | 0.232132 | 0.455424 |
| 0.076587 | 0.016167 | 0.023725 | 0.087749 | 0.079987 | 0.309069 |
| 0.228134 | 0.366708 | 0.177374 | 0.081751 | 0.301289 | 0.791898 |
| 0.167442 | 0.669424 | 0.09043 | 0.020431 | 0.272776 | 0.439662 |
| 0.153993 | 0.113568 | 0.134841 | 0.029229 | 0.150157 | 0.491761 |
| 0.039325 | 0.010505 | 0.045777 | 0.045086 | 0.496963 | 0.019762 |
| 0.085333 | 0.676007 | 0.298592 | 0.055044 | 0.406862 | 0.225419 |
| 0.57246 | 0.592268 | 0.09027 | 0.049984 | 0.245848 | 0.718651 |
| 0.251269 | 0.091671 | 0.038651 | 0.017711 | 0.087825 | 0.095326 |
| 0.368763 | 0.176406 | 0.645471 | 0.621117 | 0.744317 | 0.084564 |
| 0.227845 | 0.077032 | 0.034195 | 0.692 | 0.255504 | 0.343816 |
| 0.653968 | 0.046949 | 0.118782 | 0.493974 | 0.868687 | 0.095692 |
| 0.149838 | 0.006699 | 0.06762 | 0.181788 | 0.250101 | 0.286165 |
| 0.897026 | 0.750534 | 0.437146 | 0.642959 | 0.304423 | 0.661019 |
| 0.476088 | 0.037949 | 0.213912 | 0.198394 | 0.179696 | 0.117096 |
| 0.050019 | 0.02363 | 0.052825 | 0.076367 | 0.449428 | 0.767063 |
| 0.05064 | 0.011614 | 0.005644 | 0.057659 | 0.632673 | 0.407759 |
| 0.019579 | 0.383193 | 0.002807 | 0.083446 | 0.169956 | 0.566911 |

| | | | | | |
|---|---|---|---|---|---|
| 0.094812 | 0.016299 | 0.073499 | 0.073515 | 0.156732 | 0.126482 |
| 0.092969 | 0.024356 | 0.045404 | 0.531495 | 0.051947 | 0.101383 |
| 0.061444 | 0.023376 | 0.132432 | 0.096775 | 0.267746 | 0.091755 |
| 0.079136 | 0.008338 | 0.161933 | 0.026436 | 0.124242 | 0.178557 |
| 0.066202 | 0.326747 | 0.197183 | 0.035538 | 0.264092 | 0.538177 |
| 0.039325 | 0.065224 | 0.519911 | 0.032287 | 0.6495 | 0.10275 |
| 0.074536 | 0.026555 | 0.19158 | 0.257388 | 0.517424 | 0.113309 |
| 0.082861 | 0.103271 | 0.093021 | 0.31399 | 0.081391 | 0.123385 |
| 0.836449 | 0.070747 | 0.63156 | 0.13096 | 0.075968 | 0.092892 |
| 0.044587 | 0.021243 | 0.073828 | 0.016536 | 0.836747 | 0.094539 |
| 0.135096 | 0.026555 | 0.015515 | 0.051499 | 0.123254 | 0.054903 |
| 0.05064 | 0.044648 | 0.005401 | 0.621117 | 0.332243 | 0.111385 |
| 0.206898 | 0.019699 | 0.269487 | 0.081235 | 0.180999 | 0.083986 |
| 0.237019 | 0.019967 | 0.193443 | 0.318269 | 0.219793 | 0.100564 |
| 0.450746 | 0.54245 | 0.053401 | 0.047623 | 0.047574 | 0.036886 |
| 0.272369 | 0.415179 | 0.15568 | 0.181169 | 0.348797 | 0.072588 |
| 0.21336 | 0.015286 | 0.015728 | 0.113857 | 0.28541 | 0.146534 |
| 0.588742 | 0.037671 | 0.676176 | 0.522898 | 0.107401 | 0.063302 |
| 0.88104 | 0.092975 | 0.685516 | 0.101919 | 0.081391 | 0.054336 |
| 0.790172 | 0.009831 | 0.005401 | 0.801493 | 0.19759 | 0.029948 |
| 0.098759 | 0.036597 | 0.041416 | 0.05934 | 0.067438 | 0.070063 |
| 0.196242 | 0.156294 | 0.254811 | 0.518108 | 0.560957 | 0.063564 |
| 0.804383 | 0.033302 | 0.125157 | 0.12875 | 0.065892 | 0.065348 |
| 0.100645 | 0.070747 | 0.471955 | 0.083147 | 0.096203 | 0.026579 |
| 0.828001 | 0.655524 | 0.036653 | 0.522898 | 0.315948 | 0.630807 |
| 0.147098 | 0.16745 | 0.578296 | 0.138077 | 0.045942 | 0.019848 |
| 0.178985 | 0.006911 | 0.003637 | 0.036319 | 0.169326 | 0.101358 |
| 0.185123 | 0.264772 | 0.750436 | 0.162516 | 0.120592 | 0.027072 |
| 0.082861 | 0.034149 | 0.073575 | 0.130007 | 0.043366 | 0.05263 |
| 0.292508 | 0.47024 | 0.601702 | 0.263616 | 0.077176 | 0.023635 |
| 0.57246 | 0.70569 | 0.497813 | 0.566436 | 0.211941 | 0.019762 |
| 0.249785 | 0.512979 | 0.340165 | 0.299583 | 0.816203 | 0.024347 |
| 0.227845 | 0.037791 | 0.377377 | 0.200252 | 0.169404 | 0.073161 |
| 0.288612 | 0.146716 | 0.118782 | 0.424491 | 0.799387 | 0.872124 |
| 0.683129 | 0.509073 | 0.394645 | 0.68841 | 0.903711 | 0.8358 |
| 0.175313 | 0.013801 | 0.005401 | 0.025088 | 0.290513 | 0.159563 |
| 0.147098 | 0.127819 | 0.068031 | 0.029883 | 0.03711 | 0.039698 |
| 0.925515 | 0.434248 | 0.26385 | 0.545064 | 0.055938 | 0.15114 |
| 0.251269 | 0.004121 | 0.169957 | 0.007296 | 0.011481 | 0.018146 |
| 0.084565 | 0.00661 | 0.130531 | 0.051505 | 0.059482 | 0.072503 |
| 0.03921 | 0.00211 | 0.364269 | 0.06669 | 0.178017 | 0.33241 |
| 0.365004 | 0.363205 | 0.084636 | 0.124007 | 0.493749 | 0.475599 |
| 0.17967 | 0.064677 | 0.022483 | 0.529348 | 0.60067 | 0.319827 |
| 0.23898 | 0.56336 | 0.350212 | 0.26501 | 0.196146 | 0.045881 |
| 0.317879 | 0.284841 | 0.713108 | 0.51032 | 0.37876 | 0.666124 |
| 0.682362 | 0.188425 | 0.322701 | 0.414685 | 0.373483 | 0.150723 |
| 0.57246 | 0.058662 | 0.237504 | 0.458464 | 0.123293 | 0.101832 |
| 0.728619 | 0.251748 | 0.067113 | 0.391785 | 0.103072 | 0.136157 |
| 0.018824 | 0.005788 | 0.906332 | 0.020893 | 0.011481 | 0.03224 |
| 0.931516 | 0.163684 | 0.498897 | 0.096254 | 0.838274 | 0.869873 |
| 0.025757 | 0.114386 | 0.51119 | 0.017743 | 0.011481 | 0.06191 |
| 0.076385 | 0.04006 | 0.546148 | 0.113857 | 0.022796 | 0.022139 |
| 0.112915 | 0.165483 | 0.068331 | 0.156358 | 0.069995 | 0.020883 |
| 0.232211 | 0.014825 | 0.395537 | 0.323079 | 0.056064 | 0.061826 |
| 0.311883 | 0.024005 | 0.257278 | 0.806289 | 0.225064 | 0.019636 |
| 0.073189 | 0.024238 | 0.220203 | 0.268149 | 0.406862 | 0.040944 |
| 0.286315 | 0.008343 | 0.085003 | 0.012284 | 0.01791 | 0.00896 |
| 0.018824 | 0.003688 | 0.165433 | 0.029234 | 0.011481 | 0.019636 |
| 0.125692 | 0.02363 | 0.06354 | 0.149747 | 0.192924 | 0.131825 |
| 0.1181 | 0.062481 | 0.112205 | 0.091343 | 0.084291 | 0.157922 |
| 0.314886 | 0.169778 | 0.2863 | 0.159118 | 0.057202 | 0.111385 |
| 0.067344 | 0.609382 | 0.062906 | 0.026971 | 0.301289 | 0.086396 |

| | | | | | |
|---|---|---|---|---|---|
| 0.827653 | 0.54245 | 0.236464 | 0.630632 | 0.06568 | 0.1434 |
| 0.050019 | 0.347737 | 0.120584 | 0.855138 | 0.560188 | 0.753979 |
| 0.382999 | 0.827485 | 0.070212 | 0.309704 | 0.388696 | 0.457689 |
| 0.915283 | 0.87911 | 0.174744 | 0.655948 | 0.262189 | 0.439662 |
| 0.428797 | 0.285764 | 0.001346 | 0.294413 | 0.240393 | 0.065753 |
| 0.698286 | 0.387348 | 0.14746 | 0.330017 | 0.74784 | 0.726112 |
| 0.039547 | 0.004845 | 0.007403 | 0.016056 | 0.892196 | 0.308412 |
| 0.914633 | 0.466992 | 0.277796 | 0.31553 | 0.682937 | 0.568431 |
| 0.914633 | 0.466992 | 0.277796 | 0.31553 | 0.682937 | 0.568431 |
| 0.761821 | 0.025958 | 0.068031 | 0.047623 | 0.068522 | 0.06549 |
| 0.516484 | 0.055237 | 0.107184 | 0.33546 | 0.161187 | 0.434586 |
| 0.060143 | 0.023709 | 0.12277 | 0.209608 | 0.078169 | 0.100564 |
| 0.933351 | 0.027775 | 0.100569 | 0.111317 | 0.047574 | 0.096731 |
| 0.138796 | 0.730517 | 0.175751 | 0.072325 | 0.698657 | 0.270435 |
| 0.129003 | 0.053808 | 0.168374 | 0.737905 | 0.096703 | 0.507499 |
| 0.204919 | 0.124553 | 0.699428 | 0.293625 | 0.208168 | 0.213122 |
| 0.341064 | 0.880553 | 0.053044 | 0.521134 | 0.325249 | 0.066062 |
| 0.080788 | 0.118554 | 0.313227 | 0.516367 | 0.101329 | 0.095801 |
| 0.125812 | 0.843743 | 0.384194 | 0.35654 | 0.06632 | 0.037598 |
| 0.061444 | 0.04006 | 0.091679 | 0.446055 | 0.19118 | 0.117096 |
| 0.175313 | 0.176178 | 0.107553 | 0.050503 | 0.377214 | 0.063294 |
| 0.227845 | 0.176965 | 0.34142 | 0.036924 | 0.449428 | 0.144049 |
| 0.57246 | 0.061939 | 0.174744 | 0.393963 | 0.406862 | 0.068284 |
| 0.761821 | 0.637628 | 0.123236 | 0.523887 | 0.359898 | 0.190783 |
| 0.146004 | 0.847491 | 0.276669 | 0.039346 | 0.09558 | 0.449878 |
| 0.57246 | 0.666844 | 0.202554 | 0.090136 | 0.011481 | 0.039698 |
| 0.57246 | 0.666844 | 0.202554 | 0.090136 | 0.011481 | 0.039698 |
| 0.648737 | 0.669424 | 0.141011 | 0.174805 | 0.251553 | 0.075981 |
| 0.027399 | 0.092975 | 0.072895 | 0.012941 | 0.023646 | 0.029948 |
| 0.717762 | 0.098875 | 0.091075 | 0.083147 | 0.047574 | 0.579767 |
| 0.408784 | 0.074109 | 0.811925 | 0.811544 | 0.229153 | 0.029948 |
| 0.574643 | 0.286416 | 0.670734 | 0.736112 | 0.037887 | 0.089585 |
| 0.244482 | 0.107332 | 0.316803 | 0.081751 | 0.011481 | 0.029948 |
| 0.375884 | 0.765237 | 0.084636 | 0.555308 | 0.034819 | 0.054644 |
| 0.355895 | 0.254864 | 0.110013 | 0.782926 | 0.037887 | 0.096731 |
| 0.241638 | 0.015349 | 0.442378 | 0.111317 | 0.025334 | 0.080842 |
| 0.210475 | 0.02289 | 0.639307 | 0.581918 | 0.052829 | 0.038918 |
| 0.313508 | 0.040405 | 0.693804 | 0.261222 | 0.020857 | 0.087613 |
| 0.912596 | 0.019967 | 0.304456 | 0.454911 | 0.028466 | 0.072503 |
| 0.530436 | 0.023759 | 0.213248 | 0.249557 | 0.03743 | 0.308235 |
| 0.086055 | 0.018227 | 0.156832 | 0.477142 | 0.045539 | 0.038918 |
| 0.018824 | 0.843743 | 0.031259 | 0.016056 | 0.010345 | 0.012009 |
| 0.168985 | 0.177384 | 0.328411 | 0.272329 | 0.035784 | 0.036886 |
| 0.21336 | 0.387179 | 0.84024 | 0.642 | 0.046107 | 0.159085 |
| 0.0419 | 0.008285 | 0.696319 | 0.030112 | 0.037574 | 0.383822 |
| 0.087027 | 0.019793 | 0.397292 | 0.026727 | 0.029392 | 0.185104 |
| 0.349575 | 0.03529 | 0.369926 | 0.174805 | 0.099177 | 0.509309 |
| 0.3522 | 0.021513 | 0.674898 | 0.57412 | 0.048657 | 0.194447 |
| 0.126786 | 0.051797 | 0.137568 | 0.475634 | 0.019763 | 0.048441 |
| 0.153993 | 0.034915 | 0.72849 | 0.849682 | 0.051402 | 0.02908 |
| 0.289983 | 0.02145 | 0.910341 | 0.113517 | 0.118376 | 0.787288 |
| 0.025381 | 0.003694 | 0.084971 | 0.012941 | 0.010345 | 0.015024 |
| 0.013729 | 0.039027 | 0.375844 | 0.020431 | 0.011481 | 0.037925 |
| 0.018824 | 0.009971 | 0.145328 | 0.014091 | 0.011481 | 0.028678 |
| 0.019355 | 0.005829 | 0.06674 | 0.048212 | 0.045942 | 0.029948 |
| 0.024815 | 0.003312 | 0.054298 | 0.047583 | 0.015085 | 0.027415 |
| 0.251269 | 0.078876 | 0.002187 | 0.120046 | 0.048657 | 0.101358 |
| 0.043144 | 0.086636 | 0.142272 | 0.548853 | 0.127632 | 0.101463 |
| 0.086055 | 0.018458 | 0.071208 | 0.659698 | 0.075841 | 0.067079 |
| 0.018841 | 0.793434 | 0.00654 | 0.037975 | 0.740299 | 0.790176 |
| 0.167896 | 0.095857 | 0.014039 | 0.828105 | 0.070758 | 0.014194 |
| 0.413321 | 0.544887 | 0.560562 | 0.321566 | 0.196752 | 0.368773 |

| | | | | | |
|---|---|---|---|---|---|
| 0.152539 | 0.165483 | 0.151308 | 0.815336 | 0.216206 | 0.461431 |
| 0.847625 | 0.035778 | 0.779604 | 0.782926 | 0.047574 | 0.098699 |
| 0.074189 | 0.004845 | 0.68567 | 0.02217 | 0.056816 | 0.028159 |
| 0.389271 | 0.388258 | 0.431982 | 0.212275 | 0.004882 | 0.118446 |
| 0.068843 | 0.006396 | 0.010472 | 0.025967 | 0.278346 | 0.201946 |
| 0.128619 | 0.010262 | 0.062906 | 0.06733 | 0.037887 | 0.099378 |
| 0.353297 | 0.004845 | 0.007334 | 0.126187 | 0.052499 | 0.45758 |
| 0.950125 | 0.022873 | 0.002187 | 0.050285 | 0.04247 | 0.155339 |
| 0.233384 | 0.004845 | 0.000379 | 0.020431 | 0.118376 | 0.87006 |
| 0.425888 | 0.000892 | 0.161933 | 0.072325 | 0.022985 | 0.2872 |
| 0.31433 | 0.024356 | 0.142272 | 0.424491 | 0.451459 | 0.280445 |
| 0.092534 | 0.018257 | 0.016472 | 0.020431 | 0.252247 | 0.19149 |
| 0.053998 | 0.034526 | 0.006725 | 0.025967 | 0.031615 | 0.024982 |
| 0.108587 | 0.012947 | 0.628282 | 0.432556 | 0.00787 | 0.058384 |
| 0.125812 | 0.03722 | 0.026199 | 0.68841 | 0.069995 | 0.336449 |
| 0.292508 | 0.206841 | 0.630034 | 0.414673 | 0.392123 | 0.2076 |
| 0.567105 | 0.033089 | 0.024237 | 0.072091 | 0.057794 | 0.087439 |
| 0.189264 | 0.074382 | 0.002776 | 0.164116 | 0.006313 | 0.155815 |
| 0.040192 | 0.016287 | 0.025239 | 0.074275 | 0.149423 | 0.039302 |
| 0.580759 | 0.255554 | 0.84024 | 0.522884 | 0.903927 | 0.833937 |
| 0.874707 | 0.53588 | 0.347447 | 0.566436 | 0.311602 | 0.280594 |
| 0.627296 | 0.065093 | 0.17991 | 0.159118 | 0.004882 | 0.076763 |
| 0.031039 | 0.006656 | 0.010373 | 0.174805 | 0.011481 | 0.021682 |
| 0.147098 | 0.713908 | 0.602236 | 0.083147 | 0.393326 | 0.566485 |
| 0.025808 | 0.02363 | 0.002748 | 0.016056 | 0.011824 | 0.029948 |
| 0.018824 | 0.847741 | 0.295021 | 0.015679 | 0.139647 | 0.507499 |
| 0.390506 | 0.098596 | 0.001443 | 0.126005 | 0.518458 | 0.686976 |
| 0.570668 | 0.305926 | 0.071281 | 0.658332 | 0.820839 | 0.258278 |
| 0.149862 | 0.037174 | 0.011243 | 0.242787 | 0.066204 | 0.015713 |
| 0.343063 | 0.011121 | 0.497973 | 0.80373 | 0.080691 | 0.135841 |
| 0.405275 | 0.276237 | 0.192112 | 0.52588 | 0.343952 | 0.036886 |
| 0.361286 | 0.427823 | 0.104801 | 0.096628 | 0.363461 | 0.220644 |
| 0.441111 | 0.015349 | 0.043084 | 0.479936 | 0.038072 | 0.074892 |
| 0.323467 | 0.034915 | 0.043084 | 0.190569 | 0.348797 | 0.397402 |
| 0.749265 | 0.354518 | 0.857531 | 0.374382 | 0.287044 | 0.460674 |
| 0.31387 | 0.277219 | 0.058013 | 0.407902 | 0.903927 | 0.719214 |
| 0.772174 | 0.193115 | 0.499534 | 0.765685 | 0.331139 | 0.562937 |
| 0.815955 | 0.358763 | 0.086458 | 0.826443 | 0.047574 | 0.115824 |
| 0.370182 | 0.043135 | 0.00581 | 0.025967 | 0.08036 | 0.139493 |
| 0.370182 | 0.389958 | 0.012965 | 0.367762 | 0.022182 | 0.202187 |
| 0.794392 | 0.025387 | 0.018324 | 0.076709 | 0.074826 | 0.149846 |
| 0.107978 | 0.045175 | 0.031259 | 0.447427 | 0.045539 | 0.119326 |
| 0.698286 | 0.153769 | 0.008867 | 0.058615 | 0.523379 | 0.690208 |
| 0.092969 | 0.041352 | 0.325724 | 0.047623 | 0.013645 | 0.012009 |
| 0.191448 | 0.027499 | 0.463132 | 0.527977 | 0.082814 | 0.019762 |
| 0.503374 | 0.00478 | 0.863506 | 0.439593 | 0.088226 | 0.023408 |
| 0.031039 | 0.001085 | 0.002753 | 0.051499 | 0.041825 | 0.041366 |
| 0.068843 | 0.054638 | 0.00009 | 0.02875 | 0.162461 | 0.178557 |
| 0.125692 | 0.227554 | 0.287111 | 0.133312 | 0.211941 | 0.045898 |
| 0.147098 | 0.00858 | 0.033149 | 0.332953 | 0.801879 | 0.101358 |
| 0.146129 | 0.006911 | 0.526238 | 0.131665 | 0.185761 | 0.24899 |
| 0.118967 | 0.065944 | 0.120059 | 0.493566 | 0.894366 | 0.19149 |
| 0.071155 | 0.114653 | 0.000379 | 0.117253 | 0.289331 | 0.243379 |
| 0.127957 | 0.026853 | 0.101155 | 0.736449 | 0.205682 | 0.117096 |
| 0.036622 | 0.030327 | 0.013344 | 0.271128 | 0.345694 | 0.579767 |
| 0.090777 | 0.039748 | 0.648037 | 0.209565 | 0.678899 | 0.548089 |
| 0.018824 | 0.061176 | 0.01303 | 0.435286 | 0.29513 | 0.397402 |
| 0.635417 | 0.158005 | 0.062085 | 0.248405 | 0.418358 | 0.573692 |
| 0.251269 | 0.001085 | 0.019949 | 0.020431 | 0.019624 | 0.029948 |
| 0.036559 | 0.11789 | 0.049849 | 0.454911 | 0.398923 | 0.242544 |
| 0.093296 | 0.008995 | 0.22954 | 0.566092 | 0.392123 | 0.315168 |
| 0.031039 | 0.443135 | 0.000424 | 0.454992 | 0.489593 | 0.8358 |

| | | | | | |
|---|---|---|---|---|---|
| 0.128619 | 0.338036 | 0.016721 | 0.587498 | 0.399625 | 0.439074 |
| 0.084565 | 0.004845 | 0.010509 | 0.062902 | 0.448401 | 0.267488 |
| 0.508147 | 0.01397 | 0.001346 | 0.477142 | 0.518458 | 0.436924 |
| 0.313508 | 0.383004 | 0.150394 | 0.209565 | 0.449428 | 0.661019 |
| 0.031039 | 0.049293 | 0.085865 | 0.126005 | 0.406862 | 0.351703 |
| 0.031039 | 0.011169 | 0.34131 | 0.109588 | 0.36977 | 0.769347 |
| 0.065462 | 0.012104 | 0.07546 | 0.458464 | 0.449428 | 0.354879 |
| 0.053408 | 0.026853 | 0.306479 | 0.269034 | 0.160262 | 0.086396 |
| 0.13155 | 0.138101 | 0.328411 | 0.101165 | 0.444974 | 0.2872 |
| 0.247999 | 0.051687 | 0.096914 | 0.041678 | 0.479869 | 0.687826 |
| 0.578826 | 0.021745 | 0.062276 | 0.038612 | 0.883871 | 0.159563 |
| 0.018824 | 0.00981 | 0.387471 | 0.204193 | 0.07041 | 0.198853 |
| 0.305742 | 0.290758 | 0.045034 | 0.117971 | 0.101858 | 0.137308 |
| 0.744272 | 0.202953 | 0.063434 | 0.053629 | 0.270402 | 0.438914 |
| 0.26978 | 0.72362 | 0.389178 | 0.477739 | 0.228643 | 0.465995 |
| 0.153993 | 0.008608 | 0.037822 | 0.034785 | 0.029392 | 0.059239 |
| 0.194563 | 0.439731 | 0.750135 | 0.105464 | 0.118376 | 0.47928 |
| 0.125812 | 0.020449 | 0.163161 | 0.063919 | 0.069995 | 0.026579 |
| 0.58515 | 0.207683 | 0.033075 | 0.72187 | 0.657419 | 0.037925 |
| 0.174474 | 0.043098 | 0.902345 | 0.338453 | 0.162039 | 0.559799 |
| 0.363448 | 0.076954 | 0.098373 | 0.020431 | 0.022182 | 0.065753 |
| 0.90437 | 0.362791 | 0.526238 | 0.101535 | 0.8865 | 0.11236 |
| 0.174773 | 0.09172 | 0.630034 | 0.806289 | 0.777344 | 0.097965 |
| 0.282863 | 0.052801 | 0.114552 | 0.030112 | 0.154997 | 0.115824 |
| 0.349575 | 0.116896 | 0.033905 | 0.550291 | 0.075917 | 0.029445 |
| 0.135637 | 0.051713 | 0.06084 | 0.045049 | 0.087407 | 0.03224 |
| 0.915283 | 0.031939 | 0.39983 | 0.220556 | 0.313554 | 0.343816 |
| 0.358789 | 0.322983 | 0.559074 | 0.334149 | 0.038072 | 0.288417 |
| 0.739825 | 0.029496 | 0.130398 | 0.328183 | 0.138575 | 0.202703 |
| 0.502564 | 0.023434 | 0.909757 | 0.038088 | 0.029392 | 0.126482 |
| 0.685123 | 0.036681 | 0.155016 | 0.058513 | 0.022896 | 0.106072 |
| 0.784208 | 0.028576 | 0.26683 | 0.36077 | 0.108976 | 0.203442 |
| 0.57246 | 0.092142 | 0.202554 | 0.063783 | 0.028466 | 0.118446 |
| 0.594342 | 0.096023 | 0.288397 | 0.392734 | 0.13575 | 0.211909 |
| 0.543211 | 0.062505 | 0.09043 | 0.424491 | 0.00787 | 0.040699 |
| 0.018824 | 0.011987 | 0.060196 | 0.505828 | 0.311602 | 0.019762 |
| 0.176418 | 0.080849 | 0.451885 | 0.397899 | 0.184483 | 0.416513 |
| 0.915283 | 0.287412 | 0.244966 | 0.316531 | 0.150552 | 0.346824 |
| 0.752956 | 0.611403 | 0.63156 | 0.587666 | 0.458958 | 0.566911 |
| 0.784208 | 0.519632 | 0.414152 | 0.364696 | 0.392123 | 0.693784 |
| 0.813884 | 0.475344 | 0.578453 | 0.213361 | 0.45662 | 0.2566 |
| 0.748619 | 0.254953 | 0.671883 | 0.250406 | 0.290513 | 0.256234 |
| 0.089303 | 0.609382 | 0.001346 | 0.613828 | 0.777344 | 0.510249 |

| | | | |
|---|---|---|---|
| dUTP-nega | C9H15N2O14P3 | C00460 | 19110.41365 |
| ethanolamine | C2H7NO | C00189 | 42051.16202 |
| FAD | C27H33N9O15P2 | C00016 | 485926.3981 |
| Flavone | C15H10O2 | C15608 | 6927.795207 |
| FMN | C17H21N4O9P | C00061 | 88273.55147 |
| folate | C19H19N7O6 | C00504 | 3212.120299 |
| fructose-1,6-bisphosphate | C6H14O12P2 | C05378 | 6881612.067 |
| fructose-6-phosphate | C6H13O9P | C05345 | 531280.3081 |
| fumarate | C4H4O4 | C00122 | 5709756.796 |
| GDP | C10H15N5O11P2 | C00035 | 1322901.876 |
| Geranyl-PP | C10H20O7P2 | C00341 | 50468.60851 |
| glucono-$\delta$-lactone | C6H10O6 | C00198 | 46574.33789 |
| glucosamine | C6H13NO5 | C00329 | 12983.23712 |
| glucose-1-phosphate | C6H13O9P | C00103 | 8316089.332 |
| glucose-6-phosphate | C6H13O9P | C00668 | 438252.1341 |
| glutamate | C5H9NO4 | C00025 | 63882695.31 |
| glutamine | C5H10N2O3 | C00064 | 2462653.506 |
| glutathione | C10H17N3O6S | C00051 | 282891.806 |
| glutathione disulfide | C20H32N6O12S2 | C00127 | 4282955.939 |
| glutathione | C10H17N3O6S | C00051 | 235634.6139 |
| glycerate | C3H6O4 | C00258 | 80660.53434 |
| glycerol 3-phosphate | C3H9O6P | C00093 | 27946784.24 |
| Glycerophosphocholine | C8H21NO6P | C00670 | 109003689 |
| glycine | C2H5NO2 | C00037 | 2093.545671 |
| glycolate | C2H4O3 | C00160 | 51338.22606 |
| glyoxylate | C2H2O3 | C00048 | 105659.8 |
| GMP | C10H14N5O8P | C00144 | 13074585.04 |
| GTP | C10H16N5O13P3 | C00044 | 794603.3295 |
| Guanidoacetic acid | C3H7N3O2 | C00581 | 29123588.57 |
| guanine | C5H5N5O | C00242 | 939512.642 |
| guanosine | C10H13N5O5 | C00387 | 328084.5399 |
| guanosine 5-diphosphate,3-diphosphate | C10H11N5O17P4 | C01228 | 178070.1238 |
| hexose-phosphate | C6H13O9P | C05345 | 21281860.91 |
| histidine | C6H9N3O2 | C00135 | 5568474.769 |
| histidinol | C6H11N3O | C00860 | 5827.563993 |
| HMG-CoA | C27H44N7O20P3S | C00356 | 4282.976548 |
| homocysteic acid | C4H9NO5S | C16511 | 162658.1334 |
| homocysteine | C4H9NO2S | C00155 | 48913.29464 |
| homoserine | C4H9NO3 | C00263 | 54802.78067 |
| Hydroxyisocaproic acid | C6H12O3 | HMDB00746 | 80879746.11 |
| Hydroxyphenylacetic acid | C8H8O3 | C05852 | 357424.605 |
| hydroxyphenylpyruvate | C9H8O4 | C01179 | 10510.30656 |
| hydroxyproline | C5H9NO3 | C01157 | 37455034.52 |
| hypoxanthine | C5H4N4O | C00262 | 52236990.7 |
| IDP | C10H14N4O11P2 | C00104 | 3247854.412 |
| Imidazoleacetic acid | C6H8N2O2 | C02835 | 209711.8948 |
| IMP | C10H13N4O8P | C00130 | 10333159.72 |
| indole | C8H7N | C00463 | 937845.5738 |
| Indole-3-carboxylic acid | C9H7NO2 | HMDB03320 | 2087939.389 |
| Indoleacrylic acid | C11H9NO2 | HMDB00734 | 784911.4213 |
| inosine | C10H12N4O5 | C00294 | 7587643.377 |
| isocitrate | C6H8O7 | C00311 | 228510.8815 |
| itaconic acid | C5H6O4 | HMDB0002092 | 808939.3585 |
| Kynurenic acid | C10H7NO3 | C01717 | 2755836.128 |
| Kynurenine | C10H12N2O3 | C00328 | 265687.3322 |
| lactate | C3H6O3 | C00186 | 69302660.97 |
| L-arginino-succinate | C10H18N4O6 | C03406 | 170787.9818 |
| leucine-isoleucine | C6H13NO2 | C00123 | 227432731.3 |
| lipoate | C8H14O2S2 | C00725 | 7546.6044 |
| lysine | C6H14N2O2 | C00047 | 41096759.06 |
| malate | C4H6O5 | C00149 | 34076504.61 |
| Maleic acid | C4H4O4 | C01384 | 5197549.859 |
